# Competition and growth in *Aedes aegypti* larvae: effects of distributing food inputs over time

**DOI:** 10.1101/2020.06.02.129478

**Authors:** Kurt Steinwascher

**Affiliations:** Formerly of the Florida Medical Entomology Laboratory, 200 9^th^ St SE, Vero Beach, FL 32962 United States of America

**Author notes:** [retired] 445 Kailua Rd 5209 Kailua HI 96734 United States of America.

**Keywords:** *Aedes aegypti*, age at pupation, ANOVA, competition, egg size, exploitative, feeding behavior, female advantage, filter feeding, food x density, growth rate, intra specific, interference, inter specific, larval density, larval development, life history, MANOVA, maximum size, minimum size, multiple food inputs, multivariate analysis of variance, oviposition site selection, propagule size, provisioning, pupal mass, pupation, pupation triggers, resource partitioning, sex differences, size distribution, starvation, survival, timing of food inputs

## Abstract

Male and female mosquito larvae compete for different subsets of the yeast food resource in laboratory microcosms. Males compete more intensely with males and females with females. The amount and timing of food inputs alters both growth and competition, but the effects are different between sexes. Increased density increases competition among males. Among females, density operates primarily by changing the food/larva or total food; this affects competition in some interactions and growth in others. Food added earlier in the life span contributes more to mass than the same quantity added later. After a period of starvation larvae appear to use some of the subsequent food input to rebuild physiological reserves in addition to building mass. The timing of pupation is affected by the independent factors and competition, but not in the same way for the two sexes, and not in the same way as mass at pupation for the two sexes. There is an effect of density on the timing of pupation for females independent of competition or changes in food/larva or total food. Male and female larvae have different larval life history strategies. Males grow quickly to a minimum size, then pupate, depending on the amount of food available. Males that do not grow quickly enough may delay pupation further to grow larger, resulting in a bimodal distribution of sizes and ages. Males appear to have a maximum size determined by the early food level. Females grow faster than males and grow larger than males on the same food inputs. Females affect the growth and competition among males by manipulating the number of particles in the microcosm through changes in feeding behavior. Mosquito larvae appear to have evolved to survive periods of starvation and take advantage of intermittent inputs of food into containers.

## Introduction

Competition may organize communities in nature, but this is difficult to demonstrate unambiguously. Competition for food appears to affect the outcomes of laboratory microcosms [1–21]. For *Aedes aegypti* larvae in these microcosms, the food level and the density of larva both affect intraspecific competition, which shows up as the interaction between food and density. Furthermore, the total food affects the nature of competition among the larvae, also apparent in the interaction of food and density. Density, total food, and food per larva are interdependent, but all three factors affect competition among mosquito larvae (see [1] and literature review therein). Male and female larvae compete differently from one another for yeast cells; female larvae outcompete males through larger size and by retaining cells within their gut at low total food levels. As competition intensifies, the pupal masses of both males and females decrease, so the effect of competition is a reduced apparent food level. The age at pupation is also affected by food and density. Male and female larvae delay pupation to grow larger when food is limiting and also when food is abundant, but the cues that result in these delays differ between the sexes [1].

This report covers three experiments that investigate competition and growth of mosquito larvae in response to the distribution of food inputs during the larval life span. The first experiment covers competition among larvae, but necessarily also covers the growth of larvae. The second experiment looks at the difference between larvae grown in isolation and larvae raised with one or two competitors. Differential survival of the single larva obstructed the analysis of this data, but the survivorship results informed the design of the third experiment along with the first experiment. The third experiment extends the results of the first experiment by examining the growth of single larvae and the effect of starvation and late food additions on pupation.

The first experiment tests the hypothesis that the distribution of food inputs over time will affect intraspecific competition for food among mosquito larvae as has been shown for tadpoles [9] [22]. In nature, rainfall washes additional food into the tires and other containers that the larvae inhabit, so this is a reasonable test. Other investigators have found that food inputs affect the outcome of mosquito larval growth in microcosms [5] [7] [23–28]. Here, food and density treatments were selected to cover the span of density and food determined in a prior experiment [1] to produce significant differences in the nature of non-lethal competition among *A. aegypti* larvae. The food and density treatments should produce an interaction that will feature microcosms with the most competition, least competition and two with intermediate levels of competition. The other two factors vary the amount of food being added and the timespan over which the food is added to each microcosm. If either or both of these factors affect competition among the larvae, there should be significant 3-way or 4-way interactions among food, density, and the other two factors. The total food levels and relative food levels are high enough that interference mechanisms of competition are unlikely [1] [8] [29–33]. Individual larvae may compete exploitatively by adjusting consumption rate or the efficiency of processing food particles. Previous results suggest that males maximize consumption rate while females vary efficiency to maximize growth [1].

In addition to the hypotheses about competition, the first experiment also tests hypotheses about the growth of larvae independent of density, and about potential competition for food independent of the total food level.

The second experiment tests the hypothesis that isolated larvae grow and pupate similarly to larvae at very low densities.

The third experiment extends the investigation into the growth of mosquito larvae after a period of starvation similar to the longer delay in the first experiment. It looks at the value of late food additions and the effect of the timing and amount of those additions on the growth of the single larva. Data from the first experiment indicate that competition and differential growth results in qualitatively and quantitatively different outcomes in the treatments with longest intervals between food inputs. The third experiment tests hypotheses about the equivalence of food inputs after a period of starvation similar to those in the first experiment. There is only a single larva in each test tube, so this experiment investigates growth rather than competition.

## Methods

### First experiment

Eggs were obtained from fifth generation female *A. aegypti* in a colony derived from collections in tires near the Dade County Public Works Department (Florida). Eggs were hatched in distilled water and larvae were counted into test tubes (new, disposable, round bottomed, approximately 25 x 150 mm) containing 20 ml distilled water. Two density treatments, 4 or 8 larvae/test tube, were crossed with two total food levels treatments, 16 or 32 mg baker’s yeast/test tube. Two further factors were added to investigate the distribution of food input over time. For the aliquot factor, the total food was divided into 2 or 4 aliquots. The first aliquot was always presented on the day of hatching (day 0). The subsequent aliquots were distributed across a timespan of either 3 or 6 days. Each additional input was suspended in 0.5 ml distilled water. The amount and distribution of aliquots across the 16 treatments of this 4-factor experiment are listed in S1 Table. Ten biological replicates of each treatment combination were initiated. 960 1st instar larvae were counted into 160 test tubes according to the 16 treatments. Each test tube was then provided with the appropriate amount of suspended yeast cells for that treatment. The test tubes were stored in randomized sequence and kept at ambient conditions throughout the experiment. Additional food was provided according to treatment on the appropriate days. The test tubes were examined for pupae from day 4 until day 35 when the last larva died. Pupae were removed, identified to sex, and weighed to the nearest 0.01 mg. On day 6, pupae were removed before the final aliquots of food were added. [The detailed, step-by-step methodology is presented in dx.doi.org/10.17504/protocols.io.bddhi236].

This is a 2×2×2×2 experimental design. The experiment crosses two additional factors, aliquot and timespan, with the four competition treatments to look at how the rate and timing of food inputs affect larval competition among females, among males and between the sexes. There are four main effects, six 2-way interactions, four 3-way interactions and one 4-way interaction. Some of the interactions describe competition, others describe growth, and others could describe competition or growth. Results from the prior experiment [1] suggest the outcome of competitive interactions. Females dominate males in larval competition. Females should be largest and pupate earliest in the treatments with the least competition and smallest and pupate latest in the treatments with the most competition. The two intermediate competition treatments offer the same food per larva at different densities (so the total food will be higher at the higher density). Females in the intermediate competition treatments should be larger at the higher density than at the lower density because of the higher total food despite the same food per larva in both treatments. Males respond differently to the competition treatments and are also affected by competition with females. Males and females may extend their larval growth in response to competition and to take advantage of food abundance.

In addition to the interactions involving competition (food and density), there are also interactions involving only aspects of the food supply (food, aliquot and timespan), and interactions between density, aliquot and timespan. The interactions involving only food, aliquot and timespan describe the effects of these attributes of the food supply on growth, not competition. The interactions involving density, aliquot and timespan may describe competitive effects that are independent of the total food level, and due only to the distribution of the food inputs in time. Alternatively, these interactions may describe the effects of these factors on growth, not competition.

At the conclusion of the experiment, 7 dependent variables were calculated for each replicate: Survival, Prime male mass and age at pupation, Average male mass, Prime female mass and age at pupation, and Average female mass. The Prime male was the male with the greatest growth rate; this is the first male to pupate, or the largest of the first males to pupate. The Prime female was the largest female to pupate. The distinction between the two Prime individuals reflects differences in the growth and pupation of the two sexes [1] [34–36]. Test tubes with survivors of only one sex were excluded from analysis. The data were analyzed as a multivariate, four-way analysis of variance (MANOVA) [37, 38].

### Second experiment

Eggs were obtained and hatched as previously described, to produce 120 1^st^ instar larvae. 60 flat-bottomed shell vials (25 mm diameter x 95 mm tall) were numbered and filled with 20 ml distilled water containing a concentration of baker’s yeast to produce the food level treatments (2 mg, 3 mg, 4 mg, and 5 mg dry weight of yeast per larva). 1, 2 or 3 larvae were added to each vial according to randomized treatment. 5 replicates of each treatment were initiated. S2 Table lists the treatments and replicates. The test tubes were examined for pupae from day 4 until the last larva died. Pupae were removed, identified to sex, and weighed to the nearest 0.01 mg. [The detailed, step-by-step methodology is presented in dx.doi.org/10.17504/protocols.io.bddhi236].

### Third experiment

Eggs were obtained and hatched as previously described, to produce 150 1^st^ instar larvae. 150 test tubes (new, disposable, round bottomed, approximately 25 x 150 mm) were numbered, then assigned to a treatment using a random number table. There were 6 treatments with 25 physical replicates each. S3 Table lists the treatments with the incremental food and delay. [The detailed, step-by-step methodology is presented in dx.doi.org/10.17504/protocols.io.bddhi236].

Each test tube received 20 ml distilled water with 1 mg Baker’s yeast and a single 1^st^ instar larva on day 0. Test tubes were examined for pupa beginning on day 4, the day before any pupa were expected. On day 6, after examining for dead larvae and pupae, treatments 1-3 received 1 mg, 2 mg, or 3 mg additional yeast (dry weight) respectively. On day 7, test tubes were examined for dead larvae and pupae. On day 8, after examining for dead larvae and pupae, treatments 4-6 received 1 mg, 2 mg, or 3 mg additional yeast (dry weight) respectively. On subsequent days, test tubes were examined for dead larvae and pupae.

Each pupa was blotted on paper toweling to remove excess water and weighed to the nearest 0.01 mg. The weight (mg), sex (M/F), and age at pupation (days) were recorded for each tube with a pupa. Larvae that died before pupating are also recorded by day at which death is observed.

The mosquito larvae are not expected to be able to pupate on the initial amount of food (1 mg), but they should be able to survive until the second food input. The amount of food at the second food input (1 mg, 2 mg or 3 mg) should allow pupation and the size and age at pupation should improve with increasing amounts. The shorter delay should be better for both size and age than the longer delay. Better outcomes mean larger masses and earlier ages at pupation. Males and females may delay pupation in response to abundant food, so there are possible interactions between the mass and age variables, which should show up in the MANOVA. There is no competition in this experiment because larvae are alone in their test tubes, so sex is treated as a factor rather than as covarying dependent variables.

The data are analyzed as a multivariate, three-way analysis of variance (MANOVA) [37, 38] with 3 factors: amount of incremental food (1 mg, 2 mg, or 3 mg); delay (6 or 8 days between the start of the experiment and the input of the incremental food); and sex (male or female). The two dependent variables are: mass at pupation (mg) and days to pupation after the second input of food. The age at pupation is calculated as: day of pupation MINUS the delay, 6 days or 8 days, depending on the treatment. The transformation of the age by subtracting the delay removes the numerical effect of the delay from the biological effect of the delay.

This research was conducted according to the standard guidelines at the time (1979-1982), sanctioned by the NIH, and under the supervision of the appropriate personnel at the Florida Medical Entomology Laboratory (IFAS and the University of Florida at Gainesville). Drs. J. Howard Frank and L. P. Lounibos specifically approved this study. The protocols are detailed at: dx.doi.org/10.17504/protocols.io.bddhi236

## Results

### First experiment

776 of the 960 1st instars pupated (81% survival). 348 females and 428 males pupated (55% males). Females ranged in size from 1.41 mg to 5.69 mg (average 3.55 mg, standard deviation 0.89 mg) and males ranged from 0.94 mg to 4.73 mg (average 2.26 mg, standard deviation 0.46 mg). Females pupated from day 5 through day 17 (average 6.69 days, standard deviation 2.07 days) and males pupated from day 5 through day 13 (average 5.70 days, standard deviation 1.47 days). About 1/3 of the females (106) pupated on day 5 and a second 1/3 (118) pupated on day 6. 39 females pupated after day 6 in the test tubes with the 3 day timespan and 85 females pupated after day 6 in the test tubes with the 6 day timespan (additional food was added at the end of the 6th day in these test tubes). More than half of the males (296) pupated on day 5 and another 75 pupated on day 6. 11 males pupated after day 6 in the test tubes with the 3 day timespan and 46 males pupated after day 6 in the test tubes with the 6 day timespan (additional food was added at the end of the 6th day in these test tubes). Histograms of age and mass at pupation for males and females are presented in S1 Fig-S4 Fig. The raw data is presented in S1 Dataset.

There are seven dependent variables: Survival, Prime male mass and age at pupation, Average male mass at pupation, Prime female mass and age at pupation, and Average female mass at pupation. Each of these variables is potentially affected by each of the main effects and interactions in the MANOVA (and the ANOVAs). There are also biologically meaningful relationships among the dependent variables: the relationship between Survival and each of the other six; between Prime male mass and age; between Prime female mass and age; between Prime male mass and Average male mass; between Prime female mass and Average female mass; between Prime female mass and Prime male mass; and between Average female mass and Average male mass. A strong relationship between Survival and any of the other six variables would be undesirable. The Prime mass and age variables potentially vary independently of each other, although previous experiments suggest a negative correlation between mass and age at pupation: large masses are associated with early ages and small masses with later ages at pupation. The estimated growth rate (mean mass divided by mean age) describes the joint response of these two variables to the treatments for Prime males and females. High estimated growth rates are indicative of good conditions for the larvae (low competition and abundant food). The Prime mass is included in the Average, so the difference between Prime mass and Average mass indicates the size distribution of the larvae, and the relative advantage of the Prime over the non-Prime individuals across the treatments. In the case where the Average mass is affected by a treatment, but the Prime is not, the affected individuals are the non-Prime larvae in the test tubes. In the case where the Prime mass is affected, but the Average is not, the effect of the treatment on the non-Prime individuals is collectively equal and opposite to the effect on the Prime. If the Prime individual gets larger, but the Average stays the same, then the non-Prime individuals must have suffered a reduction in size to offset the larger Prime. The difference between the Prime female mass and the Prime male mass and between the estimated growth rates for the Prime males and females, both indicate the relative advantage of the females over the males. The two Prime individuals represent the competitive winners in each microcosm. The difference between the Average female mass and the Average male mass also indicates the relative advantage of the females over the males, but reflects the non-Prime individuals as well. All of these relationships must be considered for each of the interactions to interpret the biological meaning of the statistical results.

### MANOVA

All four of the main treatments have a significant effect in the MANOVA as does the food x density interaction (indicating the effect of competition). There is no significant 4-way interaction in the MANOVA. The factors aliquot and timespan do not jointly interact with food and density to affect competition. There are three significant 3-way interactions including food, density and aliquot, and food, density and timespan, so both aliquot and timespan have an effect on competition. The entire MANOVA table is presented in S4 Table. Explanations of the abbreviations for the individual contrasts and descriptions of the meanings of the contrasts are presented in S5 Table.

Using the R squared values to rank the contrasts, it is clear that food, density and timespan are the most consequential factors in the MANOVA. The food x density x timespan 3-way interaction has the highest R squared across the interactions. The 2-way interactions: density x timespan, food x density, and food x timespan, all have much larger R squared values than the other interactions. The main factors, food, density and timespan, also have high R squared values.

The aliquot treatment and the interaction between aliquots and timespan have significant R squared values, but below most of the food, density and timespan contrasts; the interesting 3-way interaction between food, density and aliquot has the second lowest R squared of the significant contrasts.

Table 1 shows the three significant interaction contrasts involving food and density (competition): food x density x timespan (FxDxT), food x density x aliquot (FxDxA), and food x density (FxD). The standardized discriminant function coefficients for each dependent variable indicate the magnitude and sign of the contribution to the MANOVA significance for each individual contrast. For the FxDxT 3-way interaction, the Prime male age at pupation, the Average male mass at pupation and the Average female mass at pupation have the largest coefficients (all positive) followed by the negative coefficient with Prime male mass at pupation. The other variables have low coefficients for this contrast. This interaction has the highest R squared value of any interaction and indicates an effect of timespan on competition for the Prime male (mass and age) and the Average male and female (masses). For the FxDxA 3-way interaction, the largest values are for the Prime male mass and the Average male mass; they have opposite signs. The coefficients for Prime female mass and the Average female mass are also large and also of opposite sign, with the two Prime masses being positive and the two Average masses being negative. The other three dependent variables have small coefficients for this contrast (all negative). While this 3-way interaction has a low R squared value, it does indicate that there is an effect of the aliquot treatment on competition, primarily on the 4 mass variables, and of opposite direction on the Prime and Average masses. For the FxD interaction the largest coefficients are for the Prime male age at pupation, the Prime female age at pupation and the Average female mass at pupation (all positive) followed by the negative coefficient for the Prime female mass at pupation and the positive coefficient for the Average male mass at pupation. The FxD interaction has the third largest R squared value across the interactions. It represents the residual effect of food and density (competition) on the dependent variables after the higher order interactions have been accounted for. The Prime male age has a large coefficient in FxDxT interaction, but also in the FxD interaction indicating that there is an effect of competition on Prime male age at pupation in addition to the effect of the 3-way interaction. The Prime female age only has a large coefficient in the FxD interaction, indicating that the two 3-way interactions do not much affect the Prime female age, but that competition (FxD) does affect it. The FxDxA interaction does not have large coefficients for the two age variables so there is little effect of this interaction on age at pupation. The Average male and female masses have large coefficients in all three of these interactions indicating that each of the interactions has a distinct effect on the Average mass variables. The Prime male mass is affected by the FxDxT and FxDxA interactions, but the Prime female mass is affected by the FxDxA and FxD interactions. Survival is not strongly affected by any of these interactions.

**Table 1.** MANOVA contrasts for competition interactions (food, density, aliquot, timespan: FxDxT, FxDxA, FxD). R squared values, significance levels and discriminant function coefficients by dependent variable for the three interactions.

| Contrast | Survival | Prime male mass at pupation | Prime male age at pupation | Average male mass at pupation | Prime female mass at pupation | Prime female age at pupation | Average female mass at pupation | MANOVA P< | R squared |
| --- | --- | --- | --- | --- | --- | --- | --- | --- | --- |
| F x D x T | 0.199 | -0.355 | 0.68 | 0.79 | -0.098 | 0.191 | 0.646 | 0.001 | 0.87 |
| F x D x A | -0.085 | 1.579 | -0.053 | -1.67 | 0.515 | -0.127 | -1.175 | 0.001 | 0.22 |
| F x D | 0.126 | -0.097 | 0.857 | 0.322 | -0.45 | 0.503 | 0.542 | 0.001 | 0.73 |

The standardized discriminant function coefficients do not lend themselves to easy interpretation of the biology underlying the differences; however, that there are differences across the dependent variables suggests that males and females respond differently to the independent factors, and that the Prime and Average individuals respond differently within sexes. The MANOVA and these coefficients describe the relationships among the dependent variables and will be reconsidered again along with the individual ANOVA results later.

Table 2 shows the three significant interaction contrasts for the three attributes of the food supply: food level, aliquot, and timespan. The 3-way interaction between these three factors (FxAxT) is not significant in the MANOVA, but the three 2-way interactions are significant. These interactions do not include density, so do not describe competition. Instead, they describe the growth of larvae on different food regimens: the two food levels, and the distribution of the food inputs in time (aliquot and timespan). The FxA contrast has large positive coefficients for the Prime male mass and age at pupation and smaller coefficients for Survival and the Prime female mass, and for the Prime female age at pupation (negative). The coefficient for Survival is the largest for that variable across the MANOVA. The R squared value for this contrast is only slightly larger than that of the FxDxA contrast. The FxA interaction represents the residual after the FxDxA interaction is accounted for, so this is the effect of food level and aliquot on the growth of the larvae after the competition has been explained. The FxT interaction has a high coefficient for the Prime male age at pupation followed by smaller coefficients for Survival, Prime male mass at pupation, and Average male mass at pupation. The coefficients for the three variables measuring female mass and age are smaller still. The R squared value for this contrast ranks fourth across the interactions. The FxT interaction represents the residual after the FxDxT interaction is accounted for, so this is the effect of food level and timespan on the growth of larvae after the competition has been explained. The AxT interaction has large positive coefficients for Average male mass and Average female mass and a smaller positive coefficient with Prime male age at pupation and Survival and a smaller negative coefficient with Prime female age at pupation. The R squared value is the fifth ranked across the interactions (and the three lower ranked values are much smaller). The AxT interaction represents the residual after the FxAxT and DxAxT interactions are accounted for (see below for a description of the DxAxT interaction). It describes the effect of aliquot and timespan, the distribution of food in time, on the growth of larvae. These three interactions have the three highest coefficients for Survival across the interaction contrasts; the parameters of the food supply affect Survival more than any interaction with density affects it. The Prime male age at pupation is also affected by these three interactions. The Prime male mass and to a lesser extent, the Prime female mass, are affected by the FxT and FxA interactions, but not by the AxT interaction; however, Average male mass and Average female mass are affected by FxT and AxT, but not by FxA. Prime female age at pupation is slightly affected by FxA and AxT, but not by FxT. As in Table 1, the discriminant function coefficients suggest that male and female larvae grow differently in response to the independent factors. The MANOVA and these coefficients describe the relationships among the dependent variables and will be reconsidered again along with the individual ANOVA results later.

**Table 2.** MANOVA contrasts for interactions involving only attributes of the food supply (food, aliquot, timespan: FxA, FxT, AxT). R squared values, significance levels and discriminant function coefficients by dependent variable for the three interactions.

| Contrast | Survival | Prime male mass at pupation | Prime male age at pupation | Average male mass at pupation | Prime female mass at pupation | Prime female age at pupation | Average female mass at pupation | MANOVA P< | R squared |
| --- | --- | --- | --- | --- | --- | --- | --- | --- | --- |
| F x A | 0.337 | 0.597 | 0.777 | 0.002 | 0.194 | -0.214 | -0.096 | 0.001 | 0.25 |
| F x T | 0.32 | 0.345 | 0.596 | 0.371 | 0.191 | -0.025 | 0.233 | 0.001 | 0.61 |
| A x T | 0.211 | -0.094 | 0.462 | 0.721 | 0.014 | -0.191 | 0.621 | 0.001 | 0.51 |

Table 3 shows the two significant interaction contrasts for density and the non-food level attributes of the food supply (DxAxT and DxT). The 2-way interaction, DxA, is not significant in the MANOVA. The AxT interaction does not include density and is described in Table 2. Density has an interaction with aliquot and timespan, and with timespan, that is independent of the food x density (competition) interaction. However, aliquot and timespan are both characteristics of the rate of food delivery, so these interactions could indicate competitive interactions that are not related to the total food level per test tube. For this DxAxT interaction, the largest values are for the Prime male mass and the Average male mass followed by the Prime female mass at pupation. The Prime male mass is positive and the Average male mass and Prime female mass are both negative. The other coefficients are small. This interaction has the smallest R squared value of all the significant contrasts, but the 2-way interaction, DxT, has the second largest R squared value across the interaction. DxT represents the residual effect of density and timespan after the interactions FxDxT and DxAxT have been accounted for. The coefficients for Prime male age, Average female mass, Average male mass and Prime female age are large and positive. The coefficients for the other three variables are small (and negative for the Prime male and female masses). If either of these interactions describe competition based on density and one or more attributes of the food supply (independent of the food level) then the ANOVAs should show a relationship between food availability and density, such that: a) growth of larvae is good at high availability (earlier delivery: 4 aliquots and/or 3 day timespan) and low density; b) growth of larvae is less good at high availability and high density, or at low availability (later delivery: 2 aliquots and/or 6 day timespan) and low density; and c) growth of larvae is much worse at low availability and high density. As in Tables 1 and 2, the discriminant function coefficients suggest that male and female larvae grow differently in response to the independent factors. The MANOVA and these coefficients describe the relationships among the dependent variables and will be reconsidered again along with the individual ANOVA results later.

**Table 3.** MANOVA contrasts for interactions between density and the non-food level attributes of the food supply (density, aliquot, timespan: DxAxT, DxT). R squared values, significance levels and discriminant function coefficients by dependent variable for the three interactions.

| Contrast | Survival | Prime male mass at pupation | Prime male age at pupation | Average male mass at pupation | Prime female mass at pupation | Prime female age at pupation | Average female mass at pupation | MANOVA P< | R squared |
| --- | --- | --- | --- | --- | --- | --- | --- | --- | --- |
| D x A x T | 0.065 | 1.261 | 0.079 | -1.661 | -0.662 | -0.135 | 0.028 | 0.05 | 0.15 |
| D x T | 0.165 | -0.163 | 0.786 | 0.491 | -0.122 | 0.341 | 0.558 | 0.001 | 0.84 |

### MANOVA summary

The experiment looks for interactions between the aliquot and timespan treatments and food x density (representing different competitive regimes). The main effects are all significant as expected and Survival is not strongly affected by any treatments or interactions, as hoped. There is no 4-way interaction, so the aliquot treatment and the timespan treatment do not interact jointly with food x density. There is only one 3-way interaction with a high R squared value: food x density x timespan. The 2-way interactions across these 3 factors are also significant with high R squared values: food x density, food x timespan, and density x timespan. These four interactions account for the highest R squared interactions across the MANOVA. The 3-way interaction between food x density x aliquot is significant, but has a low R squared value. There is a small interaction between the aliquot treatment and competition.

The food x aliquot interaction, the food x timespan interaction, and the aliquot x timespan interaction describe the growth of larvae rather than competition among them. These interactions are not related to competition because they do not include density as a factor, but both aliquot and timespan affect the rate of delivery of food to the larvae and thus the growth and eventual pupation of the individuals.

The 3-way interaction, DxAxT, has the lowest R squared value across the significant contrasts. This interaction and DxT (the second largest R squared value) are independent of the FxD competition contrast. Aliquot and timespan are both characteristics of the rate of food delivery, so interactions between density and either, or both, could indicate other types of competitive interactions than density and food level. Alternatively, they could describe the growth of the larvae.

The benefit of the MANOVA over multiple independent ANOVAs is the relationships among the variables shown by the discriminant function coefficients for each contrast. Across the significant contrasts Average male mass and Average female mass are consistently related (both large or both small, and of the same sign). Other pairs of consistently related variables are: Prime male age and Average female mass; Prime male mass and Average male mass; Prime male age and Prime female age; Prime male mass and Prime male age; Prime male age and Survival; Prime female age and Average female mass; Prime female mass and Average female mass; and Prime female mass and Prime female age. [See the detailed numerical analysis of the ANOVAs in S1 Text for more detail.]

The discriminant function coefficients do not themselves allow for interpretation of the biology underlying these significant contrasts. The individual ANOVAs are a product of the MANOVA result and allow us to examine the effect of each contrast on the means and standard errors of the dependent variables for each contrast.

### ANOVAs

Table 4 shows the r squared values for the main treatments (summed), and the significant interactions identified by the MANOVA. The total r squared for the ANOVAs is very high (0.79-0.93) except for Survival (0.27). The full set of ANOVA results is presented in S6 Table. The main treatments (summed) and the 8 interactions in Table 4 account for almost all of the r squared totals across the dependent variables (see the row labelled “Subtotal r squared).”

**Table 4.** ANOVA significant r squared values for the main treatments and significant interaction contrasts from the MANOVA for the 7 dependent variables. The row labelled “Subtotal r squared” includes only the r squared values for the contrasts listed in this table; the row labelled “Total r squared” includes all of the r squared values in the respective ANOVAs.

| Contrast | Survival | Prime male mass at pupation | Prime male age at pupation | Average male mass at pupation | Prime female mass at pupation | Prime female age at pupation | Average female mass at pupation |
| --- | --- | --- | --- | --- | --- | --- | --- |
| Main treatments: F, D, A, T | 0.12 | 0.56 | 0.45 | 0.48 | 0.54 | 0.54 | 0.56 |
| F x D |  |  | 0.11 | 0.01 |  | 0.08 | 0.01 |
| F x A |  | 0.01 | 0.01 | 0.01 |  |  |  |
| F x T | 0.08 | 0.09 | 0.02 | 0.08 | 0.03 |  | 0.02 |
| D x T | 0.02 | 0.03 | 0.17 | 0.06 | 0.10 | 0.11 | 0.10 |
| A x T |  | 0.05 | 0.01 | 0.06 | 0.03 |  | 0.03 |
| F x D x A |  |  |  |  | 0.00 |  | 0.01 |
| F x D x T | 0.05 | 0.10 | 0.16 | 0.16 | 0.16 | 0.05 | 0.16 |
| D x A x T |  |  |  | 0.01 | 0.01 |  |  |
| Subtotal r squared | 0.27 | 0.84 | 0.93 | 0.87 | 0.87 | 0.78 | 0.89 |
| Total r squared | 0.27 | 0.84 | 0.93 | 0.87 | 0.88 | 0.79 | 0.90 |

The very high total r squared values for the mass and age variables suggest that, within the limits of this experiment, a set of equations could be developed to describe the growth and pupation of the larvae using these four factors. The current experiment, which uses only two values for each of the factors, does reveal the mechanisms of interaction, but does not provide enough points to generate these hypothetical equations. That a determinist solution exists for a model population in a microcosm is interesting, but does not mean that such a solution exists in natural systems. However, it is likely that the mechanisms of interaction described by the experiment do occur in the natural system that the microcosm emulates. It is therefore the mechanisms of competition, growth and pupation that are the investigatory targets of the experiment.

The main effects. For the four mass variables and two age variables, high food is better than low food, low density better than high density, 4 aliquots better than 2 aliquots, and the 3 day timespan is better than the 6 day timespan, where better means larger mass at pupation and earlier age at pupation. (S7 Table.) The sole exception is that Prime males pupate slightly earlier with 2 aliquots than with 4 aliquots. Survival is better at high food and low density as for the mass and age variables, but better at 2 aliquots and the 6 day timespan, the opposite of the mass and age variables. Better survival means that more larvae survived.

The interactions. The food x density (FxD) treatment combinations are designed to create different competitive environments against which to test the effects of aliquot and timespan on competition among larvae. The food x density x timespan (FxDxT) 3-way interaction is the most significant interaction contrast in the MANOVA; it is also the most significant interaction contrast in the ANOVAs for most of the variables. (The r squared values for the interaction between density and timespan, DxT, are larger for both Prime male and Prime female age at pupation than are the r squared values for the FxDxT interaction.) The timespan treatment has the greatest effect on competition among larvae. There is a much smaller interaction among food, density and aliquot (FxDxA) for the two female mass variables and an even smaller 4-way interaction for the Average mass of females (FxDxAxT, in the ANOVA only). The aliquot treatment does affect competition among females (and for the Average female mass there is an interaction between competition and both aliquot and timespan). The FxD interaction describes any residual effect of competition on the larvae (the Prime male age, the Prime female age, the Average male mass and the Average female mass).

Males and females compete differently from one another. Males are only affected by the timespan treatment, while females are also affected by the aliquot treatment.

There are two other groups of interactions: the interactions that involve only attributes of the food supply (food, aliquot, timespan); and the interactions that involve density and attributes of the food supply, but not food level (density, aliquot, timespan). The first group describes the growth of larvae independent of competition. The second group may describe competition independent of food level or other aspects of the growth of larvae.

The interactions in Table 4, and the three interactions that were not significant in the MANOVA, but had significant mean squares for one or more of the variables in the ANOVA, are interpreted by comparing the means accumulated across the interaction terms in S1 Text. The number of replicates for each treatment in the experiment, and the treatment means with standard deviations for each of the 7 dependent variables, are presented in S8 Table-S15 Table. The detailed numerical analysis of the significant interactions for the seven dependent variables is in S1 Text and the associated tables, S16 Table-S43 Table. Heuristic graphs of the relationships are presented in S5 Fig-S35 Fig. The explanation of results in S1 Text is extracted into S2 Text, condensing the numerical analysis, tables, figures and point-by-point observations into a more readable explanation. A summary of the results is presented below.

### Summary of results for Survival across experiment one

The experimental conditions were selected so that Survival would be high. Survival is treated as a dependent variable to investigate whether it is affected by the independent factors, and specifically whether it is affected by competition (FxD interactions). Survival is lower in the most competition treatments than in the other treatments. The interaction arises because the Survival is better with the 6 day timespan than with the 3 day timespan except at the lowest food level (most competition). There are two other interactions that describe the residual effects of these factors on Survival: FxT and DxT. In both interactions Survival is better at the 6 day timespan at the higher food level. Survival appears to be improved by the late addition of food at high food levels and possibly compromised by too much food in the high food, 3 day timespan treatment.

Finally, Survival is affected by the main effect of the aliquot treatment. Survival is higher in the 2 aliquot treatment compared to the 4 aliquot treatment. There are no interactions involving the aliquot treatment that are significant for Survival, so the effect of aliquot on Survival is independent of food, density and timespan. Survival is not strongly linked to competition (the FxD interaction and higher order interactions including both food and density). For this set of experimental conditions Survival and competition are largely independent.

### Summary of results for masses and ages at pupation for males and females across experiment one: the FxD set of interactions (FxDxT, FxDxAxT, FxDxA, and FxD)

The interaction between food and density is expected to produce 4 different competitive environments: least competition (high food, low density); most competition (low food, high density) and two intermediate levels of competition (high food, high density, and low food, low density). The least competition test tubes receive 32 mg of food or 8 mg food/larva. The most competition test tubes receive only 16 mg of food or 2 mg food/larva. The two intermediate competition treatments receive 32 mg (for 8 larvae) or 16 mg (for 4 larvae) of food resulting in 4 mg food/larva. Mosquito larvae should grow larger, faster, and pupate earlier in the least competition treatment, followed by the intermediate competition treatments. Pupae should be smallest, grow slowest and pupate latest in the most competition treatment. Within the intermediate competition treatments, the high food, high density treatment is expected to be better for mosquito larval growth than the low food, low density treatment because the total food is greater. These four FxD treatments are crossed with the two timespan treatments (3 days or 6 days) for a total of 8 treatment combinations. The 3 day timespan is expected to be better than the 6 day timespan because more food is offered earlier in the larval period. The interaction between the FxD treatments and the timespan treatment reveals how timespan affects the competition for food among the larvae. The four FxD treatments are also crossed with the two aliquot treatments (2 aliquots or 4 aliquots) for 8 different treatment combinations. The four FxD treatments are also crossed with both aliquot and timespan in the 4-way interaction, FxDxAxT, which is only significant in the ANOVA for the Average female mass. Finally, the FxD interaction is the residual effect of food and density (competition) on the larvae after the previous three higher order interactions have been accounted for.

The FxDxT interaction (R squared = 0.87). This interaction describes competition among larvae as affected by the timespan treatment. It is the most important interaction in the MANOVA, and in the ANOVAs, for the four mass variables and among the highest for the two age variables. Prime female mass and Prime male mass correspond to the total food in the test tube after day 4 at the higher levels of food/larva (4 mg food/larva and greater, also measured after day 4). When food is abundant Prime males and females grow as though they had access to all the resources in the test tube. Prime female age at pupation always corresponds to the total food (at high and low levels of food/larva). When food is less abundant Prime female mass and Prime male mass correspond to the food/larva. This probably indicates a change in the feeding behavior of females, switching from actively filtering to retaining particles in their guts. The Average female mass always corresponds to the food/larva. Because the Average female mass includes the Prime female mass, the non-Prime females must do less well relative to the Prime female at the higher levels of food/larva; the Prime females grow larger in response to the total food so the non-Prime females do not do even as well as the food/larva would indicate. Competition among females is explained by total food and by food/larva. [Note that total food equals density times food/larva, so density is implicit in the two measures.] Non-Prime females do not appear to benefit from the late addition of food on day 6 or any release from competition after the pupation of the Prime female. The Prime male age at pupation does not correspond to either total food or food/larva. Prime males mostly pupate on day 5. Some Prime males delay pupation slightly when food is abundant. Other Prime males delay pupation when food is scarce and especially in the test tubes with the most competition. The Prime males that pupate earliest are in the test tubes where the Prime females are competing most intensely with the other females (at the intermediate food level and high density). Prime males and Prime females respond to different conditions in the timing of pupation. Average male mass also does not correspond to either total food or food/larva. Non-Prime males grow larger after the Prime male pupates and also on the final aliquot of food in the 6 day timespan treatment. Another difference between the Average male mass and the other three mass variables occurs in the intermediate treatments. The high food, high density, 3 day timespan treatment results in increased competition which benefits the Prime male over the non-Prime males; the Prime males grow faster and larger on the higher total food and the non-Prime males are smaller than the non-Prime males in the low food, low density, 3 day timespan treatment. This suggests that males compete differently or for a different aspect of the food supply than females. The Prime male dominates competition among males, and females dominate competition in general through their larger size and retention of particles, so the non-Prime males are most affected by competition and the reduced food levels that increase competition. Prime males appear to minimize their age at pupation, reaching a minimum size determined by environmental conditions, and pupating. Non-prime males may adopt a different life history strategy. It is possible that after missing the early pupation target, these male larvae grow to a larger size on the increased food in order to benefit from being a larger adult rather than just being another small male.

The primary effect of timespan on competition is through changing the total food and food/larva over time. Males and females respond differently to competitive stress; Prime males pupate consistently early and respond to competition by pupating at a lower mass. Prime females require more total food (16 mg in this experiment) in order to pupate at all, and extend their larval lifespan to grow larger in response to additional food late in the lifespan in those treatments where the food was insufficient to pupate. At the same time, Prime and Average females and non-Prime males do not grow as large on the equivalent food provided late in the lifespan compared to that provided earlier. There may be physical constraints (size of body parts at the molt from 3rd instar) that cause this observation in addition to possible physiological constraints. Prime males, Average males, Prime females and Average females all do relatively well at the lower levels of competition (when the food/larva is 4 mg/larva or greater), but there are large differences across the sexes and smaller differences within the sexes when competition is more intense (when the food/larva is less 4 mg or less). The Prime male is strongly affected by competition (most competition, 6 day timespan) pupating at the smallest size and latest across the interaction. The Average male grows larger than the Prime male in this treatment due to the release of competition after the pupation of the Prime male, and also the effect of the additional food on day 6. The Prime and Average females are also smallest in this treatment combination, but they are more similar to the most competition, 3 day timespan treatment. They are less affected by timespan when competition is intense and more affected by timespan when competition is intermediate than the males. Females may change the way that they feed (filtering versus retention of particles) as the total food and/or food/larva changes and this may explain the difference between the 3 day timespan and 6 day timespan treatments at the intermediate levels of competition.

The FxDxAxT interaction (not significant in the MANOVA). The 4-way interaction is only significant for the Average mass of females in the ANOVA. A large input of food on day 3 effectively eliminates competition among females for a period of time, allowing the non-Prime females to grow larger relative to the non-Prime females that received equivalent food earlier in the life cycle. A similar large input of food on day 6 does not have the same effect. For females, food early in the larval period is more beneficial than food later, and multiple inputs are better than single inputs, but a large input of food on day 3 benefits the non-Prime females by eliminating competition at a critical point in the larval growth.

Female larvae outcompete male larvae, but neither the Prime male mass nor the Average male mass is affected by this interaction and the elimination of competition due to the large food input on day 3. The lack of effect of this specific food input suggests that either the food level is already so high in this treatment that males do not respond to the additional food, or that the final mass of males is already determined by the food level before day 3 (at this high food level, some non-Prime males grow larger than the Prime males in treatments at low food levels, so the second alternative is less likely).

The FxDxA interaction (R squared = 0.22). The FxDxA contrast is only significant for the Prime female mass and the Average female mass in the ANOVAs; the interaction indicates that the aliquot treatment does have an effect on competition among female larvae (in addition to the FxDxAxT interaction described above). Multiple food inputs (the 4 aliquot treatment) result in larger sizes at pupation for Prime and Average females. Multiple food inputs appear to reduce competition among females, but have no effect on competition among males. Prime females and Average females grow larger on multiple food inputs even when the food/larva is the same.

Competition among females does not seem to be affected by the different initial inputs (day 0) or by the second input (on day 1 or day 2 depending on the timespan treatment), but rather by the third input (on day 2 or day 4 depending on the timespan treatment). The third aliquot of food changes the competitive environment among females, probably causing them to switch from retention to active filtering. This effect is more pronounced in the test tubes with the least competition and those with the higher food level, probably because the third aliquot is larger in these treatments. In contrast, the females in the most competition treatments are smaller and these Prime females grow slowest across the interaction. The Prime females in the most competition test tubes are more similar in size (across the aliquot treatment) than the Prime females in the test tubes with the least competition and the intermediate levels of competition. The Average females in these test tubes are also closer in size (across the aliquot treatment) than the Average females in the test tubes with the least competition and the intermediate levels of competition. This suggests that at low food/larva levels females retain particles and differences in outcome are caused by differential rates of absorption rather than differential filtering ability. Large females probably have faster rates of absorption, but the difference is not as great as size differences and growth rates due to filtration ability.

Males are affected by competition with females in the interactions FxDxT (above) and FxD (below) as well as in other sources (see [1]). That they are not affected by competition with females in this interaction suggests that the third aliquot of food that is driving this effect among females is too late to affect the outcome of competition among males (see the FxDxAxT above). One possibility is that there is already surplus food at the higher food level for the males to grow and pupate, and that at lower food levels the females monopolize the smaller incremental food input so that males do not benefit. Another possibility is that males determine their final size before day 2 or day 4 and subsequent inputs of food don’t affect that target size. A third possibility is that the molt to the third or fourth instar limits the final size of the male pupa due to some physical constraint or some aspect of the feeding apparatus.

The FxD interaction (R squared = 0.73). This interaction represents residual effects of competition on the seven dependent variables (the primary effects being in the higher level interactions, FxDxT, FxDxAxT and FxDxA, described previously). It is the third largest interaction in the MANOVA, but it is only significant for four of the dependent variables in the ANOVAs: Prime male age, Average male mass, Prime female age and Average female mass. This contrast is much more important for the two age variables than for the two Average mass variables. Increased competition decreases the Average masses and also increases the Prime ages at pupation. Competition (food x density) appears to have a residual effect on the Prime male and female ages at pupation and the Average male and female masses at pupation that is independent of the aliquot or timespan treatments. Because this interaction affects the Average masses and not the Prime masses, the residual competition is affecting the mass of the non-Prime individuals.

The Prime female age followed the total food in the interaction FxDxT; the effect of that interaction has been removed and the residual effect of FxD on Prime female age remains. In this interaction both the food/larva after day 4 and the total food after day 4 influence the Prime female age at pupation, but the females in the test tubes with the most competition pupate disproportionately later than the other treatments.

The non-Prime females in the most competition treatment are disproportionately smaller than their counterparts in the other treatments. There is also an effect on the relative advantage of Prime females over the non-Prime females in the high food, high density, intermediate competition treatment. The size of the Prime female is reduced relative to the least competition treatment, but the Average females are even smaller, indicating that the increased density/competition primarily affects the non-Prime females.

The age of the Prime male increases disproportionately with increasing competition, so the Prime male grows to a size dependent on the main effects of food and density, but takes longer to reach that size and pupate as competition increases.

The non-Prime males in the most competition treatment are disproportionately smaller than their counterparts in the other treatments, but they are larger than the Prime male in the most competition treatment. This is likely due to the competitive release of non-Prime males after the Prime male pupates. This anomaly is numerically much smaller than the competitive release plus added food observed in the FxDxT interaction above. There is also an effect on the relative advantage of Prime males over the non-Prime males in the high food, high density, intermediate competition treatment. The size of the Prime male is reduced relative to the least competition treatment, but the Average males are even smaller, indicating that the increased density/competition primarily affects the non-Prime males.

Prime males respond to food and density (and competition with females) differently than Prime females. The Prime female pupates earliest in the test tubes with the least competition; the Prime male delays pupation in response to abundant food in those test tubes. The Prime females’ estimated growth rates reflect their ages at pupation; the Prime males’ estimated growth rates reflect their masses rather than their ages at pupation. The Prime males pupate in a much tighter window of time than the Prime females, so there is more variability in the masses than in the ages at pupation. Prime males pupate earliest in the test tubes where the Prime females appear to dominate the non-Prime females to the greatest degree, the high food, high density treatment. The more intense competition among females in these test tubes compared to the least competition test tubes appears to cause the Prime male to pupate sooner and at a smaller size.

There are two quantitative differences in the relationship between the Prime male mass and the Average male mass that are different from the females. First, the Average male mass is greater than the Prime male mass in the most competition treatment; at least one non-Prime male grows larger than the Prime male after the Prime male pupates. Second, the Prime male mass is larger in the high food, high density treatment than in the low food, low density treatment (same food/larva, but more total food in the high food, high density treatment), but the Average male mass is larger in the low food, low density treatment. In the high food, high density test tubes there is the same amount of food as in the least competition ones, but 4 more larvae. Competition among the larvae is more intense and this appears as a reduction in the amount of food. Because there is a high food level the nature of competition among the females is active filtering and rapid passing of the particles through the gut, the distribution of sizes is broad and the relative advantage of the Prime male over the non-Prime males is large. In the low food, low density test tubes, the total amount of food is half of that in the high food treatments, but the lower density means that the food/larva is the same across the two intermediate competition treatments (4 mg/larva). In this case, the lower number of particles causes the females to switch to retaining particles in their guts, further reducing the number of particles available for the males and compressing the size distribution. The Prime male is reduced in size and the Average male mass is similar to that in the high food, high density treatment, so the non-Prime males do better in the low food, low density treatment than in the high food, high density one (both relative to the Prime male in the low food, low density treatment and absolutely against the non-Prime males in the high food, high density treatment). The Average male mass is larger than expected at both low food treatments. The non-Prime males grow larger after the Prime male pupates in the treatments where the females are retaining food because of the low number of particles. The Prime male pupates, releasing food particles and reducing competition; this benefits the non-Prime males and allows them to grow larger. Timespan and aliquot are not involved in this interaction, so this effect (competitive release) is not affected by those factors.

### Masses and ages at pupation for males and females across experiment one: Summary of the FxD set of interactions (competition)

Prime females: The interaction FxDxT shows that the Prime female mass at pupation follows the total food level (on day 4 of the experiment) when the food/larva is 4 mg or greater, but follows the food/larva level at lower levels. This is probably related to the mechanism of feeding (active filtering versus retention) associated with the abundance or scarcity of food particles. Late additions of food do not have the same value as equal earlier inputs. The Prime female age at pupation follows the total food level across all treatments. In the interaction FxDxA, the Prime female mass follows the food/larva (on day 4 of the experiment) across all treatments. Late additions of food do not have the same value as equal earlier inputs in this interaction either. The Prime female age at pupation is not affected by the FxDxA interaction. In the FxDxA interaction, the third aliquot of food (only in the 4 aliquot treatment) appears to positively affect the mass of the Prime females. For the Prime female, the timespan treatment reveals that competition changes in response to the highest levels of total food at high levels of food/larva, but that the timing of pupation is related to the total food. The FxDxA interaction reveals that multiple inputs of food are beneficial and that the third aliquot results in additional growth, probably by providing more food at a critical time in the larval lifespan. The FxD interaction shows that competition affects the timing of pupation while other factors (and other interactions) affect the mass at pupation. The mass at pupation of the Prime female is determined by one set of environmental parameters, and the age at pupation is determined by another set, which partially overlaps the first set. Mass and age at pupation are partially independent, but both respond to some of the same environmental factors (e.g. total food). When these factors change during the larval life, early changes have a greater affect than later changes. This could be because of biochemical and physiological characteristics of the larvae and/or physical characteristics such as the size of the head capsule or the feeding apparatus.

Average females: The interaction FxDxT shows that the Average female mass at pupation follows the food/larva level (on day 4) across all treatments. Average females do not appear to differentially benefit from the late addition of food on day 6 or any release from competition after the pupation of the Prime female. The Prime female controls the food supply and benefits exclusively from the high total food, while the Average females grow according to the food/larva level. The interaction FxDxAxT reveals that a large input of food on day 3 effectively eliminates competition among females for a period of time, allowing the non-Prime females to grow larger relative to non-Prime females that received equivalent food earlier in the life cycle. The specific timing of this food input makes food particles available when the Prime female would be switching from active filtering to retention, and allows the non-Prime females to grow larger (increasing the Average female mass). The FxDxA interaction reveals that multiple food inputs result in larger masses for Average females. Food/larva (on day 4) describes the size order of the mass at pupation of the Average females. Early food inputs are more important than later ones for the Average females (but see FxDxAxT above). The FxD interaction reveals that the mass of the non-Prime female is affected by competition after the higher order interactions are removed; the residual effect of food and density jointly on the mass of the non-Prime female is independent of the two other factors: aliquot and timespan.

The Prime female dominates competition in the test tube and solely benefits from the high total food level. The non-Prime females grow in response to the food/larva, although they can benefit from a large input of food (on day 3 in this experiment) that coincides with the switch from active filtering to retention by the Prime female as the relative abundance of particles decreases. Food is probably unlimited for 1^st^ instar larvae and similarly abundant for 2^nd^ instar larvae. As the larvae molt from the 2^nd^ instar to the 3^rd^ instar, their demand for particles increases and their ability to filter particles increases, reducing the availability of particles. During the 3^rd^ or 4^th^ instar, the demand for particles exceeds the availability and the females switch from actively filtering particles and passing them rapidly through their guts to retaining the particles to extract more nutrients. This is further exacerbated because the quality of the particles decreases over time. An input of food in the 3^rd^ instar would offset the relative decrease in particles and allow the females to continue to actively filter. Multiple such inputs would result in larger females by reducing competition as well as increasing food availability. The timing of a food addition in the third instar may offset multiple earlier inputs (of equal food) because the non-Prime females escape competition at the time when the Prime female would normally switch to retention, reducing the growth rate of all the females and reducing their eventual mass at pupation. Individual females grow as fast as possible in response to environmental conditions. As 1^st^ instars, food may be unlimited, but chance and the initial size of the larva (or egg size, or some other uncontrolled factor in the experiment) determine how fast the larva grows. At some point, this larva molts to the 2^nd^ instar. The larva may delay molting in order to grow larger because food is available, but there is also an advantage to molting in order to use the larger feeding apparatus of the 2^nd^ instar. This trade off between growing larger and molting should also be true for the 3^rd^ and 4^th^ instar molts. Other similar trade offs may be important for the timing of pupation.

In this experiment, the total food and food/larva in the test tubes change depending on the aliquot and timespan treatments. Both total food and food/larva affect competition among females; timespan is more important than aliquot, and their effects are independent (with the exception of non-Prime females in the FxDxAxT interaction). It appears that the environmental parameters on day 4 of the larval lifespan are the most important to determining the final size and age at pupation of females, although at lower total food levels females do not pupate until there is 16 mg total food in their test tubes (after the final food input for the 6 day timespan treatment).

Prime males: Males are affected by competition with females [1]. The Prime male benefits from reduced competition with females (at higher food levels, for instance) usually at the expense of non-Prime males. This suggests that males compete more intensely among themselves than with females. The Prime male mass at pupation is affected by total food and food/larva in the same way as the Prime females (on day 4 of the larval period), but Prime males are smaller and more similar in size to each other at pupation than are the Prime females. Competition with females may account for some of the compression of the size distribution, but the early, largely simultaneous pupation on day 5 suggests that Prime males minimize the time to pupation rather than maximizing size at pupation. Prime male age at pupation is affected by food and competition. In two treatments, Prime males pupate slightly after day 5 apparently because food is abundant (3 day timespan); both of these pupate at larger masses than their 6 day timespan peers. In the two most competition treatments, Prime males pupate slightly after day 5, with the 6 day timespan treatment pupating mostly on day 6, but still before the addition of food at the end of day 6. The mass at pupation of Prime males is determined by food, but can be modified by delaying pupation to take advantage of abundant food, or to continue growing because food is scarce. The aliquot treatment does not interact with competition for Prime males; there is no effect of dividing the total food into 2 or 4 aliquots. In the FxD interaction, the mass of the Prime male is determined by other factors, but the age at pupation is affected by the residual competition after the higher order interactions have been removed. Prime male age at pupation is earliest with the high food, high density treatment, a little later in the least competition and low food, low density treatments and disproportionately delayed in the most competition treatment.

Average males: The Average male mass responds to food level, density, competition and timespan, but is not affected by the interactions between competition (FxD) and aliquot. The Average male mass deviates from the pattern shown by the Prime male and female mass variables and by the Average female mass. Non-Prime males experience a release from competition after the Prime male pupates, and in the 6 day timespan treatments, they receive additional food and appear to grow larger as a result. Prime males outcompete non-Prime males in the high food, high density, 3 day timespan resulting in larger Prime males and smaller non-Prime males compared to the equivalent food/larva treatment, low food, low density, 3 day timespan. Despite the same food/larva (4 mg food/larva) the Prime males are smaller and the Average male mass is larger at the low density than at the high density. There is also a residual effect of competition on the size of non-Prime males (in the FxD interaction). Non-Prime males decrease in size in the treatments with increased competition, but they are disproportionately small in the most competition treatment (FxD). Despite the small size of the non-Prime males in the most competition treatment (FxD), at least one of them grows larger than the Prime male so that the Average male mass is larger than the Prime male mass. This is similar to the observations in the FxDxT interaction. There is no timespan treatment here, so the increase in size of the non-Prime male over the Prime males must be due to the release from competition rather than additional food. There is also evidence that competition among males is more intense in the high food, high density treatment than in the low food, low density treatment (with the same food/larva). The Prime male is larger, the Average male mass is smaller and the difference between them is larger in the high food, high density test tubes compared to the low food, low density test tubes.

In the FxDxT and FxD interactions, Prime and Average males are largest in the least competition treatments, and smallest in the most competition treatments. The high density intermediate competition treatments (4 mg food/larva) produce larger Prime males and smaller Average males than the low density intermediate competition treatments (also 4 mg food/larva). The largest difference between Prime and Average males is in the high density intermediate competition treatments. Average males grow larger than the Prime males in the most competition treatment; this represents the incremental difference in the size of the non-Prime males. In the FxD interaction, the Average male is 0.01 mg larger than the Prime male. In the FxDxT interaction, the Average male is 0.14 mg larger. The FxD increment represents only the release of competition after the Prime male pupates, while the FxDxT increment represents both the release of competition and the additional food at the end of day 6 in the 6 day timespan treatment. The effect of the release from competition with the Prime male is much smaller than the effect of the additional food after day 6.

In the FxDxT interaction, the total food after day 4 explains the size order of Prime males when the food/larva is 4 mg or above. The food/larva after day 4 explains the size order of Prime males below 4 mg food/larva. The outcome of competition for the Prime males is directly related to the availability of food after day 4. This is not true of the Average male mass. The outcome of competition among males in the 3 day timespan treatments of the FxDxT interaction resembles that of the FxD interaction. The outcomes for males in the 6 day timespan treatments are divergent. Prime males are smaller in the 6 day timespan for each treatment. The non-Prime males grow larger in the 6 day timespan treatment than in the 3 day timespan treatment in the least competition test tubes. The non-Prime males grow larger than the Prime males in the most competition test tubes. Both are due to the release from competition and the additional food after day 6. In the intermediate competition treatments (4 mg food/larva), the effect of the 6 day timespan is to reduce the size of both Prime and Average males, and to reduce the difference between them at the high density, but to increase that difference at the low density. This is different from the effect of the 6 day timespan on Prime and Average females in the intermediate competition treatments.

Competition among males is affected by females, but males appear to be competing more intensely among themselves than directly with females. Males grow to a size determined by various environmental factors and pupate earlier and at a smaller size than females. However, males that fail to pupate early may delay pupation and grow larger, suggesting that adult longevity and other benefits of larger size may be viable alternatives to early emergence.

### Summary of the results for masses and ages at pupation for males and females across experiment one: Interactions not involving FxD competition

The previous four interactions describe competition (FxD) and the way that the factors timespan and aliquot affect competition (FxDxT, FxDxA, and FxDxAxT). Timespan and aliquot change the amount of food in the test tubes over time and this influences the nature and intensity of the competition among the larvae in those test tubes. The remaining 7 interactions describe the separate effects of food and density on timespan and aliquot (FxT, DxT, FxA, DxA), the interaction between timespan and aliquot (AxT) and the separate effects of food and density on that interaction (FxAxT, DxAxT).

FxAxT, then FxA, FxT and AxT. These four interactions only involve factors that affect the food supply. They describe the growth of larvae, not competition between larvae. The larvae are competing with each other in the test tubes, but without density as a factor in the interaction, the effect of competition is averaged out across the densities.

The FxAxT interaction (not significant in the MANOVA). The interaction of food, aliquot and timespan is significant only for the Prime female mass and the Average female mass in the ANOVAs, and not significant at all in the MANOVA. However, the 2-way interactions between these three variables are significant in the MANOVA. It is necessary to account for the 3-way interaction before considering the 2-way ones. This interaction is between three characteristics of the food supply. No density is involved, so the interaction describes growth rather than competition. The behavior of the female larvae may still be affected, switching from active filtering to retaining particles, but the end points, mass and age at pupation indicate the nature of growth on different characteristics of the food, not different competitive environments.

The Prime female masses are arranged in 3 groups with large gaps between each group. The largest Prime females are in the high food treatment; the two largest receive 32 mg of food by day 3 and the third largest receives 24 mg of food by day 4. The next group receives 16 mg of food on day 0 or by day 3. The third group doesn’t receive 16 mg of food until day 6. Prime females require 16 mg of food to pupate, so the last group delays pupation compared to the first two groups. The total food is the most important factor, but the early delivery of food (3 day timespan and 4 aliquots, jointly) increases the size of the Prime females.

In the group with the highest food level the Prime females in the 4 aliquot, 3 day timespan treatment grow largest, but those in the 2 aliquot, 3 day timespan treatment grow fastest and pupate earliest. The large input of food on day 3 appears to accelerate the growth of females. The timing of the input and/or the relative abundance of food particles affects the triggers for pupation in the Prime female at high food levels.

Within the middle group of Prime females, the ones that receive 16 mg of food on day 0 grow largest and fastest, followed by the ones in the 4 aliquot treatment (4 mg of food per day on day 0 - day 3). The Prime females in the 2 aliquot treatment (8 mg of food on day 0 and day 3) are smaller and grow more slowly than the other two. Early delivery of food (1 large input, then 4 aliquots, then 2 aliquots) increases the size of Prime females and their growth rates at moderate food levels. The estimated growth rates of the Prime females in middle group correspond to their masses. There is no acceleration due to the large input on day 3 similar to that observed at the high food level.

At the low food level and the 6 day timespan, the Prime females do not have enough food to pupate until after the last aliquot on day 6. They grow larger and faster on the 4 aliquot treatment than the 2 aliquot treatment. Despite equal food and a longer larval period, they do not grow as large as the Prime females in the 3 day timespan treatments. Food early in the larval period is more important than later additions for both size and growth rate.

The Average female masses are also arranged in 3 groups with large gaps between each group. In the group with the highest food level the Average female mass in the 4 aliquot, 3 day timespan is the largest and the closest to that of the Prime female mass. In this treatment with abundant food, the non-Prime females do better than in any other treatment. There is a bigger difference between the Average female mass in this treatment and the next largest Average female, than between the corresponding Prime females, so the non-Prime females do better on the 4 aliquot treatment than on the 2 aliquot treatment (at the high food level and 3 day timespan).

There is a larger gap in size between the 3 largest Average females and the middle group of Average females (compared to the same gap for the Prime females). This is due to the smaller size of the Average females in the high food, 2 aliquot, 6 day timespan relative to the Prime female in that treatment and also relative to the group of largest Average females. The three treatments in this middle group all receive 16 mg of food either on day 0 or by day 3. The Prime females that receive 16 mg of food on day 0 grow larger than the other two, but the Average females in this treatment do not. The Average females in the 4 aliquot treatment do better than the other two, so again the non-Prime females benefit from the 4 aliquot treatment. There is also a smaller difference between the Prime female mass and the Average female mass in the 4 aliquot treatment, compared to the other two in this middle group. However, the difference between the largest and smallest of these three Average female masses in the middle group is smaller than the difference for the middle group of Prime females. This may indicate that the food supply attributes, aliquot and timespan, are less important to the non-Prime females than to the Prime females in this middle range of food.

The gap between the middle group of Average female masses and the two smallest Average female masses is about the same as that for the corresponding Prime females. The Average females in the 4 aliquot treatment do better than the ones in the 2 aliquot treatment (both in test tubes with low food, and 6 day timespan). The difference between the Prime and Average female masses also indicates that the non-Prime females grow relatively larger in the 4 aliquot treatment. The Prime and Average female masses in the low food, 2 aliquot, 6 day timespan treatment are the smallest across the interaction, and the difference between the Prime and Average female masses is the second largest, so the non-Prime females face the worst food environment in this treatment.

Large initial inputs of food allow the Prime female to grow faster and to dominate the food supply. Smaller, regular inputs (4 aliquot treatment) allow the non-Prime females to grow larger than equivalent amounts of food in fewer, larger inputs. The food level is the most important factor for the Prime females, followed by the interaction between aliquot and timespan. The interaction between aliquot and timespan is relatively more important for the non-Prime females, especially at the highest and lowest food levels. The final aliquot in the 6 day timespan appears essential to Prime females at the lowest food level, but does not appear to benefit the Prime females in the other treatments (because they pupate before the addition of the food), or the non-Prime females (perhaps because the size distribution is fixed by the molt into the fourth instar).

The FxA interaction (R squared = 0.25). The FxA interaction indicates whether the apparent amount of food changes depending on the number of aliquots it is divided into. There is no density involved, so this examines the effect of food and aliquot on growth not competition. This interaction is significant for the Prime male mass and age at pupation and the Average male mass at pupation in the ANOVAs. The three variables: Prime male mass and age, and Average male mass, are not significant in the FxDxAxT, FxDxA, or FxAxT interactions described earlier. This is the highest order interaction between food and aliquot for males.

This interaction shows that the growth of males is affected by the both the amount of food and the number of aliquots it is divided into, separately from competition for food (food x density) and timespan. Both Prime and non-Prime males grow better on 4 aliquots than 2 aliquots. It appears to be the food delivered in the day 2 to day 4 period (the third aliquot) that causes all the male larvae to grow larger in the 4 aliquot treatment (the main effect of aliquot). This interaction between food and aliquot arises because males grow larger than expected and take longer in the low food, 4 aliquot treatment. Males are growing rapidly and filtering particles, so the added particles accelerate their growth, and this has a larger effect at low food levels than at high food levels. Prime males defer pupation and extend their growth in response to the third aliquot of food, and grow larger than Prime males at the same food level and the 2 aliquot treatment. Although the Prime and Average males are smaller in the low food treatment, the third aliquot has a larger impact on growth at the low food level compared to the high food level.

The low food treatment causes the females to switch from active filtering to retaining particles in their guts, reducing the available particles. This reduces the size and size distribution among the females, and also reduces the size and the size distribution among males. Males are smaller than projected at the low food level, especially in the 2 aliquot treatment, but the size distribution of the males is also compressed. This can be explained by exponential growth processes. At the low food level, Prime males have less of an advantage over the non-Prime males than at the high food level. Prime and Average male masses in the low food, 4 aliquot treatment are larger than in the low food, 2 aliquot treatment, and are almost as large as their projected values. This is due to the third aliquot of food that is delivered in the middle of the larval growth period (day 2 or day 4, depending on timespan treatment). Prime and non-Prime males grow larger but the relative advantage of the Prime female over the Prime male is smaller, so the Prime male benefits more from this third aliquot food addition than the non-Prime males. Males appear to actively filter at all food levels, unlike females, which appear to switch from active filtering to retaining particles in their guts. In this situation, the males respond sooner to the third aliquot and the Prime male grows larger relative to the Prime female. The Prime male also grows larger relative to the non-Prime males due to exponential growth processes and the initial size distribution before the third aliquot. The Prime males extend their larval period to grow larger on the extra food, and pupate latest in this treatment (across this interaction).

The Average male masses are in the same order as the Prime male masses, so all males appear to benefit from the third aliquot, but the Average male mass in the low food, 2 aliquot treatment is the same as the Prime male mass. The non-Prime males in this treatment grow as large as the Prime males. They don’t grow as large as the non-Prime males in the low food, 4 aliquot treatment; the third aliquot delivers more food earlier in the larval lifespan. The non-Prime males in the low food, 2 aliquot treatment must take advantage of the additional food in the final aliquot (on day 3 or day 6) to grow as large as the Prime male. This suggests that there is a minimum mass at pupation for males that is determined by environmental conditions before day 3. Male larvae grow until they reach that minimum, or grow larger than the minimum, even delaying pupation, in response to greater availability of food.

The main effect of the 4 aliquot treatment increases the size of all larvae compared to the 2 aliquot treatment, but for males at the low food level, it provides a disproportionate benefit. There are two different anomalies: the Prime male mass in the low food, 4 aliquot treatment is disproportionately large relative to both the Average male and to the Prime female; and the Average male mass in the low food, 2 aliquot treatment is the same as the Prime male mass in that treatment. The Prime male grows larger on the third aliquot at the low food level in relation to the Prime female. The non-Prime males grow to the same size as the Prime males at the lowest food level, suggesting a minimum mass at pupation for males depending on the availability of food before day 3.

The FxT interaction (R squared = 0.61). The interaction between food and timespan shows the residual effect after the higher order interactions have been explained (FxDxT for all variables, FxAxT for Prime and Average female mass, and FxDxAxT for Average female mass). The ANOVAs are significant for all the variables except for Prime female age. This interaction is about 3 times more important for males than for females. There is no density in this interaction, so the contrast describes the effect of food and timespan on the growth of larvae.

Males and females both grow according to the food level in the test tubes, but this food level is modified by the timespan treatment, and significantly affects the growth of both sexes. Prime females do not pupate until 16 mg have been added to the test tubes (at the end of the timespan for the low food level); males do pupate on less food. Females grow larger in stepwise increments as the amount of food increases; males also grow larger, but the increments are smaller and less regular. Both females and males are larger on the high food, 3 day timespan than projected by the main effects, and smaller than projected in the low food, 6 day timespan. The range of sizes of the Average females is slightly larger than that of the Prime females. The range of the Average males is compressed relative to that of the Prime males. Females may change their feeding behavior in response to the food inputs across the timespan, producing different size distributions. Most Prime females pupate on day 6; the Prime females in the low food, 6 day timespan pupate 2.28 days later. Most Prime males pupate on day 5; none appear to pupate late enough to take advantage of the day 6 food input. Non-Prime males do grow larger after the Prime males pupate; in some cases they grow larger than the Prime males. Females also grow larger on the late addition of food on day 6, and this may increase the size distribution suggesting that the females switch from retention to active filtering when the last aliquot of food is added.

In this interaction, females grow according to the total food in the test tubes after day 4; total food is affected by the food level treatment and the timespan treatment. The early instars experience unlimited food for the first few instars, but later instars are limited by food in some treatments. Prime and Average females are much larger than expected at the high food level and 3 day timespan treatment. There is about 0.20 mg difference in size between the Prime and Average females in each treatment. Both Prime and Average females are about 0.5 mg smaller with each decrease in total food after day 4 across the 4 treatments. The residual effect of this interaction on females (after removing the effects of the FxDxT interaction, the FxAxT interaction, and the FxDxAxT interaction) is the direct, almost linear, relationship between the amount of food after day 4 and the mass at pupation.

This interaction affects male mass three to four times as much as female mass, and it also affects the Prime male age at pupation. The residual effect on males (after removing the effects of the FxDxT interaction) differs from that for the females. Male mass at pupation is also directly related to the total food after day 4, but the three treatments with more food are closer in size for Prime males, and for Average males, while the treatment with the least total food is much smaller for both Prime and Average males. The three treatments with more food all are larger than expected and the treatment with the least food is smaller than expected for both Prime males and Average males. Prime males pupate before the final input on day 6, but they delay pupation at the highest total food and at the two lower total food levels. They grow larger and pupate later when food is most available, and also pupate later when food is least available, but they still pupate on day 5 or day 6. Prime males decrease in size only about 0.20 mg with each decrease in total food (compared to 0.50 mg for Prime females). Prime males at the lowest total food are disproportionately small (0.50 mg smaller than the next largest), nevertheless, they are able to pupate before the final addition of food on day 6. This suggests that the available food for the males is much lower in this treatment, probably due to retention by females at low food levels.

The Average male mass at pupation is larger than the Prime male mass in the low food, 6 day timespan treatment, where both masses are lowest across the interaction. There is no competition in this interaction, so the difference represents the incremental effect of the food after day 6 on the growth of the non-Prime males at the lowest total food.

This interaction describes the growth of larvae rather than competition. Males actively filter particles throughout their larval period. Females in the low food levels switch to retention at some point as the number and quality of particles decreases relatively to demand. The Prime male grows at the expense of the non-Prime males and pupates in response to unknown triggers that may include size, physiological status, age, and environmental cues (availability of food particles).

The females and the non-Prime males experience a benefit when the final input of food is added on day 6 and grow larger. Some non-Prime males grow to be larger than the Prime male on the extra food. The larger females grow even larger on the extra food and perhaps switch from retention to actively filtering, but the incremental food is less valuable than the same amount of food earlier in the larval period (for instance, the 3 day timespan). None of the larvae in the low food, 6 day timespan treatment reach the size of corresponding larvae in the low food, 3 day timespan treatment despite the extra food and longer larval periods.

The estimated growth rates of the Prime females are greater than those for the Prime males in all treatments. For females, the estimated growth rates reflect similar Prime female sizes and divergent ages at pupation (affected by other interactions and main effects, but not by this interaction). For males, the estimated growth rates reflect similar ages at pupation, but divergent sizes. The estimated growth rates are in the same order as the total food for both males and females. The largest difference between the rates for females and those for males is at the highest total food. The differences decrease as the size of the pupae and the total food decrease, suggesting that this is due to exponential growth processes.

Males and females respond to food differently after the effects of competition and aliquot have been removed by the higher order interactions. Females grow linearly in response to the total food after day 4. They required 16 mg of food in order to pupate and delay pupation until after day 6 in the low food, 6 day timespan treatment. Males also grow larger in response to more total food, and they delay pupation both when food is available and when it is limiting, but pupate mostly on day 5. Males in the low food, 6 day timespan treatment are much smaller than the other males, but the non-Prime males grow larger than the Prime males on the final addition of food on day 6. This appears to be due to exponential growth processes in response to the total food, but modified by different pupation triggers between the two sexes.

The AxT interaction (R squared = 0.51). This interaction between aliquot and timespan represents the residual effect of these factors after the higher level interactions have been resolved (FxAxT for Prime female mass and Average female mass; FxDxAxT for Average female mass; DxAxT for Average male mass and Prime female mass, below). This interaction is significant in the ANOVA for the four mass variables and the Prime male age at pupation.

Prime and Average females grow largest on the 3 day timespan treatments with only a small difference between the 2 aliquot and 4 aliquot treatments. Prime and Average females in the 4 aliquot, 3 day timespan treatment are largest; those in the 2 aliquot, 3 day timespan treatment are a little smaller, but still larger than their expected values. Both Prime and Average females do much worse than expected at the 2 aliquot, 6 day timespan treatment. The obvious reason for this is that there is less food in these test tubes than in any of the others; the first aliquot of food is delivered on day 0 and no more food is delivered until after pupae are collected on day 6. The Prime females pupate almost 2 days after the final aliquot is delivered, but do not grow as large as any of the Prime females in other treatments. This is an effect of the distribution of food in time independent of the food level, density, or competition (food x density).

The difference between the Prime female mass and the Average female mass indicates changes in size distribution of the females and the relative advantage that the Prime female has over the non-Prime females. The difference between the mass of the Prime female and that of the Average female is smallest in the test tubes where the females grow largest (4 aliquots, 3 day timespan) and greatest in the test tubes where the females are smallest (2 aliquots, 6 day timespan). This is the opposite of the effect due to exponential growth processes. The 4 aliquot, 3 day timespan combination provides the most food, earliest in the larval cycle across both food levels and both densities; the masses of the Prime and Average females, and the estimated growth rate of the Prime female are largest in this treatment. The difference between the Prime and Average female mass is smallest, which indicates that all the females benefit from this food delivery treatment.

The temporal distribution of food affects the actual availability of food. Furthermore, the value of identical aliquots of food decreases as the timespan increases, so late additions do not increase the size of females as much as the same input delivered earlier in the larval lifespan.

Prime and Average males also grow largest on the 3 day timespan treatments with only a small difference between the 2 aliquot and 4 aliquot treatments. Prime and Average males in the 2 aliquot, 3 day timespan treatment are largest and much larger than their expected values; those in the 4 aliquot, 3 day timespan treatment are a little smaller, but still larger than their expected values. This is different from the Prime and Average females above. Both Prime and Average males do much worse at the 2 aliquot, 6 day timespan treatment than expected. This is similar to the Prime and Average females. Prime males pupate before the day 6 food input, so there is even less food for them than for the Prime and Average females.

In contrast to the Prime females, the Prime males pupate latest in the 4 aliquot, 6 day timespan treatment, but still before the final input of food on day 6. The Prime males in both 6 day timespan treatments delay their pupation to grow larger, but the ones with more food (3 aliquots instead of 1 aliquot) delay longer and grow larger than the ones in the 2 aliquot treatment. The estimated growth rates follow the Prime male mass; this interaction affects the mass 5 times more than the Prime male age.

The difference between the Prime male mass and the Average male mass indicates changes in the size distribution of males and the relative advantage of the Prime male over the non-Prime males; it is almost entirely affected by the timespan. The difference between the Prime and Average males is larger with the 3 day timespan than with the 6 day timespan. Both Prime and Average males are larger in the test tubes with the 3 day timespan, so this is possibly due to exponential growth processes, another contrast to the observation for females.

All the Prime males pupate before the addition of the final aliquot on the 6th day. In the 6 day timespan treatments some of the non-Prime males grow larger on the final aliquot and the mass of the Average males is more similar to that of the Prime males. This is likely due to increased food for the non-Prime males after the pupation of the Prime male and the final input on day 6.

The deviations from the expected values for all 4 mass variables are greatest and negative in the 2 aliquot, 6 day timespan treatment where all of the larvae are smallest. The second largest deviations from the expected values for all 4 mass variables are in the 2 aliquot, 3 day timespan treatments; these deviations are positive. All larvae grow better than expected with a large input of food on day 3. Males grow larger and faster in the 2 aliquot, 3 day timespan treatment than in the 4 aliquot, 3 day timespan, while females grow larger and faster in the 4 aliquot, 3 day timespan. There is the same amount of food/larva in each of these treatments; males respond more to the size of the input on day 3 and females grow better on more food earlier in the larval period (the 4 aliquot treatment). Females probably switch from actively filtering particles to retention between the first input on day 0 and the second input on day 3; males respond to the second input sooner than females and this accelerates their growth in this treatment. Females continue to actively filter particles throughout the larval period in the 4 aliquot, 3 day timespan treatment because there is an input of food each day. Females dominate the food supply due to their larger size and better filtering ability, so the males don’t experience the large influx of food as in the 2 aliquot, 3 day timespan treatment. This explains the larger masses of males on the 2 aliquot, 3 day timespan and the larger masses of females on the 4 aliquot, 3 day timespan.

The amount of food in the test tubes is altered by the aliquot and timespan treatments, and this affects the growth of both males and females. The 3 day timespan treatments have the most food by day 4; the 4 aliquot, 6 day timespan has three quarters as much food, and the 2 aliquot, 6 day timespan has half as much. At the higher food level (the 3 day timespan treatment) males and females grow differently on the two different aliquot treatments. At the lower food levels (the 6 day timespan treatment) males and females are disproportionately smaller in the 2 aliquot treatment compared to the 4 aliquot treatment. Furthermore, the distribution of sizes of females is largest in this treatment (2 aliquot, 6 day timespan) while the distribution of sizes of males is smallest. The 6 day timespan treatment allows the non-Prime males to grow larger on the additional food in the final aliquot affecting the distribution of sizes among males, but not allowing the non-Prime males to catch up with non-Prime males in treatments with more food earlier in the larval period. The addition of a large amount of food on day 6 (the 2 aliquot treatment) allows females to switch from retention back to active filtering and this increases the size distribution among females, but also doesn’t allow the females to catch up to females in treatments with more food earlier in the larval period. The distribution of food (the aliquot and timespan treatments) affects the growth of males and females differently, and independently of the food level, density, and competition (food x density).

### Summary of the results for masses and ages at pupation for males and females across experiment one: density, aliquot and timespan

The remaining interactions cross density with aliquot and timespan (DxAxT, DxA, DxT). Aliquot and timespan are aspects of the food supply independent of each other and of the food level. In the previous interactions, both aliquot and timespan, and their interaction, AxT, appear to alter the availability of food for the larvae. The interaction of density (4 larvae or 8 larvae per test tube) and food level (16 mg food per test tube and 32 mg food per test tube) produces 4 different competitive environments (the FxD set of interactions, above). These three interactions between density, aliquot and timespan may also result in different competitive environments, or they may reveal additional information about growth. Competition among larvae alters the mass and age of the Prime individuals, the growth rate of the Prime individual, the Average mass, and the difference between the Prime and the Average mass. Furthermore, the specific differences are sex-dependent. For females, the 4 aliquot, 3 day timespan appears to provide the largest amount of food, earliest in the larval period (at both food levels) and represents the best growth conditions in the AxT interaction (above). Also for females, the 2 aliquot, 6 day timespan provides the least amount of food early in the larval period and represents the worst growth conditions across this interaction. The 2 aliquot, 3 day timespan and the 4 aliquot, 6 day timespan are intermediate, but the expectation is that the 3 day timespan treatment will provide more food earlier than the 6 day timespan. High density should increase competition for food and it should have a disproportionate effect at the lower food supply (both food levels, 32 mg/test tube and 16 mg/test tube, are present in each treatment combination, so this would be competition independent of the food level).

The 3-way interaction, DxAxT, is the highest order interaction for these factors for the two dependent variables, Average male mass and Prime female mass. DxT is the second most significant interaction in the MANOVA (after FxDxT). It is significant for all 7 dependent variables in the ANOVAs (also like FxDxT). DxA is not significant in the MANOVA, but is significant for Prime female age at pupation in the ANOVA.

The DxAxT interaction (R squared = 0.15). The interaction explains only a small amount of the variance for the Prime female mass and the Average male mass, but the AxT and DxT interactions are among the most significant across the ANOVAs and the 3-way interactions have to be addressed before the 2-way interactions. (The AxT interaction was addressed before the DxAxT interaction because it does not involve density as a factor and therefore describes the growth of larvae regardless of competition among them.) Because the Prime female mass is included in the Average female mass, the non-Prime females must be affected collectively in an equal and opposite manner than the Prime female. The effect of this interaction on the Average male mass but not on the Prime male mass indicates that only the non-Prime males are affected by the 3-way interaction.

If the DxAxT interaction is the result of competition for food, the Prime females should pupate at sizes that reflect the analogous treatments in the FxD set of interactions, with the AxT combinations standing in for food availability rather than food level (F). This is clearly not true. Overall, the masses of the Prime female correspond to the food availability due to the DxAxT treatments, but the size distributions among the females associated with competition (in the FxD set of interactions) do not appear. For the Prime female, the DxAxT interaction describes the growth of females rather than competition among females. As the Prime female benefits from the aliquot and timespan treatments, the non-Prime females correspondingly decrease in size (collectively) so the Average female mass is not affected by this interaction.

This interaction arises because the Prime female grows larger in the low density, 2 aliquot, 3 day timespan treatment than in the low density, 4 aliquot, 3 day timespan treatment. The main effects predict that the 4 aliquot treatment should be larger. The large increment of food on day 3 benefits the Prime female at the expense of the Average female at this high food availability. The size of the Prime female in the treatments with less food in this interaction follow the food/larva level after day 4. In the case where the food/larva is the same across cells, the total food affects the outcome, and for the two cells where the food/larva and the total food are equal, the 4 aliquot treatment is larger than the 2 aliquot treatment.

The Average male mass in the DxAxT interaction is similar to the Average male mass in the FxDxT interaction in several ways. The largest Average male occurs in the 6 day timespan treatment with the most food in both interactions. The next largest Average males occur in the 3 day timespan treatments with the most food in both interactions. There is a gap in size between the treatments with 4 mg food/larva or higher (the highest food levels and the largest Average males) and the rest of the Average males. The smallest Average males occur in the 6 day timespan treatments with the least food; in both interactions some of the non-Prime males grow larger than the corresponding Prime male so that the Average male mass is greater than the Prime male mass. This appears to be due to the release from competition after the Prime male pupates and the additional food on day 6. Finally, in each interaction there is one treatment at an intermediate food level that results in a large Prime male mass, a smaller Average male mass, and a large difference between them. The relationships between Prime male mass, Average male mass, the difference between the two masses, and the food/larva after day 4 suggest that the non-Prime males are competing in the DxAxT interaction in the same way that they are competing in the FxDxT interaction; aliquot interacts with density and timespan for the non-Prime males similarly to the way that food level does in the FxDxT interaction.

The order of the size of the Average male does not correspond to either the total food after day 4 or the food/larva after day 4. The largest Average males are in the test tubes where the Prime male has the highest estimated growth rate. In this interaction, the Prime male estimated growth rate predicts the Average male mass better than the total food or food/larva or any combination of the independent factors. The Prime male mass and age at pupation are not affected by this interaction, but the growth rate of the Prime male predicts the mass of the Average male. Growth rate is an indication of the competitive environment; high growth rates occur in the test tubes with the least competition (in the FxD interactions; estimated growth rates in the FxDxT interaction are in the same order as Prime males size and food level). This suggests that the Prime males are not affected by competition governed by aliquot and timespan, but that the non-Prime males are affected.

The DxA interaction (not significant in the MANOVA). The Prime female age is the only variable significantly affected by this interaction in the ANOVA. This is the smallest of the 4 significant interaction contrasts for Prime female age. None of the higher order interactions between density and aliquot are significant for the Prime female age at pupation, so this represents the only interaction between these factors for Prime female age. The asymmetry that causes this interaction for the Prime female age appears to be the small difference due to aliquot at low density compared to the much larger effect at high density. Prime females pupate at almost the same age at low density regardless of aliquot treatment. At the high density, Prime females pupate much later in the 2 aliquot treatment than in the 4 aliquot treatment. In the other significant interactions for the Prime female age at pupation, increased total food is related to earlier pupation. Total food does not appear to be related to age at pupation in this interaction. Prime females pupate relatively early in the test tubes with the most initial food (low density and 2 aliquots), although they don’t grow as large, as fast or pupate as early as those in the test tubes with more food during the larval period (low density and 4 aliquots). This suggests that the initial availability of food affects the trigger for pupation for females. In this interaction, the large initial input (low density, 2 aliquots) results in much earlier pupation than expected for the Prime female while not affecting Prime female mass or any other dependent variables. Whether this is due to competition or exponential growth processes is not clear. The mass at pupation of females is affected by large inputs of food on day 3 (FxDxAxT for Average females, FxDxA for all females, DxAxT for Prime females), but the age at pupation appears to be affected the amount of the initial input on day 0.

The DxT interaction (R squared = 0.84). The interaction between density and timespan shows the residual effect after the higher order interactions have been explained (FxDxT for all variables, FxDxAxT for Average female mass, DxAxT for Prime female mass and Average male mass).

Because this interaction involves density and timespan, an attribute of the food supply, it is possible that competition is involved. Food level (total food) and density are independent factors in the experiment, and they jointly affect competition among the mosquito larvae, but growth and competition among the larvae are also affected by food/larva, which is necessarily confounded with both food level and density. The DxT interaction reflects the residual effects of density and timespan on growth and competition. Timespan clearly affects the amount of food in the test tubes on a daily basis, so could be affecting the competition among larvae as well as the growth of larvae.

Referring to the discussion of FxD competition in the DxAxT interaction above, the same relationships should apply to the DxT interaction for females: the largest Prime female mass, Average female mass, estimated growth rate, and the earliest age at pupation all occur in the low density, 3 day timespan treatment. This treatment is also associated with the smallest difference in mass between the Prime and Average females, and the largest positive deviations from the expected values for the masses of the Prime and Average females. All these indicators line up with the least competition treatment in the FxD set of interactions. The smallest Prime female mass, Average female mass, estimated growth rate, and the latest age at pupation occur in the high density, 6 day timespan treatment. This treatment is also associated with the largest difference in mass between the Prime and Average females, and the largest negative deviations from the expected values for the masses of the Prime and Average females. This doesn’t correspond entirely with the outcomes from the most competition treatment (FxD set of interactions); specifically, at low food levels the Prime female should switch from actively filtering to retention and this should result in a smaller size distribution between the Prime and Average females. Despite the low food/larva level, the ongoing food inputs over the timespan appear to allow the females to actively filter particles rather than switching to retention.

The two intermediate treatments fall between the low density, 3 day timespan and the high density, 6 day timespan treatments for the masses of the Prime and Average females, the estimated growth rates, the ages at pupation, the differences in mass between the Prime and Average females, and the deviations from the expected values. If competition were the predominant factor influencing the growth of these females, the high density, 3 day timespan treatment should produce the larger, faster growing, earlier pupating females with a larger difference between the Prime and Average females. This is not true, except that the Prime females in the high density, 3 day timespan treatment pupate slightly earlier than those in the low density, 6 day timespan treatment, probably due to the higher total food after day 4 as in other interactions. The outcome of this interaction for female mass seems best explained by the food/larva rather than total food or competition. The residual effect of this interaction on the growth of females is an almost linear correspondence between the food/larva after day 4 and the masses at pupation for both Prime and Average females. Note that females in the FxT interaction had a linear residual relationship with total food after day 4.

Again referring to the discussion of FxD competition in the DxAxT interaction above, the same relationships should apply to the DxT interaction for males: the largest Prime male mass, Average male mass, and estimated growth rate, all occur in the low density, 3 day timespan treatment. This treatment is also associated with the largest positive deviations from the expected values for the masses of the Prime and Average males, but not with the earliest age at pupation or the smallest difference in mass between the Prime and Average males. All these indicators line up with the least competition treatment in the FxD set of interactions. The smallest Prime male mass, Average male mass, estimated growth rate, and the latest age at pupation occur in the high density, 6 day timespan treatment. This treatment is also associated with the largest negative deviations from the expected values for the masses of the Prime and Average males, but not with the largest difference in mass between the Prime and Average males. Non-Prime males grow larger after the pupation of the Prime males, and especially on the final input of food on day 6. Some non-Prime males in the high density, 6 day timespan grow larger than the Prime male in that treatment. For males, this interaction resembles the competition in the FxD set of interactions.

For males, the two intermediate treatments also fall between the low density, 3 day timespan and the high density, 6 day timespan treatments for the masses of the Prime and Average males, and the estimated growth rates, but not for the ages at pupation, the differences between the Prime and Average male masses, and the deviations from the expected values. Like the Prime male in the high food, high density treatment (FxD interaction), the Prime male in the high density, 6 day timespan treatment is larger than in the low density intermediate treatment, and the difference between the Prime and Average males is largest across the interaction. This resembles the competitive interaction (FxD set of interactions) and differs from the outcome for the females.

For males, the congruence between the Prime male mass, the Average male mass, the estimated growth rate, the difference between the Prime male and the Average male and the deviations from the expected values of the masses are clear. Based on the outcomes for the mass of males, the DxT interaction looks like males are competing for food due to the timespan treatment. Rather than the linear residual relationship between food/larva and mass at pupation observed for the females in this interaction, the males are disproportionately smaller in the high density, 6 day timespan treatment. The DxT interaction is analogous to the FxD interaction for males; competition appears to affect the outcome of this interaction despite the absence of food level as a factor. Furthermore, the non-Prime males experience a release from competition after the Prime male pupates, and they also grow larger on the input of food on day 6. This residual effect is on the order of the release from competition alone in FxD (0.01 mg incremental difference in size between the Average male mass and the Prime male mass).

The food/larva doesn’t govern age at pupation for males or for females in this interaction. Total food appears to be more important to the Prime female age at pupation that the food/larva in this interaction as well as in the FxDxT interaction. The Prime males pupate early in three of the four treatments in the DxT interaction. They pupate latest in the high density, 6 day timespan treatment which corresponds to the most competition treatment in the FxD set of interactions, so competition delays pupation of the Prime male in the DxT interaction as well as in the FxD set of interactions, although all the Prime males pupate before the final input of food on day 6. It is not clear what causes the Prime male to delay pupation when food is available; neither food/larva nor total food consistently predict this response. Prime males consistently delay pupation when competition is intense, but do not delay it long enough to benefit from the additional food on day 6 in the day 6 timespan treatments.

Males versus females. Density and timespan jointly affect the mass and age variables for males and females, but they affect them differently for the two sexes. Females grow largest, fastest, the Prime females pupate earliest, and the relative advantage of the Prime female over the non-Prime females is smallest at the low density and 3 day timespan. Either the 6 day timespan or the high density reduce the size, the growth rate, and increase the age at pupation and the relative advantage of the Prime female over the non-Prime females. The test tubes with the high density and the 6 day timespan produce the smallest females, the slowest growth rate, the latest age at pupation, and the largest relative advantage of the Prime female over the non-Prime females. The females do disproportionately well in the low density, 3 day timespan and disproportionately poorly in the high density, 6 day timespan treatments. The interaction between density and an attribute of the food supply could be an indication of competition independent of the food level (FxD), but there is no evidence for it. As the food availability early in the larval period changes due to a longer timespan and higher density, the females all pupate at smaller sizes, grow more slowly, and the size distribution increases. The simple explanation is that these are the result of exponential growth processes rather than competition, because females should switch from active filtering to retention at low food levels (and they appear to do so in other interactions in this experiment), and that would cause the size distribution to compress rather than to increase as observed.

Males are also largest and grow fastest at the low density, 3 day timespan treatment, and smallest, grow slowest and pupate latest at the high density, 6 day timespan treatment. They differ from the females in that the earliest age at pupation is in the low density, 6 day timespan treatment and relative advantage of the Prime male over the non-Prime males is more affected by timespan than by density. The size distribution of the males increases with density in the 3 day timespan treatments, similar to that of the females. However, the size distribution of males decreases with increasing density in the 6 day timespan treatments, probably due to the added food after the Prime male pupates. In contrast with the females in this interaction, the Prime male mass and age, the Average male mass, and the difference between the Prime and Average male masses jointly resemble the pattern in the set of FxD interactions that describe competition (above). This suggests that males are competing for food in response to the changes in availability caused by the timespan treatment independently of the food level treatment. This is another indication that males and females compete for different aspects of the food resource.

Females grow larger and faster than males in all treatments. The relative difference in size of the Prime female over the Prime male and that of the Average female over the Average male could also indicate competitive interactions. These differences correspond to the relative advantage of the Prime female over the Prime male and of the non-Prime females over the non-Prime males. With one exception, the advantage decreases as the mass of the males and females decreases, suggesting exponential growth processes. The Prime male in the high density, 3 day timespan treatment is relatively larger than in the high density, 6 day timespan (so the difference between the Prime female and the Prime male is smaller). Prime males in this treatment are also relatively larger than the Average males. The high density with the 3 day timespan increases the relative advantage of the Prime male over the non-Prime males, but also as compared to the females. This interaction does not affect competition among females, but it appears to affect competition among males and possibly between males and females.

Masses and ages at pupation for males and females across experiment one: main effects. The main effects represent the residual variance that is explained by the individual factors after the interactions have been removed. For the 4 mass variables and 2 age variables, the explained variance due just to the main effects is around 50% (45% to 56%). This is primarily an indication that the levels of the factors affected the growth of the larvae as hoped for during the design of the experiment.

There is one anomaly among the main effects: no residual effect of density on either the Prime female mass or the Average female mass; all of the effects of density on female mass are explained by the interactions between density and the 3 factors that describe the amount and timing of food inputs. This contrasts with the large residual effect of density on age at pupation for the Prime females (32%). It also contrasts with the significant effect of density on male mass, although the explained variance is low (1%, and less than 1%, for Prime and Average male mass, respectively). The Prime male age at pupation also has a large residual effect of density (31%).

### Second experiment

The endpoint of the experiment for each individual larva was either pupation, or death. The endpoint of each treatment vial was the last pupation or the death of the last larva. The experiment was designed to measure the mass and age at pupation in response to density (1-3 larvae/vial) and food level (2 mg – 5 mg food/larva). This analysis was not possible because of the differential mortality shown in Table 5. The raw data are presented in S2 Dataset.

**Table 5.**
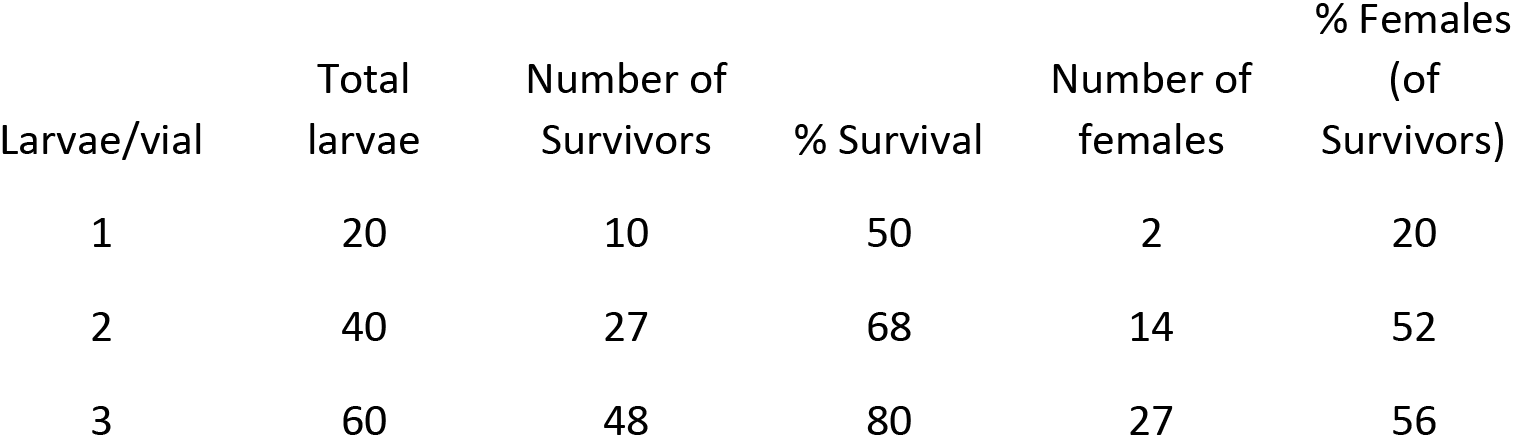
Survival of larvae across low densities showing sex ratio distortion at the lowest density.

Survival at the lowest density (1 larva per vial) was lower than in the other two treatments, and favored the survival of males over females (8 males, 2 females). The comparison between the pupae raised alone and those raised with competitors was not possible because too many cells were empty. No statistical tests were performed on the survival data (the data did not conform to the original experiment due to the differential mortality).

In Table 6, males and females are smaller at the lowest density despite the same food/larva treatments as for the vials with 2 and 3 larvae. Males at the lowest density take longer to pupate, while females take less time to pupate at the lowest density. No statistical tests were performed on the mass and age data (the data did not conform to the original experiment due to the differential mortality).

**Table 6.**
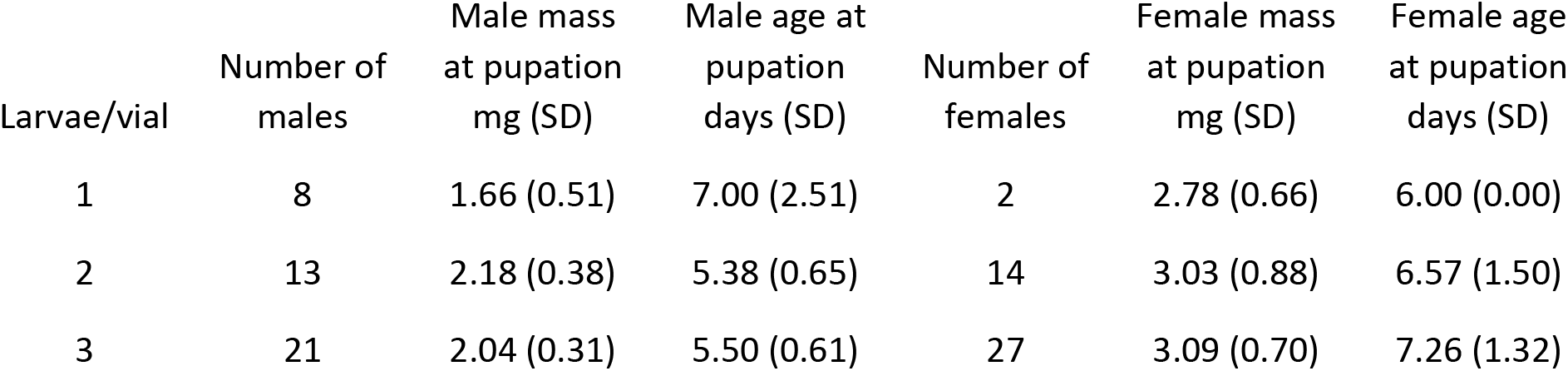
Mass at pupation and age at pupation for males and females at low densities.

| Larvae/vial | Number of males | Male mass at pupation mg (SD) | Male age at pupation days (SD) | Number of females | Female mass at pupation mg (SD) | Female age at pupation days (SD) |
| --- | --- | --- | --- | --- | --- | --- |
| 1 | 8 | 1.66 (0.51) | 7.00 (2.51) | 2 | 2.78 (0.66) | 6.00 (0.00) |
| 2 | 13 | 2.18 (0.38) | 5.38 (0.65) | 14 | 3.03 (0.88) | 6.57 (1.50) |
| 3 | 21 | 2.04 (0.31) | 5.50 (0.61) | 27 | 3.09 (0.70) | 7.26 (1.32) |

The differential mortality at the lowest density (1 larva/vial) disrupted the analysis of mass and age at pupation at low densities. It appears that solitary larvae do not survive as well and do not grow as well as larvae with one or two competitors, but this was not tested statistically. The appropriate follow on is to design an experiment to investigate this observation. (See the results and discussion in [2]. This has already been incorporated into the discussion in [1]).

The second experiment influenced the design of the third experiment. The initial amount of food per test tube is reduced to improve the survival of the single larvae. The first experiment also suggests that less food added over a longer time was better for survival (but not necessarily for growth). Survival decreases at both low levels of larval food and at high levels of larval food [1] [39].

### Third experiment

Mortality is higher for single larvae in test tubes than for multiple larvae (see the second experiment above). 50 of the 150 larvae died during the experiment or were moribund 15 days after receiving the second food input (33%). Nevertheless, there are sufficient numbers of both males and females to perform the analysis as designed. The raw data are presented in S3 Dataset.

Table 7 shows the number of pupae, their mean weight and age at pupation, sorted by treatment. The number of observations varies for several reasons. The 1^st^ instar larvae were not sexed when they were assigned to a test tube, so there is some variability due to that. Some larvae died before the second input of food, others died after the second input of food during the experiment, and another group was moribund after 15 days (the end of the experiment). (See Tables 8 and 9 below.) Table 7 shows the 50 males and 40 females that were analyzed in the experiment (56% male). In addition to those, 10 males pupated before the second input of food and were excluded from the analysis. Overall, there were 60 males and 40 females (60% male) that survived to pupation including those males that pupated before the second food input.

**Table 7.** Means (SD) for mass and age at pupation by treatment factors.

| Second food input (mg) | Day of second food input | Sex | Number of observations | Mean mass (mg) | Mass (SD) | Mean age (days) | Age (SD) |
| --- | --- | --- | --- | --- | --- | --- | --- |
| 1 mg | day 6 | M | 10 | 1.652 | 0.276 | 9.500 | 1.080 |
| 1 mg | day 6 | F | 10 | 2.086 | 0.347 | 11.900 | 1.370 |
| 1 mg | day 8 | M | 6 | 1.682 | 0.271 | 12.167 | 0.408 |
| 1 mg | day 8 | F | 6 | 2.097 | 0.275 | 13.667 | 1.033 |
| 2 mg | day 6 | M | 10 | 1.947 | 0.438 | 9.200 | 1.135 |
| 2 mg | day 6 | F | 8 | 3.034 | 0.225 | 10.625 | 0.916 |
| 2 mg | day 8 | M | 9 | 2.272 | 0.124 | 11.889 | 0.782 |
| 2 mg | day 8 | F | 6 | 2.787 | 0.165 | 13.333 | 0.816 |
| 3 mg | day 6 | M | 10 | 2.265 | 0.268 | 9.300 | 0.823 |
| 3 mg | day 6 | F | 5 | 3.744 | 0.178 | 10.400 | 0.548 |
| 3 mg | day 8 | M | 5 | 2.262 | 0.259 | 12.200 | 1.095 |
| 3 mg | day 8 | F | 5 | 3.488 | 0.281 | 13.400 | 0.894 |

**Table 8.**
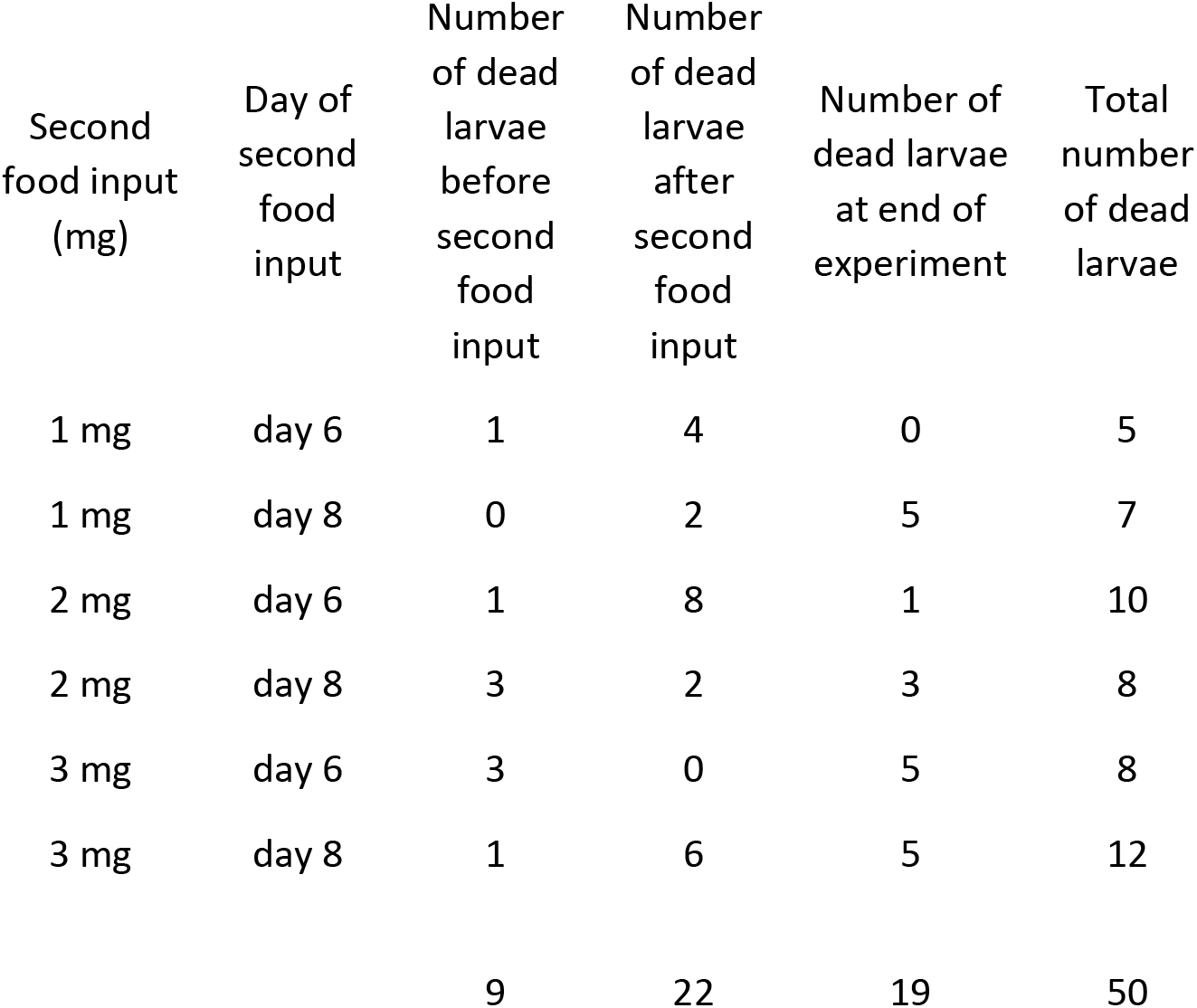
Counts and totals for larval deaths by treatment and time period.

By inspection of Table 7 the mass of the pupae increases as the second food input increases: mean mass (SE) for 1 mg is 1.88 mg (0.25), for 2 mg is 2.51 mg (0.49) and for 3 mg is 2.94 mg (0.79). The mass of the pupae with the 6 day delay is slightly larger than those with the 8 day delay: mean mass (SE) for 6 day delay is 2.45 mg (0.78) and for 8 day delay is 2.43 mg (0.63). The mass of females is larger than the mass of males: mean mass (SE) for females is 2.87 mg (0.69) and that for males is 2.01 mg (0.30). The ages at pupation are the unadjusted ages (see Methods above for an explanation of the adjustment). Males pupate before females in each treatment: the mean age for males is 10.71 days (1.51) and for females is 12.22 days (1.46). Males and females pupate later in the day 8 delay treatments than in the day 6 delay treatments: the mean age for the 6 day delay is 10.15 days (1.04) and for the 8 day delay is 12.78 days (0.77). Females in the 1 mg second food input with the day 8 delay pupate less than 2 days later than the females in the 1 mg second food input with the day 6 delay, indicating possible interactions between factors. Males and females pupate earlier at the higher second food inputs compared with the 1 mg second food input: the mean age for the 1 mg second food input is 11.81 days (1.72), for the 2 mg second food input it is 11.26 days (1.76) and for the 3 mg second food input it is 11.33 days (1.83). Males in the 3 mg second food input on the day 8 delay pupate later than males in the 1 mg second food input with the day 8 delay, again suggesting possible interactions between factors. (See also S44 Table for the main effects described above.)

Table 8 shows the number of deaths by treatment and by time period. 50 of 150 larvae died (33%). There were not enough deaths for tests of significance across the time period or food input treatments. The 6 or 8 day period of starvation is expected to be the largest source of differential mortality. 23 larvae died in the 6 day treatment and 27 larvae died in the 8 day treatment across the entire experiment.

Table 9 shows the age at death by treatment. In addition to the 31 larvae that died during the experiment, another 19 were moribund after 15 days and never pupated. 9 of the 31 larvae in Table 8 died before the second food input. No analyses were performed on this data.

**Table 9.** Mean (SD) age at death (days) of larvae by treatment.

| Second food input (mg) | Day of second food input | Number of dead larvae | Mean age at death | SD |
| --- | --- | --- | --- | --- |
| 1 mg | day 6 | 5 | 10.20 | 2.49 |
| 1 mg | day 8 | 2 | 11.00 | 2.83 |
| 2 mg | day 6 | 9 | 8.56 | 2.07 |
| 2 mg | day 8 | 5 | 8.60 | 3.71 |
| 3 mg | day 6 | 3 | 6.00 | 0.00 |
| 3 mg | day 8 | 7 | 11.17 | 2.14 |

### MANOVA

Table 10 shows the MANOVA results for mass and age at pupation for each contrast. For brevity, the contrasts are labelled Food 1, Food 2, Delay and Sex. The first contrast, Food 1, stands for the comparison between the second input of 1 mg of food and the input of 2 mg of food. The second contrast, Food 2, is the comparison between the average of the 1 mg and 2 mg inputs and the 3 mg input. Delay is the comparison between the day 6 delay and the day 8 delay. Sex is the comparison between the males and females. The main effects are all significant as are most of the interactions. The 2-way interaction between delay and sex is not significant, although both the 3-way interactions among food x delay x sex are significant. The 3-way interactions explain the joint effects of the factors. The 2-way interactions and the main effects show the residual explained variance of the factors after the higher order interactions have been accounted for. The R squared value from the MANOVA is analogous to the r squared value in the ANOVA (although the R squared values are not additive, unlike the r squared). The largest R squared values are associated with the most important contrasts. The largest R squared value across the interactions is for the 3-way interaction between food 2 x delay x sex. The second largest R squared value across the interactions is the 2-way interaction between food 1 x delay; however, there is a significant 3-way interaction that includes these two factors: food 1 x delay x sex. The 3-way interactions mean that the amount of food added and the length of the delay (period of starvation) jointly affect the growth of the larvae, and this effect is different depending on the sex of the larva. The discriminant function coefficients indicate the weight of each of the two dependent variables on the contribution to the significance of that contrast. In most of the contrasts the mass variable is more important than the age variable. In those contrasts where this is not true, the ANOVA (see Table 11) shows that the mass variable is not significantly affected by the contrast, but the age variable is significantly affected. (These contrasts are: the main effect, food 2; the interaction food 1 x sex; the interaction food 2 x sex).

**Table 10.** MANOVA discriminant function coefficients, R squared values and P values for the single df contrasts.

| Contrast | df | MANOVA<br>Mass<br>discriminant<br>function<br>coefficients | MANOVA<br>Age<br>discriminant<br>function<br>coefficients | MANOVA<br>R<br>squared | MANOVA<br>P < |
| --- | --- | --- | --- | --- | --- |
| Food 1 | 1 | 0.991 | -0.388 | 0.56 | 0.001 |
| Food 2 | 1 | -0.159 | 1.015 | 0.37 | 0.001 |
| Delay day 6 vs day 8 | 1 | 0.956 | 0.173 | 0.70 | 0.001 |
| Sex M vs F | 1 | 0.845 | 0.409 | 0.28 | 0.001 |
| Food 1 x Delay | 1 | 0.819 | 0.450 | 0.34 | 0.001 |
| Food 1 x Sex | 1 | 0.311 | 0.898 | 0.17 | 0.001 |
| Food 2 x Delay | 1 | 1.012 | -0.099 | 0.08 | 0.039 |
| Food 2 x Sex | 1 | -0.350 | 0.999 | 0.17 | 0.001 |
| Delay x Sex | 1 |  |  |  | ns |
| Food 1 x Delay x Sex | 1 | 1.015 | -0.150 | 0.16 | 0.001 |
| Food 2 x Delay x Sex | 1 | 0.850 | 0.400 | 0.44 | 0.001 |
| Residual | 78 |  |  |  |  |

**Table 11.** Mean squares (MS), r squared values and P values for the ANOVAs for the two variables mass and age at pupation for each single df contrast.

| Contrast | df | ANOVA<br>Mass<br>MS | ANOVA<br>Mass r<br>squared | ANOVA<br>Mass P < | ANOVA<br>Age MS | ANOVA<br>Age r<br>squared | ANOVA<br>Age P < |
| --- | --- | --- | --- | --- | --- | --- | --- |
| Food 1 | 1 | 6.70 | 0.18 | 0.001 | 4.45 | 0.02 | 0.034 |
| Food 2 | 1 | 0.00 |  | ns | 42.02 | 0.20 | 0.001 |
| Delay day 6 vs day 8 | 1 | 14.07 | 0.37 | 0.001 | 19.83 | 0.10 | 0.001 |
| Sex M vs F | 1 | 2.03 | 0.05 | 0.001 | 8.92 | 0.04 | 0.003 |
| Food 1 x Delay | 1 | 2.57 | 0.07 | 0.001 | 13.43 | 0.06 | 0.001 |
| Food 1 x Sex | 1 | 0.27 |  | ns | 13.61 | 0.07 | 0.001 |
| Food 2 x Delay | 1 | 0.54 | 0.01 | 0.011 | 0.04 |  | ns |
| Food 2 x Sex | 1 | 0.04 |  | ns | 13.65 | 0.07 | 0.001 |
| Delay x Sex | 1 | 0.00 |  | ns | 0.21 |  | ns |
| Food 1 x Delay x Sex | 1 | 1.18 | 0.03 | 0.001 | 0.01 |  | ns |
| Food 2 x Delay x Sex | 1 | 4.16 | 0.11 | 0.001 | 17.65 | 0.08 | 0.001 |
| Residual | 78 | 0.08 |  |  | 0.96 |  |  |
| Totals |  |  | 0.83 |  |  | 0.64 |  |

### ANOVAs

Table 11 shows the mean squares, r squared values and significance levels (P values) for mass and age at pupation for each contrast. Unlike the MANOVA, not all the contrasts are significant for both variables, but only the delay x sex interaction is not significant for either variable. As explained for the MANOVA above, the 3-way interactions indicate that the amount of food in the second food input and the delay to the second food input jointly affect the mass and age at pupation, and this effect is different for the two sexes. The food 2 x delay x sex interaction is the most significant for both mass and age (as it was in the MANOVA). The second largest interaction for mass at pupation is the food 1 x delay, as it was for the MANOVA, and there is also a significant 3-way interaction for these factors (food 1 x delay x sex). However, this is not the case for age at pupation; there is no significant 3-way interaction (food 1 x delay x sex) for age comparable to the one for mass, and there are two 2-way interactions that have larger r squared values than the food 1 x delay for age. These are: food1 x sex and food 2 x sex, and they are only significant for age, not for mass. Males and females respond differently to the food input and delay treatments; both mass and age at pupation are affected, but the treatments affect them differently.

### Third experiment result summary

The detailed analyses of the interactions for the dependent variables, mass and age at pupation, across the MANOVA and ANOVAs are presented in S3 Text and accompanying tables, S45 Table-S60 Table, and S37 Fig-S38 Fig. The results are summarized by contrast below.

Food 2 x delay x sex. Sex is more important to mass at pupation than food input or delay for this interaction. Food is more important to mass at pupation than delay. Females grow larger on the day 6 delay than on the day 8 delay at both food levels. Males grow larger on the day 8 delay at the lower food inputs, but grow to the same size at the 3 mg food input. This suggests that males reach a size determined by environmental conditions and pupate as soon as they reach that size, while females grow as large as possible. Sex is more important to age at pupation than either food input or delay for this interaction. Delay is more important to age at pupation than food input amount.

Females pupate earliest in the test tubes with the 3 mg food input and the 6 day delay. The females in the other 3 treatment combinations pupate almost a day later. The similarity of the ages at pupation of the females in the treatments with lower food inputs, and in the 3 mg food input with the longer delay suggests that females may have an optimal maximum duration of the larval period. They grow to different sizes depending on the food input and the delay, but they take almost the same amount of time after the food input to do so.

Males also pupate earliest in the test tubes with the 3 mg food input and the 6 day delay, but the males in the test tubes with the lower food input and the 6 day delay pupate shortly afterwards, followed by the lower food input and the 8 day delay, then the 3 mg food input and the 8 day delay. Males postpone pupation at the 3 mg food input and the day 8 delay to grow to the same size as the males in the 3 mg food input with the day 6 delay. This suggests that these males are replenishing physiological resources that were depleted during the extra two days of starvation.

The estimated growth rates of females are higher than those of males in both 3 mg food input treatments, but lower than those of males in both lower food input treatments. Males and females grow more slowly on the day 8 delay than the day 6 delay at both food input amounts. The additional 2 days of starvation in the day 8 delay treatments results in the lower growth rates; males take longer to reach the same size (at 3 mg food input) while females grow for longer, but still pupate at a smaller mass than at the 6 day delay. This suggests that both males and females are improving their depleted physiological state at the same time as they are adding mass in the day 8 delay treatments.

Food 1 x delay x sex. Food input amount (1 mg vs 2 mg) appears to be more important to the mass at pupation than sex and also more important than the delay treatment; in the prior interaction, food input amount (1 mg + 2 mg vs 3 mg) is less important than either sex or delay. Food is probably limiting the growth of larvae in the test tubes with the 1 mg second food input, and may be limiting the growth of larvae in the test tubes with the 2 mg second food input.

The delay treatment has little effect on the mass of either males or females at the 1 mg food input. The delay treatment has a larger effect at the 2 mg food input, and the effect on males and females is opposite. Males pupate at the largest mass with the 2 mg food input and the day 8 delay, while females pupate at the largest mass with the 2 mg food input and the day 6 delay. The males in the 2 mg food input, day 8 delay test tubes pupate at the same size as the males in the 3 mg food input test tubes.

They apparently reach the maximum size as determined by the environment. Since these males receive a different second food input than the 3 mg food input males, the maximum size of males must have been set before the second food input, so before day 6. Because the males in the 2 mg food input, day 8 delay treatment grow as large as males in both 3 mg food input treatments, males are probably not limited by food at the 2 mg food input. Females grow larger on the 3 mg second food input than on the 2 mg food input, so they probably are limited by food at the 2 mg food input.

The estimated growth rate indicates that males and females respond differently to the effects of starvation (delay) and the amount of food in the second input. The difference in growth rate across the two delay treatments for both sexes is due to the age at pupation (not significant in this interaction). The growth rates support the idea that the larvae are rebuilding physiological reserves that do not show up as mass in the treatments with the 2 mg food input and the day 8 delay.

Combining results from both food 2 x delay x sex and food 1 x delay x sex. Males increase in size in response to the second food input; they reach a maximum size of about 2.27 mg in this experiment and then pupate. Males take longer to pupate on the day 8 delay than the day 6 delay. The additional two days of starvation causes males to defer pupation in order to grow larger, and possibly to improve their physiological state because the extra time spent feeding does not show up in mass.

Females increase in size in response to the second food input; each additional 1 mg increment of food results in larger masses at pupation. Each incremental 1 mg of yeast is less valuable than the previous one; as the amount of the second food input increases, the effect of the incremental food on the pupal mass is smaller.

The effect of the delay also changes as the food input increases. At the 1 mg second food input, there is little effect of delay on the pupal mass of females. At the 2 mg and 3 mg food inputs, the day 8 delay treatment females are 0.24 mg to 0.25 mg smaller than the day 6 treatment females.

There is also an effect of delay and food on age at pupation. The day 6 delay treatment females pupate earlier as the size of the second food input increases. The incremental food reduces the age at pupation as it increases the mass at pupation, and the effect of the 1 mg of food on both mass and age is less with each additional increment. In contrast, the age at pupation for the day 8 delay treatment females doesn’t change much in response to the incremental additions of food.

The additional two days of starvation change the way that females respond to additional food. They still grow larger on the extra food, but they don’t grow as large and they seem to grow for an amount of time that is not affected by the size of the food input. Females may be replenishing physiological resources that were used during the extra two days of starvation (day 8 delay), or they may have some maximum larval period after the food input, or both.

Food 1 x delay. This interaction shows the residual effect of the size of the second food input and the delay on the growth of larva regardless of sex. The amount of the second food input is more important to the mass at pupation than the delay in this interaction. The mass at the 1 mg food input is much smaller than expected while the mass at the 2 mg food input is only a little larger than expected, and the difference due to the delay is larger at the 2 mg food input. The second food input is more important than the delay for age at pupation as well as mass at pupation; the larvae pupate earlier on the 2 mg food input than the 1 mg input, and earlier with the day 6 delay than the day 8 delay. The second food input increases the mass and decreases the age at pupation, but the delay treatment increases both the mass and the age.

Food 2 x delay. This interaction between food 2 and delay compares the two lower second food inputs (1 mg and 2 mg) with the highest second food input (3 mg) crossed with the two delay treatments. The size of the second food input is also more important than the delay in this interaction. The lower food inputs grow larger on the day 8 delay (see above) but the 3 mg food inputs grow larger on the day 6 delay. The age at pupation is not significantly affected by this interaction; the larvae in the 3 mg food input and day 6 delay pupate much earlier than the other treatments, while those in the 3 mg food input and day 8 delay pupate latest.

Combining the results from both food 1 x delay and food 2 x delay. These interactions describe the growth of the larvae independently of their sex. Each increment of food (1 mg dry weight of yeast) results in larger pupae, but each additional increment increases the size of the pupae by a smaller amount. The age at pupation is only significant for the food 1 contrast, the two lower food inputs. The age at pupation drops much more with the 2 mg second food input in the day 6 delay treatment, so the effect of the additional food on age at pupation changes depending on the period of starvation (delay treatment). The age at pupation decreases again in the 3 mg second food input with the day 6 delay treatment, but increases slightly in the 3 mg food input with the day 8 delay.

The larvae appear to be using the food for something other than added mass in the day 8 delay treatments. It is likely that the early growth and molts of the larvae determine the largest size of the larvae, but that is similar for both the day 6 delay treatment and the day 8 delay treatment in this experiment. This suggests that the larvae are using some of the food and extra feeding time to improve their physiological condition rather than building mass in the day 8 delay treatments. Perhaps the larvae (both sexes) use the first milligram of food primarily for growth in mass, and the second milligram allows them to grow larger and faster, but the third milligram allows them to replenish the physiological resources that were depleted during the extra two days of starvation.

Food 1 x sex and food 2 x sex. These interactions are significant for age at pupation. Males pupate earlier than females at all three food inputs. Males pupate earlier than expected and females pupate later than expected (from the main effects). The age at pupation for males differs from the expected value by a similar amount for all three food inputs. The age at pupation for females differs from the expected value by a smaller amount as the food input increases. Males and females deviate from the expected values in different ways.

Males grow faster than females except at the 3 mg food input (estimated growth rate). The males in the 2 mg food input grow faster than the males at the 1 mg food input. The males in the 3 mg food input grow at the same rate as the males in the 2 mg food input. Males do not appear to be food limited in either the 2 mg or the 3 mg second food inputs.

Females grow faster, larger and pupate earlier with each increment of food. They grow more slowly than males at the 1 mg food input, at almost the same rate at the 2 mg food input, but faster than males at the 3 mg food input. This could be due to the apparent maximum size of males (2.27 mg in this experiment) or to exponential growth processes (the larger size of females) or both. Females appear to grow in response to the amount of food in the second food input; they may still be food limited at the 3 mg food input whereas the males appear to grow equally fast in both the 2 mg and the 3 mg food input, suggesting that even the 2 mg food input is more than sufficient food for them.

Food, starvation and sex. Male and female mosquito larvae respond to food inputs after periods of starvation in different ways. Males appear to grow to a size determined before day 6 of the larval period in response to the addition of food at the end of a period of starvation. The size of the food input, and the period of starvation (delay) affect both the mass and age at pupation. At the shorter period of starvation (the day 6 delay treatment), males grow larger with each increment of food and pupate earlier. At the longer period of starvation, males take longer to pupate than at the shorter period of starvation and the relationship between pupal mass and incremental food is obscured because even at the 2 mg food input, males grow to the 2.27 mg maximum size for this experiment. Males do not appear to be food limited at either the 2 mg or 3 mg food inputs.

Females appear to grow larger in response to the amount of the second food input. Each additional increment of food contributes less to the mass at pupation, and the longer period of starvation further reduces the contribution of each increment to the pupal mass. The age at pupation of females decreases as the food input amount increases for the shorter period of starvation. For the shorter period of starvation each incremental increase in the food input increases mass and decreases the age at pupation. For the longer period of starvation (the day 8 delay) females grow larger on each increment of added food, but not as large as in the comparable shorter period treatment, and they take longer to grow that large. Females may have a maximum larval period that causes them to pupate, resulting in smaller pupae despite equivalent amounts of food. Females may be food limited at the 2 mg food input and perhaps even at the 3 mg food input.

The larvae may be using the food inputs differently in the shorter and longer starvation treatments. Both sexes may be rebuilding physiological reserves that were depleted during the additional 2 days of starvation. Male and female larvae raised in isolation respond differently to identical conditions. Each increment of added food is less valuable than the previous increment as measured by mass at pupation. The value of food increments differs across sexes. Males and females alter the age at pupation differently in response to both food input amount and the period of starvation. Males appear to have a maximum size at pupation that is determined before day 6. Females appear to have a maximum duration of larval period that is affected by the timing of the second food input. Male and female larvae may delay pupation to rebuild physiological reserves at higher levels of the second food input. This delay appears to differ across sexes.

## Conclusions

These three experiments provide insight into how the environment affects 4 aspects of mosquito larval life: competition among mosquito larvae, the survival and growth of the larvae, and the triggers that initiate pupation. Competition, growth and the pupation triggers differ across the sexes; this suggests that survival may also differ across the sexes, but because the larvae that died were not identified by sex, this is not known. Competition also has an effect on survival; survival decreases in the treatments with the most competition.

### Survival across the three experiments

The levels of the independent factors were selected so that survival would be high. Survival is high in the first and the third experiments. In the second experiment, the much lower survival at the lowest density (1 larva per vial) prevents the analysis of the data and suggests that larvae facilitate the feeding of their peers. Also in the second experiment, the second lowest density (2 larvae per vial) appears to have reduced survival, but the experiment was not designed to test the effect of density on survival, so no statistical analyses were performed.

Competition in the first experiment has a small effect on survival (FxDxT accounts for 5% of the variance), but the distribution of the food in time (timespan treatment, FxT and DxT interactions) accounts for 10% of the variance. Survival is better in the treatments where the higher amounts of food are added over the longer timespan. Survival may be negatively affected by higher amounts of food at the shorter timespan. Too little food reduces survival, but too much food too early in the larval period also reduces survival. (see also [1] [39])

The aliquot treatment in the first experiment also affects survival; larvae survive better on 2 aliquots than on 4 aliquots (12% of the variance). This is opposite of the effect of the aliquot treatment on mass and age at pupation (larvae grow better on 4 aliquots than on 2 aliquots). There are no interactions between aliquot and the other treatments for survival; the effect of aliquot is independent of the interactions described previously. The 4 aliquot treatment provides more food early in the larval period than the 2 aliquot treatment, so this result is symmetrical to the observations above that too much food too early in the larval period reduces survival, but the effect of the number of aliquots is independent of food, density, competition and timespan.

Overall, the survival observations suggest that mosquito larvae are adapted to low levels of food and conditions of starvation, and that these experiments provide food levels that may be at the high end of those encountered in nature.

### Competition among females

The first experiment revealed that competition among females affects both mass and age at pupation, and is affected by both timespan and to a lesser extent aliquot, and that non-Prime females experience a release from competition with the Prime females at the highest food level with a large input of food on the third day.

In the most significant interaction describing competition, FxDxT, Prime females at high levels of food/larva grow according to the amount of total food in the test tube, while Average females grow according to the food/larva. Prime females dominate competition and the food supply. Because the Average female mass includes the Prime female mass, as the Prime female does better compared to the Average, the non-Prime females actually fare worse relative to the Prime female. The most intense competition between Prime and non-Prime females appears to be in the high food, high density, 3 day timespan treatment where the difference between the Prime and Average female mass is greatest, and the Prime female has the largest numerical advantage over the non-Prime females. At lower levels of food/larva, the Prime female changes feeding behavior and retains food for longer times. The difference between Prime female and Average female is smaller because the nature of competition changes from active filtering, where the larger Prime female can process more food faster than the smaller females, to retention governed by the absorption rate and/or the surface area of the gut, where the advantage of the Prime female over the smaller females is not as great.

The Prime female age at pupation follows the total food in the test tube at all levels of food/larva after day 4. Age and mass at pupation are affected differently by competition and the factors: food, density and timespan.

Aliquot interacts with food and density similarly to timespan, although the explained variance is much smaller. At high levels of food/larva in this interaction, the total food and the food/larva are in the same order, so Prime females and Average females both grow according food/larva at all food levels (with the total food affecting the outcome when the food/larva levels are equal). Within each food level, the females in the 4 aliquot treatment are larger and the difference between the Prime and Average females is smaller compared to the 2 aliquot treatment. There is intense competition between Prime and non-Prime females in the high food, high density, 2 aliquot treatment where the difference between the Prime and Average female mass is greatest, and the Prime female has the largest numerical advantage over the non-Prime females. The size of both Prime and Average females is much smaller in both most competition treatments compared to the other 6 treatments. Although the effect of the FxDxA interaction is small, it is similar to the FxDxT interaction. The addition of the third aliquot of food reduces competition for all females independently of the competition in the FxDxT interaction. The Prime female age at pupation is not affected by the FxDxA interaction.

After the effects of the FxDxT and FxDxA interactions are removed, there is a residual effect of competition on the Prime female age and the Average female mass. Because the Prime female mass is not significantly affected by this interaction, the residual effect is on the non-Prime females. These are disproportionately smaller in the most competition treatment than in the other treatments, and they are also disproportionately reduced in size relative to the Prime female in the high food, high density intermediate competition treatment. These are the same effects as seen in the two higher order interactions above, but there is a residual effect only on the non-Prime females. This suggests that the non-Prime females are more affected by competition for food than the Prime females.

There is also a residual effect on the Prime female age at pupation; this increases disproportionately with increasing competition after the higher order interactions have been removed. The mass at pupation of the Prime female is determined by the higher order interactions, and by the main effects, but the age at pupation is increased by increased levels of competition in this residual interaction.

### Competition among males

The first experiment reveals that competition among males affects both mass and age at pupation, and is affected by timespan. In the most significant interaction describing competition, FxDxT, Prime males at high levels of food/larva grow according to the amount of total food in the test tube; Prime males at intermediate and low levels of food/larva grow according to the food/larva level, but Average males do not follow either food/larva or total food. Prime males dominate competition and the food supply among males. Because the Average male mass includes the Prime male mass, as the Prime male does better compared to the Average, the non-Prime males actually fare worse relative to the Prime male. The most intense competition between Prime and non-Prime males appears to be in the high food, high density, 3 day timespan treatment where the difference between the Prime and Average male mass is greatest, and the Prime male has the largest numerical advantage over the non-Prime males. In the most competition treatment, both Prime and Average males are the smallest, but the Average male is larger than the Prime male, indicating that the non-Prime males grow larger after the Prime male pupates and on the additional food in the last food input at the end of day 6. The effect of this interaction on the non-Prime males suggests that food/larva, competition among females, release from competition after the pupation of the Prime male, and additional food on day 6 all contribute to the outcome of competition as measured in the mass at pupation. Competition among males differs from that among females and males appear to compete more intensely with other males than with females. The Prime male age at pupation is not in the same order as either the total food in the test tube, the food/larva, or the Average male mass. 7 of the 8 treatments pupate early (between 5.0 and 5.12 days), but the Prime males in the most competition, 6 day timespan treatment pupate later, after 5.70 days. In two cases, Prime males with greater access to food delayed pupation slightly perhaps to increase in size or improve physiological status. For males, age and mass at pupation are affected differently by competition and the factors: food, density and timespan. Males pupate earlier than females, and at a smaller size, and extend their larval life in response to too little food (most competition), and also when food is relatively abundant.

The FxDxA interaction has no effect on competition among males, in contrast to females. After the effects of the FxDxT interaction are removed, there is a residual effect of competition on the Prime male age and the Average male mass. Because the Prime male mass is not significantly affected by this interaction, the residual effect is on the non-Prime males. These are disproportionately smaller in the most competition treatment than in the other treatments, and they are also disproportionately reduced in size relative to the Prime male in the high food, high density intermediate competition treatment. In the most competition treatment, the Average male mass is again larger than the Prime male mass. In this case, there is no timespan treatment, so the difference in size reflects only the release from competition, not the addition of the final input of food on day 6. These are the same effects as seen in the higher order interaction, FxDxT, but there is a residual effect on only the non-Prime males. As with female larvae, non-Prime males are more affected by competition than Prime males.

There is also a residual effect on the Prime male age at pupation; this increases disproportionately with increasing competition after the higher order interactions have been removed. The mass at pupation of the Prime male is determined by the higher order interactions, and by the main effects, but the timing of pupation is increased by increased levels of competition in this residual interaction.

There are two other interactions that appear to affect competition among males, DxAxT and DxT. The Average male mass appears to be affected by the DxAxT interaction in the same way that it is affected by the FxDxT interaction. In this case, the size of the Average males corresponds to the estimated growth rate of the Prime male (the Prime male age and mass are not significantly affected by this interaction, but the growth rate is an indication of the competitive stress on the Prime male). The Average male grows larger than the Prime male in the treatment with the most competition (lowest growth rate). The non-Prime males grow larger than the Prime male due to the release from competition after the Prime male pupates, and on the additional food after the final input on day 6. The differences between the Prime male and the Average male are largest in the two treatments with high density and the 3 day timespan (both 2 and 4 aliquots); these are both intermediate food/larva treatments with high total food. This interaction is analogous to the FxDxT interaction with aliquot replacing food level. Aliquot changes the competition for non-Prime males independently of the interaction with food level. Other factors affect the Prime male mass, including competition (FxDxT), but the treatments that have aliquots of food added around day 3 allow the non-Prime males to grow larger. The timing of food inputs between days 2-4 appears to enhance the growth of the non-Prime males. This appears to be another kind of release from competition similar to the effect on the non-Prime females in the FxDxAxT interaction.

Analogous to the FxD interaction, there is a residual DxT interaction after the effects of FxDxT and DxAxT are removed. For all males, this appears to describe competition similarly to the FxD competition; the interaction is significant for the Prime male mass and age and the Average male mass. The two mass variables are in the same order as the food/larva in the test tubes. The Prime male does relatively better at the intermediate food level with more total food (high density, 3 day timespan), and the Average male mass is relatively smaller, indicating more competition in this treatment than in the two low density treatments. The Prime and Average males are smallest at the lowest food/larva level, and the Average male mass is larger than the Prime male mass, indicating a release from competition for the non-Prime males after the Prime male pupates and additional growth due to the final input of food on day 6. After the effects of competition in FxDxT and DxAxT are removed, there are still residual effects on the Prime and Average male masses that appear to be due to competition. The timespan treatment interacts with density analogously to the FxD interaction, but independently. The timespan treatment changes the amount of food in the test tube over time, and affects the food/larva and competition among males after the effects of FxDxT and DxAxT have been removed. This suggests that male larvae filter actively throughout their larval period and that each input of food changes the competitive environment for males.

The Prime male delays pupation in the two treatments with the most total food, and the treatment with the least food/larva and most competition. This is consistent with observations in other interactions, especially FxDxT.

### Competition between males and females

The first experiment shows that males compete differently from females and are affected differently by the factors, aliquot and timespan. Females actively filter particles at high food levels and switch to retention at lower food levels, changing the nature of competition among females. Males appear to filter particles without switching to retention at low food levels, so they are disproportionately affected by the particle level when females switch to retention. Males compete more intensely among themselves and may be competing for a subset of the particles, perhaps because of their smaller size. When competition is most intense among female larvae, the Prime male does better relative to the Prime female, but the non-Prime males do worse. Non-Prime males experience a release from competition after the Prime male pupates and also grow larger on the final input of food in the 6 day timespan treatments. The effect of the extra food is many times larger than the effect of the release from competition (0.14 mg for food plus release in FxDxT versus 0.01 mg for just the release in FxD).

The age at pupation is affected differently by competition than the mass at pupation in both sexes, and the effect on age at pupation is different between the sexes. The Prime female age follows the total food in the test tube while the Prime male age at pupation does not. The Prime male age is earlier, more simultaneous, and the small deviations appear to be due to food scarcity or food abundance. There is a residual effect of competition on age at pupation for both sexes after the higher order interactions have been removed; increased competition delays pupation for both the Prime female and the Prime male without affecting either mass.

There are four interactions that are expected to describe competition (the FxD set of interactions), and they each describe competition among female larvae, but only FxDxT and FxD are significant for male larvae. Furthermore there are two interactions that describe competition among males, but not females (DxAxT and DxT). FxDxT and FxD identify differences between competition among males and females summarized in the previous paragraphs. The interactions that describe competition in only one sex should provide additional insights.

In the FxDxAxT 4-way interaction, only the non-Prime females at the highest food level experience a release from competition (with the Prime female) when there is a large input of food on day 3 (high food, low density, 2 aliquot, 3 day timespan treatment). The non-Prime females grow larger on this abundant food, but the Prime female, the Prime male and the Average male masses are not affected. The Prime females in this treatment already experience an unlimited food supply, so more food does not increase their masses. This is probably true for the Prime males as well; both Prime male mass and Prime female mass follow the total food after day 4 and this treatment has the highest total food across the experiment (32 mg and 8 mg/larva). The non-Prime males are competing with the Prime male for food and extra food should increase their mass, but there is no effect of this interaction on Average male mass. This suggests three possibilities: the size of the non-Prime males is already determined before day 3 and the additional food does not affect it; the additional food is monopolized by the non-Prime females so there is none for the non-Prime males; there is already so much food for the non-Prime males as well as the Prime male and Prime female that the additional food has no effect on any of them. Both Prime and non-Prime males grow larger in response to additional food after day 3, so the suggestion that the size of males is determined before day 3 is not supported by evidence in other interactions. Other interactions suggest that increased food benefits all larvae, so the third possibility seems likeliest.

In the FxDxA interaction only the females experience competition. Both Prime and Average female masses follow the food/larva with the total food affecting the mass when the food/larva is equal across treatments. The 4 aliquot treatment results in more food by day 4 than the 2 aliquot treatments and this leads to larger females and smaller differences between Prime and Average females in the 4 aliquot treatments. The additional food due to the third aliquot reduces competition among females, but has no effect on the competition among males or on the competition between males and females. This suggests three possibilities similar to the ones in the FxDxAxT interaction above: the size of both Prime and non-Prime males is determined before day 3 and the additional food has no effect on it; the additional food is monopolized by the females so there is none for the males; there is already so much food for the males that the additional food has no effect on them. Both Prime and non-Prime males grow larger in response to additional food after day 3, so the suggestion that the size of males is determined before day 3 is not supported by evidence in other interactions. Other interactions suggest that increased food benefits all larvae, so the third possibility seems likeliest. This last possibility indicates that males and females are competing for different subsets of the food resource, possibly based on the size of the particles.

In the DxAxT interaction only the non-Prime males experience competition. The Prime males are affected by other interactions and the main effects, but are not affected by this interaction. The estimated growth rate for the Prime male, a proxy for the competitive stress on the Prime male, is the best indicator of the mass of the Average male. When the estimated growth rate is high, the competitive stress is low, and the non-Prime males are large and the difference between the Prime and Average male mass is small. At the two intermediate growth rates at high density, the non-Prime males are much smaller than the Prime males, indicating more competition in these treatments. At the lowest growth rate, the non-Prime males are the smallest, but they grow larger than the Prime males in these test tubes. Density, aliquot and timespan affect the food/larva and total food as the inputs are added over the course of the experiment, but the effect on the mass of the non-Prime males is mediated through competition with the Prime males. Males compete more intensely with each other than with females. The Prime female mass is also significantly affected by this interaction, but the effect appears to be on growth rather than competition among females. This interaction further suggests that males and females are competing for different subsets of the food resource.

In the DxT interaction, the Prime male and the Average male compete similarly to the FxD interaction. The timespan treatment affects the food environment independently of the food level or number of aliquots and changes the competitive environment among males, but not among females. At the high density, the 6 day timespan has a greater effect on competition among males, reducing the size of Prime and Average males and delaying the age at pupation. Females appear to be affected by the food/larva (mass variables) and the total food (age at pupation), but these effects appear to be due to exponential growth processes rather than competition. This suggests that males and females compete differently for different subsets of the food resource and also that males respond to the timing of food inputs differently from females. Females at low food/larva levels retain particles rather than actively filtering, and they extend their larval life until there is sufficient total food (16 mg for 8 larvae) or food/larva (2 mg food/larva) for them to pupate. Males actively filter at all food/larva levels and Prime males pupate later at low food levels, but still before the final input on day 6. Non-Prime males grow larger than Prime males on the release from competition and on the additional food on day 6, but the increment in this residual interaction is on the order of that in the FxD interaction (the release from competition). The mechanisms of competition are the same as in previous interactions, but this suggests that males are more responsive to inputs than females, perhaps because it takes longer for females to switch from retention to actively filtering, or perhaps because they require a larger number of particles to initiate the switch.

These four interactions that represent competition among females independently of males (FxDxAxT and FxDxA) and competition among males independently of females (DxAxT and DxT) confirm the conclusions in the first two interactions (FxDxT and FxD). Males and females experience different levels of food in the same test tubes and compete for different subsets of the food resource, probably based on the size of the particles. Males compete more intensely with males and females compete more intensely with females. Males respond to food inputs more rapidly than females, either because they are already actively filtering or because the females need larger inputs to switch from retention back to active filtering. Non-Prime males grow larger than Prime males due to a release from competition and also in response to additional food after the Prime males pupate. Females grow larger on the late addition of food, but the size distribution of females appears determined earlier in the larval period, and non-Prime females do not appear to experience a release from competition with the Prime female on the day 6 food input. Males appear to defer pupation in response to an abundance of food and also in response to low food levels. Females require more food than males and defer pupation only in response to low food levels.

Overall, food and density are the most important determinants of the outcome of competition, and both total food and food/larva affect the outcome. The size distributions of the male and female larvae are altered by competition. Non-Prime male larvae experience a small release from competition after the Prime male pupates and also grow larger on the food added on day 6. Females are not affected by the pupation of the Prime male, but do grow larger on the food added on day 6, however this food does not appear to alter the size distribution of females as it does with males. Non-Prime females experience a release from competition with the Prime female only at the highest food level with the largest food input on day 3.

Competition among males, among females and between males and females can be explained by active filtering and retention among females, active filtering among males, exponential growth, and delaying pupation in response to low levels of food and high levels of food, along with different life history strategies (triggers for pupation). There is no evidence for interference competition among these mosquito larvae.

### Growth versus competition

The interaction of food and density produces 4 competitive environments in the first experiment and there are four interactions that involve competition, plus two that similarly affect male larvae. There are also 4 interactions that do not include density, so only affect the food level and therefore the growth of larvae: FxAxT, FxA, FxT, and AxT. There are 3 other interactions that include density, but not the main factor of food level. Two of these describe competition among males and growth among females (DxAxT, and DxT) and the third one, DxA, is only significant for Prime female age at pupation.

The second experiment also investigated competition and growth, but due to differential mortality no statistical analyses were made.

The third experiment investigated growth after a period of starvation (equivalent to the 6 day timespan). Each test tube contained only one larva so no competition was possible in this experiment.

### The growth of female larvae

In the first experiment the highest order interaction describing growth, FxAxT, is only significant for the Prime female and Average female masses at pupation. The treatments separate into three groups based on the amount of food after day 4: high food level (above 4 mg food/larva); moderate food level (about 4 mg food/larva); and low food level (less than 4 mg food/larva), so food level is the most important factor in this interaction. At the highest food level the largest Prime females are in the 4 aliquot treatment, but the Prime females in the 2 aliquot treatment grow faster and pupate earlier, probably because of the large input of food on day 3. Average females also grow larger on 4 aliquots than on 2 aliquots. The treatment with 4 food inputs probably allows the females to continue actively filtering for longer. A large input on day 3 allows the Prime female to switch back to active filtering and grow larger and faster than the non-Prime females, so the timing of food inputs affects the growth of females.

At the moderate food level, the Prime females grow largest and fastest in the treatment that receives 16 mg of food on day 0, followed by the 4 aliquot treatment (4 mg per day on days 0-3) and then by the 2 aliquot treatment (8 mg on day 0 and on day 3). The non-Prime females grow largest on the 4 aliquot treatment, followed by the treatment that receives 16 mg of food on day 0, followed by the 2 aliquot treatment. Prime female age at pupation is not affected by this interaction and most of the Prime females in this group of treatment pupate on day 6 before the final aliquot. The treatment that receives 16 mg of food on day 0, receives another 16 mg at the end of day 6, but this does not appear to benefit the non-Prime females. Unlike the non-Prime males that grow larger than the Prime male, the difference between the Prime female mass and the Average female mass is largest in the treatment that receives 16 mg of food on day 0. This suggests that the size distribution of the females is determined by the time of the final larval molt and that females grow as large as possible based on the amount of food, but also as limited by the size of their head capsule, feeding apparatus or some other physical feature or physiological state.

At the low food level, there is insufficient food for the Prime females to pupate before the final input on day 6. Both Prime and non-Prime females grow larger in the 4 aliquot treatment than in the 2 aliquot treatment. Non-Prime females do relatively better in the 4 aliquot treatment compared to the Prime female. This indicates that early food inputs are more valuable than later food inputs and also that multiple inputs result in better growth than a large initial input followed by a large later input. The females may switch to retention earlier in the 2 aliquot treatment and this may affect both their final molt and the efficiency of converting food to mass. Retention results in smaller females and smaller size distributions, suggesting that it is less efficient than actively filtering. Despite receiving the same total amount of food as the 4 aliquot treatment, the females in the 2 aliquot treatment may be handicapped by the period of starvation so that the final molt does not allow them to grow as large as the females in the 4 aliquot treatment. The Prime females in the 2 aliquot treatment also take almost 2 days longer to pupate than those in the 4 aliquot treatments suggesting that mass alone is not the reason behind the timing of pupation after the period of starvation.

The FxT interaction represents the residual effect on female growth after considering the higher order interactions: FxDxAxT, FxDxT, and FxAxT. After removing the effects of competition and growth, what remains is an almost linear relationship between the total food after day 4 and the mass at pupation for both Prime and Average females. Each incremental increase in food results in about a 0.5 mg increase in the mass of both the Prime and Average female.

The AxT interaction represents the residual effect on female growth after considering the higher order interactions: FxDxAxT (competition), FxAxT (growth), and DxAxT (growth). The four treatments in this interaction change the food distribution pattern across all food levels and densities. Females grow best in the test tubes with 4 aliquots and the 3 day timespan; this translates to the most food early in the larval period. Females do disproportionately worse in the test tubes with 2 aliquots and the 6 day timespan; the Prime females pupate almost 2 days after the final input of food, and do not grow as large as other Prime females despite the same amount of food across all treatments. The difference between the Prime female and the Average female is also largest in this treatment; this is the opposite of the pattern expected for exponential growth. There is an effect of the pattern of inputs on the growth of female larvae regardless of food or density. The disproportionately small females with the largest difference between Prime and Average masses suggests that the Prime female dominates the non-Prime females during the early larval period when food is relatively abundant, followed by a period of retention during which the Prime female maintains and slowly increases this advantage, followed by a further period of active filtering during which the Prime female extends the advantage over the non-Prime females on the second food input after day 6. The period of starvation may also differentially affect the physiological condition of the non-Prime females, further contributing to the large difference between the Prime and Average females in this treatment.

The DxAxT interaction describes competition among male larvae, but growth among female larvae. Only the Prime female mass is affected by this interaction; the non-Prime females must be reduced in size collectively as the Prime female increases in size in order for the Average female mass not to be affected by this interaction. There are different effects at the higher and lower food levels (after day 4). At the highest food level, the Prime females in the low density, 2 aliquot, 3 day timespan treatment are larger than those in the low density, 4 aliquot, 3 day timespan; the main effects predict the opposite. The timing of the large second food input on day 3 accelerates the growth of the Prime female at the expense of the non-Prime females. In the low density, 4 aliquot, 3 day timespan treatment, the Prime females are smaller, but the difference between the Prime and Average females is also smaller; the 4 aliquot treatment favors the growth of the non-Prime females in this treatment.

In the 6 treatments at lower food levels, the mass of the Prime female follows the food/larva after day 4; where the food/larva is equal across different treatments, the total food determines the larger female, and in the two treatments where both food/larva and total food are equal, the treatment with 4 aliquots is larger than the treatment with 2 aliquots. However, in the high density, 2 aliquot, 6 day timespan treatment the Prime female mass is smallest and the difference between the Prime and Average female masses is greatest.

At both the highest food level and the lowest food level the Prime female benefits from the large second aliquot relative to the non-Prime females. This is related to density, but not competition. At the low density, the Prime female benefits over the non-Prime females from the large aliquot on day 3 (across both food level treatments). At high density, the Prime female benefits over the non-Prime females from the large aliquot on day 6 (again across both food level treatments). Food/larva rather than density, aliquot and timespan appears to explain the size of the Prime female at intermediate food levels. This suggests an effect of density independent of competition interacting with the distribution of food inputs during the larval lifespan of females. The most likely explanation is that the Prime female switches from retention to active filtering with the introduction of the second aliquot in these two treatments and grows more rapidly than the non-Prime females. That the Average mass is unaffected suggests that the large input on day 3 is unevenly divided between the Prime and non-Prime females, but results in the same Average mass (for instance, across the 2 aliquot and 4 aliquot treatments at low density and the 3 day timespan). The longer period of starvation (6 days rather than 3 days) affects the distribution of sizes (and perhaps physiological states) among the female larvae so that when food is added on day 6 the Prime female is already at a greater advantage relative to the non-Prime females. The switch from retention to active filtering allows the Prime female to grow even larger relative to the non-Prime females. There is no comparable food level at the high density treatment to see whether the Average female mass would be the same across the two different aliquot treatments.

In other interactions the Prime female dominates competition based on the total food at the higher food/larva levels, and follows the food/larva at lower levels, altering the size distribution of the female larvae as a result. There is no evidence for competition in this interaction, but low density results in unequal growth at high food levels (2 aliquots, 3 day timespan versus 4 aliquots, 3 day timespan), and unequal growth at high density and the lowest food level (2 aliquots, 6 day timespan). The timing of the input on day 3 probably coincides with the peak growth of females, while the delay to day 6 results in 3^rd^ and 4^th^ instar larvae that are stunted and can not grow as large as females in test tubes with more food earlier in the larval life.

The DxA interaction is only significant for Prime female age at pupation. The age at pupation is inversely related to the food/larva after day 4, but the Prime female pupates disproportionately early in the low density, 2 aliquot treatment. Since timespan is not a factor in this interaction, the 3 day and 6 day timespans are equivalent. In contrast to interactions including timespan, the difference across the treatments appears to be the amount of food in the initial input. Prime females in the low density, 2 aliquot treatment have more food/larva on day 0 than the other treatments; this accelerates their growth and results in the early pupation relative to the expected values.

The DxT interaction represents the residual effect on Prime female mass and age and Average female mass after the higher order interaction effects are removed. These interactions include competition (FxDxT, FxDxAxT) and growth (DxAxT). The residual effect on male larvae appears to be competition, but the residual effect on females appears to affect growth. There is an almost linear correspondence between the food/larva after day 4 and the masses at pupation for both Prime and Average females. Despite the low food/larva levels in some treatments, the ongoing food inputs over the timespan appear to allow the females to actively filter particles. The incremental increases in food across these four treatments are not regular and the corresponding increases in Prime and Average female mass are also not regular, but the slope of the line through these points is about half that of the FxT residual.

The Prime female age at pupation in the DxT interaction follows the total food after day 4 rather than the food/larva after the effect of the FxDxT interaction is removed. This is similar to the age at pupation in the FxDxT interaction (competition), but the DxT interaction appears to describe exponential growth processes for the female mass variables. For the Prime female, age at pupation may be more affected by the food environment than by competition.

In the third experiment, individual larvae receive 1 mg of food on day 0 and then 1 mg, 2 mg, or 3 mg of food on day 6 or on day 8. Females do not pupate on less than 2 mg of food (total), which is consistent with the 2 mg food/larva necessary in the first experiment (16 mg total food for 8 larvae). Age and mass at pupation are both affected by the amount and timing of the second food input in this experiment.

Females grow larger in response to the amount of the second food input. Each additional increment of food contributes less to the mass at pupation, and the longer period of starvation further reduces the contribution of each increment to the pupal mass. For the shorter period of starvation each incremental increase in the food input increases mass and decreases the age at pupation. For the longer period of starvation (the day 8 delay) females grow larger on each increment of added food, but not as large as in the comparable day 6 delay treatment, and they take longer to grow that large. Females may have a maximum larval period that causes them to pupate, resulting in smaller pupae despite equivalent amounts of food. Female larvae in this experiment may be food limited at the 2 mg second food input and perhaps even at the 3 mg food input.

### Summary of the growth of female larva

The growth of female larvae in the first experiment is determined by the amount of food after day 4; this quantity is affected by all four factors in various interactions. FxT and DxT reveal that the amount of food after day 4 is linearly related to the size of Prime and Average females after the higher order interactions (both competition and growth) are removed. FxAxT and DxAxT indicate different effects of food on growth at different levels of food after day 4. At the highest food level in both interactions, the addition of a large amount of food on day 3 (2 aliquot treatment) accelerates the growth of the Prime female over the non-Prime females; the non-Prime females do better in the 4 aliquot treatment. At the lowest food level in both interactions, there is not enough food for females to pupate until after the final food input on day 6. Females grow larger on the 4 aliquot treatment than the 2 aliquot treatment because there is more food by day 4, but the large input on day 6 results in a larger difference between the Prime and Average female masses as a result of exponential growth processes and active filtering of the additional food particles. At the intermediate food levels, food/larva appears to explain the mass of females in the DxAxT interaction, but the timing of the food inputs affects both the size of the Prime female and the difference between the Prime and Average females in the FxAxT interaction. The largest Prime females and the largest difference between Prime and Average females occur in the test tubes that receive 16 mg of food on day 0. The next largest Prime females and the largest Average females (with the smallest difference between the Prime and Average females) occur in the test tubes with 4 aliquots of 4 mg of food on day 0 through day 3. The smallest Prime and Average females occur in the test tubes with 8 mg of food on day 0 and another 8 mg of food on day 3. The 16 mg food input on day 0 allows the females to grow exponentially to pupation resulting in the largest Prime and the greatest difference between Prime and Average females. The 4 aliquot distribution of those 16 mg of food results in the smallest difference between Prime and Average females and the largest Average females; this is the main effect of the 4 aliquot treatment. The 2 aliquot distribution of the 16 mg food results in the smallest Prime and Average females indicating that they switch from actively filtering to retention between day 0 and day 3 and then switch back again after the day 3 food input. Where the second input (of two) at the high food level results in acceleration of the growth of females, the same treatment at intermediate food levels results in some stunting.

The AxT interaction is the residual for these two factors after competition and growth have been removed. Early food inputs result in better growth and the 2 aliquot, 6 day timespan results in disproportionately smaller females across all food levels and densities.

Equal increments of food have different values to female growth depending on the level of food already present and the timing of the input during the larval lifespan. Early food inputs are more valuable to growth than later food inputs. Inputs during the day 2 to day 4 period at high food levels have a larger effect on growth than earlier or later inputs. Each input appears to cause females to switch from retention to actively filtering, or to continue actively filtering. Active filtering results in larger females and larger size distributions among females than retention. Incremental food inputs after a period of starvation (day 6 or day 8) result in larger females, but each additional increment contributes less to the mass of the females and the longer starvation period also reduces the value of each increment. Females probably have a maximum size at pupation, but that was not apparent in these experiments.

There is no residual main effect of density on female mass at pupation after the interactions with food, aliquot and timespan have been removed. Density operates on female mass either through competition (FxD set of interactions) or through food/larva (the non-FxD interactions describing growth).

Female age at pupation is affected by the amount of food (total food, initial food on day 0). The interactions affect the age at pupation differently than mass at pupation. Females extend their larval period until there is sufficient food for them to pupate (16 mg for 8 larvae in the first experiment, 2 mg food/larva in the third experiment). Females appear to have a maximum larval period that is affected by late food inputs after starvation (in the third experiment).

Both mass and age at pupation may be affected by the physiological state of the female larvae; this is needed to explain the differences observed at different food inputs after the two periods of starvation (in the third experiment).

### The growth of male larvae

Three interactions describe the growth of the male larvae: FxA, FxT, and AxT. FxA and AxT are the highest order interactions for males between these pairs of factors. FxT is the residual after the FxDxT interaction is removed.

In the FxA interaction, Prime males grow larger in the 4 aliquot treatment than in the 2 aliquot treatment and aliquot has a larger effect at the low food level than at the high food level. It appears that the third aliquot (day 2 to day 4) accelerates the growth of the Prime male. Although both Prime and Average male masses are smaller at the low food level, the Prime male in the 4 aliquot treatment is relatively large compared to both the Average male mass and the Prime female mass. This suggests that males feed on a different subset of the food resource, probably based on particle size, and that the third aliquot enables the Prime male to grow at the expense of the non-Prime males. This is the result of exponential growth processes rather than competition. The Average male masses are in the same order as the Prime male masses, so all males appear to benefit from the 4 aliquot treatment. The Average male mass in the low food, 2 aliquot treatment is the smallest across this interaction, but it is the same as the Prime male mass, so the non-Prime males grow as large as the Prime males (probably on the food added on day 6 in some of the test tubes). There is no release from competition in this interaction, but there is additional food in some test tubes. Despite the late input of additional food, the non-Prime males don’t grow as large as the non-Prime males in the low food, 4 aliquot treatment (equivalent food, delivered earlier in the larval life). This suggests that there is a minimum mass at pupation for males that is determined by environmental conditions around day 3. Male larvae grow until they reach that minimum, or grow larger than the minimum, even delaying pupation, in response to greater availability of food. One possible mechanism could be that food delivered earlier in the larval period results in larger 3^rd^ and 4^th^ instars and these larvae are able to grow larger than peers that did not receive the early food inputs. Additional food late in the larval life allows growth, but that growth is capped because of physiological or physical limitations (size of the feeding apparatus or other body parts).

Prime males delay their age at pupation in the 4 aliquot treatment relative to the 2 aliquot treatment at the same food level. Prime males take longer to pupate at the lower food level, and delay pupation in the 4 aliquot treatment disproportionately longer at the lower food level.

In the FxT interaction, the residual effect on female growth is an almost linear relationship between the total food after day 4 and the pupal mass of females. Prime male mass and Average male mass are also directly related to the total food in three of the four treatments (0.2 mg increments versus 0.5 mg increments for the females). The Prime male at the lowest total food is disproportionately small, but still pupates before the final input on day 6. The Average male mass in this treatment is larger than the Prime male mass, so the non-Prime males grow larger than the Prime male on the final input of food on day 6. Competition has been removed from this interaction (by FxDxT) so this represents only the effect of additional food on day 6. The reason behind the disproportionately small size for the males in the low food, 6 day timespan treatment is probably the retention of food particles by the females reducing the apparent food level for the males. The Prime male pupates before the addition of food on day 6; the non-Prime males and females grow larger on the day 6 food input, but do not grow as large as larvae that received the same amount of food earlier.

Prime males delay pupation at the highest total food and at the two lower total food levels. They grow for longer and reach a larger size when food is most available, and also delay pupation when food is less available, but they still pupate on day 5 or day 6.

In the AxT interaction, the residual effect on females indicates that more food early in the larval life increases the mass for both Prime and Average females. There are no higher order interactions for the Prime male mass and age, and only DxAxT for the Average male mass. Both Prime and Average males grow disproportionately largest in the 2 aliquot, 3 day timespan treatment and disproportionately smallest in the 2 aliquot, 6 day timespan treatment. The two 4 aliquot treatments are intermediate in size, but the 2 aliquot, 3 day timespan treatment should be smaller than the 4 aliquot, 3 day timespan treatment (based on main effects), so a large input of food on day 3 benefits all males, perhaps because females switch from retention back to active filtering on the food input. The 4 aliquot, 6 day timespan treatment has 3/4ths the food after day 4 that the two 3 day timespan treatments have, but the males grow almost as large as the males in the 3 day timespan treatments. The multiple food inputs in this treatment cause the females to switch back to active filtering and this benefits the growth of males by increasing the number of particles in the test tubes. The worst treatment for both males and females is the 2 aliquot, 6 day timespan, where females retain food during most of the larval period and reduce the mass of males as well as that of females. After the food input on day 6, the females switch back to actively filtering; this allows the non-Prime males to grow larger on the day 6 food input and approach the mass of the Prime male in this treatment. The non-Prime males also appear to grow larger on the day 6 food input in the 4 aliquot treatment and approach the larger mass of those Prime males. The growth and mass at pupation of males in this interaction makes sense in the context of the growth and feeding behavior of the female larvae.

Prime males pupate earliest in the 4 aliquot, 3 day timespan treatment (in which females grow largest) and latest in the 4 aliquot, 6 day timespan treatment, probably due to the third aliquot food input on day 4. The two 2 aliquot treatments are intermediate between these and similar to their expected values. The Prime male pupates earlier than expected in the test tubes where the females grow largest, and fastest, and later than expected in the test tubes where additional food is added late in the larval period (day 4).

In the third experiment, the masses and ages at pupation of males are affected by the period of starvation (delay treatment) and the amount of food in the second food input, but both mass and age are affected differently than for females. At the shorter delay, equivalent to the 6 day timespan treatment in the first experiment, males grow larger and pupate earlier as the second food input increases. At the longer delay (8 days, 2 extra days of starvation), males grow larger than at the shorter delay, but take longer to pupate. Males at the longer delay and 2 mg second food input, and at both delays and the 3 mg second food input, pupate at 2.27 mg, which appears to be the maximum size of males in this experiment. Because this maximum does not seem dependent on the size of the second food input, it must be determined by environmental conditions before day 6.

### Summary of the growth of male larvae

The mass and age at pupation of males are more affected by competition than by growth; all interactions involving density appear to describe competition among males. FxT is the residual after competition has been removed; male mass appears to be directly related to the total food in the treatments with the exception of lowest total food. In the 3 treatments with the higher total food, each increment of food adds 0.2 mg to the mass of the Prime male and the Average male; this is only 40% of the increase that females realize, suggesting that males are using a subset of the food that females experience. The males in the low food, 6 day timespan treatment are disproportionately smaller than the males and females in other treatments and the females in this treatment. The non-Prime males in this treatment also grow larger than the Prime males on the additional food added on day 6. At the lowest total food, the feeding behavior of the female larvae constrains the growth of the male larvae independently of competition.

Males grow largest on 4 aliquots in the FxA interaction and the effect is more pronounced at the low food level. The extra food in the third aliquot appears to contribute to the growth of Prime and non-Prime males, especially at the low food level. The Prime male in this treatment is disproportionately large relative to the Average male and the Prime female. The difference between the Prime male and Average male is due to exponential growth processes, but the difference between the Prime male and the Prime female suggests that males are using a different subset of the food particles than females.

In the AxT interaction males grow better with 2 aliquots and the 3 day timespan than in the other treatments. The timing of the food input on day 3 accelerates the growth of males probably because the females switch from retention to actively filtering. The growth of females is similarly accelerated, but females in the 4 aliquot, 3 day timespan treatment are larger than those in the 2 aliquot, 3 day timespan. This also suggests that males are using a different subset of the food particles than the females. Males do least well in the treatment with 2 aliquots and the 6 day timespan. Females retain food during the period of starvation between day 0 and day 6 and all males are smaller. The non-Prime males in both the 6 day timespan treatments grow almost as large as the Prime males in those treatments on the food input on day 6.

The growth of males is affected by the availability of food particles, which is largely controlled by the female larvae. When female larvae experience low levels of food particles, they switch to retention, further reducing the level of food particles. Males appear to be using a different subset of particles than females. Non-Prime males also benefit from the late input of food on day 6 and grow almost as large as the Prime males in the same treatments, but not as large as non-Prime males in treatments that received the same amount of food earlier in the larval lifespan.

Males delay pupation to grow larger when food is most abundant (in test tubes where females grow largest and fastest). They also delay pupation to grow larger when food is scarce, especially in test tubes where females are retaining food. Prime males pupate on day 5 or day 6, but some non-Prime males delay pupation until after day 6 and grow larger on the final input of food (especially at the lowest food levels).

Both mass and age at pupation may be affected by the physiological state of the male larvae; this is needed to explain the differences observed at different food inputs after the two periods of starvation (in the third experiment).

### Differences in the growth of males and females

Food is the most important factor in the growth of mosquito larvae of both sexes, and the first experiment shows that the amount of food during the late larval period (3^rd^ and 4^th^ instars, days 2-4) is the most important determinant of the mass at pupation. Multiple food inputs during the early larval period benefit all larvae to some extent. Large inputs on day 0, and on day 3 can affect the ultimate mass at pupation and the size distribution of pupae by altering the feeding behavior of females (actively filtering versus retention), which affects the availability of particles to both female and male larvae.

The interactions describing the growth of female larvae partially overlap those describing the growth of male larvae. Four interactions describe only the growth of females: FxAxT, DxAxT, DxA, and DxT. Two interactions describe the growth of both males and females: FxT and AxT. FxA describes only the growth of males.

FxAxT, DxAxT and AxT show the effect of aliquot and timespan on the growth of females: an input of food early in the larval period contributes more to growth than the same amount of food later; inputs during the late instars accelerate growth; more, smaller inputs favor the growth of all female larvae, while fewer inputs alter the size distribution of females depending on the timing of the inputs. Aliquot and timespan affect the total food and food/larva in the test tubes; total food and food/larva affect the feeding behavior of females and this changes the availability of particles in the test tubes. This is the same mechanism that governs competition in some interactions, but results in different size distributions in the interactions describing growth.

Of these interactions, only the AxT interaction affects the growth of male larvae and indicates that the changes in feeding behavior of females affects the growth of males. In the AxT interaction males grow largest on 4 aliquots and the 3 day timespan, similar to the effect of the FxA interaction above, and also similar to the females in the AxT interaction. This appears to be due to the third aliquot of food on day 2. However, males grow almost as large in the 2 aliquot, 3 day timespan, suggesting that the large input on day 3 is almost as beneficial to males as the 4 aliquots delivered over 3 days.

Some non-Prime males in the AxT interaction do grow larger on the day 6 food input and the Average male mass approaches the Prime male mass in the 6 day timespan treatments. Despite receiving much more food on day 6, the non-Prime males do not grow as large as the Prime males in their respective treatments. This suggests two things mentioned previously: a food input early in the larval lifespan contributes more to growth than the same food input on day 6; and, males appear to grow to a minimum size and then extend their larval period in response to food abundance, taking longer to pupate and growing larger.

Two significant residual interactions, FxT and DxT, show a linear relationship between food and the pupal mass of females. Only the FxT interaction affects the growth of male larvae and indicates a similar linear relationship between food and the pupal mass of males, except for the lowest food level, where the males are disproportionately small, probably also because of the changes in the feeding behavior of females. Females increase in mass by 0.5 mg with each increment of additional food (on day 4) across the 4 treatments. Males increase by only 0.2 mg in the 3 high total food treatments. Males appear to feed on a reduced subset of the particles compared to females. Both males and females show linear relationships between the dry weight of yeast and the wet weight of pupae once competition and higher order growth interactions have been accounted for.

The aliquot treatment does not affect competition among Prime males, but it does affect the growth of Prime and Average males. The FxA interaction indicates that males grow disproportionately large at the low food level and the 4 aliquot treatment compared to the 2 aliquot treatment. Independent of density and timespan, the 4 aliquot treatment delivers more food early in the larval life than the 2 aliquot treatment. The difference is likely the third aliquot of food on day 2 or day 4. This is similar to the effect of the third aliquot on the growth of females in the FxAxT and AxT interactions. This suggests that the early food inputs are more important to the growth of males than later ones. Furthermore, the Prime male mass is disproportionately large at the low food, 4 aliquot treatment compared to both the Average male mass and the Prime female mass. The comparison against the Average male mass can be explained by exponential growth processes, but the comparison against the Prime female suggests that males compete for a different subset of the food supply than the females.

Some non-Prime males in the FxA interaction do grow larger on the day 6 food input and the Average male mass is the same as the Prime male mass at the low food level, 2 aliquot treatment. This suggests that there is a minimum mass at pupation for males that is determined by environmental conditions around day 4 (the day the third aliquot is added for half the timespan treatments). Despite receiving much more food on day 6, the non-Prime males do not grow as large as the non-Prime males in the 4 aliquot treatment at the same food level. Males grow until they reach this minimum mass, then grow larger if there is abundant food, delaying pupation in order to grow. Prime males pupate before day 6. Some non-Prime males grow larger on the additional food after day 6 (in some treatments), but the food added on day 6 is not as beneficial as food added earlier.

The amount of food added after a period of starvation affects the mass at pupation of females. The size of the female increases as the amount of food increases, but each increment of food contributes less to the pupal mass. The longer period of starvation also reduces the value of the food to the pupal mass, and also decreases the value of each subsequent increment. It appears that females use food inputs after a period of starvation to improve physiological status as well as to increase mass. The amount of food added after a period of starvation also affects the mass at pupation of males. Similar to female larvae, the size of the male increases as the amount of food increases, and each increment of food contributes less to the pupal mass. The longer period of starvation also reduces the value of the food to the pupal mass, and decreases the value of each subsequent increment. In contrast to the females, male larvae pupate at the same mass (2.27 mg) in 3 of the 6 treatments; there is a maximum size at pupation for males that is set before the second food input on day 6. Because not all the food in the second food input results in growth, it appears that males also use food inputs after a period of starvation to improve physiological status as well as to increase mass.

Density affects the growth of females by altering the food/larva or the total food in the test tube. There are no residual main effects of density on competition or growth among female larvae. Density affects competition among males, but not the growth of males. Unlike females, there are residual main effects of density on the masses of male pupae. Higher density reduces the pupal mass of all males.

The age at pupation of Prime females is altered by the interaction DxA. The exclusion of food level and timespan from this interaction suggests that density and aliquot have an effect separate from total food or food/larva. At low densities, both aliquot treatments pupate early, but at high densities, the 2 aliquot treatment is much later than the 4 aliquot treatment. The source of the interaction appears to be that the low density, 2 aliquot treatment pupates much earlier than projected by the main effects. This is probably due to the large initial input of food; the food/larva on day 0 for this treatment is 2 mg or 4 mg, depending on the food level, twice that of the next highest initial food levels. This only affects the timing of pupation. The Prime female grows to a size and at a rate commensurate with the main effects and other interactions, but pupates earlier than expected because of the large initial food input.

There is also a residual effect on the age at pupation of Prime females in the DxT interaction. Age is inversely related to the total food in the treatments; this is the same relationship as in the FxDxT interaction for which this is one of the residuals.

The age at pupation of Prime male larvae is affected by the interactions describing growth; these differ from the interactions that affect the age at pupation of Prime female larvae (DxA and DxT). For Prime males, the growth interactions FxA, FxT, and AxT, are significant for age at pupation. Prime males delay pupation when food is abundant, and also when food is scarce. This is the same response that Prime males exhibit in the competition interactions.

In the third experiment, some males pupate on day 7 or day 8 before the addition of the second food input. These males pupated on 1 mg dry weight of yeast and attained a wet mass of 1.06 mg (SD=0.22). These males are excluded from the MANOVA analysis, but provide an additional reference point to compare the wet mass of males and females against the total dry weight of yeast in the rest of the experiment. Both males and females increase in size as the total dry weight of yeast increases. Females are larger than males at all food levels and the difference between the female wet mass and the male wet mass increases as the total dry weight of yeast increases. The slope of the 6 female wet masses plotted against the dry weight of yeast is 0.76. The slope of the 7 male wet masses plotted against the dry weight of yeast is half that, 0.38. Females appear to grow larger than males on equal dry weights of yeast. (see S38 Fig). This is similar to the difference between males and females in the residual interaction FxT above.

### Summary of the growth of males and females

The change in size due to exponential growth processes looks like a sigmoid curve when plotted against time. For mosquito larvae each instar should resemble a sigmoid curve. Without having measured the growth trajectories of each instar, the data from these experiments is consistent with female larvae growing to the top of the sigmoid curve and male larvae molting and pupating at some lower level. This takes into account the data on growth rates, and masses and ages at pupation. Females grow larger because at each molt, they attain a larger size than the males in the cohort. They grow faster because they molt/pupate at a point higher on the sigmoid curve than the males, and the line between the origin for that instar and the point at which the molt/pupation occurs will have a greater slope (growth rate) for females than males, all other conditions being equal. Females take longer than males because they defer molting/pupation at each stage in order to grow larger. There is a positive feedback loop between females deferring the molts between instars in order to grow larger, and being able to feed faster in the next instar, thereby growing even faster than males.

At the end of each instar both males and females face alternatives that appear to be determined by the availability of food. If there is too much food, the larva dies. If there is a lot of food, the larva extends growth to continue to grow (probably to some physical or physiological maximum based on maximum size for the exoskeleton, or maximum feeding rate counterbalanced by the cost of maintaining the existing mass, or to some internal physiological state). If there is just the right amount of food, the larva molts or pupates. If there is less than the optimal amount of food, the larva defers molting or pupation to grow larger. If there is not enough food to molt to the next instar or to pupate, the larva enters diapause and waits for more food (and may die eventually). If there is too little food to enter diapause, the larva dies. Females have an additional option; they can change their feeding behavior from actively filtering to retaining particles. This changes the available food for other larvae in the same microcosm. It also appears to be less efficient at extracting nutrients from the food supply and results in slower growth and a smaller size distribution among female larvae. Males appear to have a minimum size at pupation that is determined by food early in the larval life (around day 3); once they reach this size, they may pupate or continue to feed and grow if food is available. Prime males appear to have a maximum size that is determined by the amount of food before day 6. The non-Prime males are able to grow larger than the Prime male in some circumstances when food is added to the test tubes on day 6 (after the Prime male has pupated). Females in these experiments did not reach a distinct maximum size; females required more food to pupate at all than males did and grew larger than males and for longer at all food levels.

### Pupal triggers

Determining the triggers that result in pupation for these larvae is not a primary goal of these experiments. Nevertheless, the measurement of age and mass at pupation for the Prime males and females results in observations that suggest pupal triggers. Males appear to have a minimum mass determined by the food environment above which pupation can occur. Males appear to have a maximum mass at which pupation occurs, also determined by the food environment early in the larval period. Males pupate at a smaller mass than females and on less food. Females do not show the same minimum mass and maximum mass effects as males. Females appear to grow to a size commensurate with the food level and require at least 2 mg food/larva in order to pupate.

In addition to the pupal triggers based on mass, there appear to be different triggers based on the age of the larvae. Males pupate earlier than females and for Prime males the distribution of ages is much tighter than for females. Non-Prime males sometimes extend their larval period to grow larger than the Prime male; this suggests an alternate life history strategy for males based on larger size and longevity rather than early emergence. Females appear to have a maximum larval period after starvation that is not affected by the amount of food added to end the period of starvation.

The age and mass triggers are not independent of each other; both Prime males and Prime females appear to extend the larval period at low food levels in order to grow larger. Prime males also appear to delay pupation at high food levels in order to grow larger.

In the absence of sufficient food to complete pupation, both males and females wait for additional food inputs. Both males and females pupate at a larger size and earlier as the amount of the late food input increases, but the length of the period of starvation affects the value of the food and the contribution to pupal mass. Males grow larger and take longer to pupate after the longer period of starvation compared to the shorter (6 day) period. Females do not grow as large after the longer period of starvation and appear to be limited to a specific interval of growth after the late food input. Males reach their maximum size and pupate, while females grow until they reach their maximum larval period and then pupate.

Both males and females appear to use some of the second food input to replenish internal physiological factors after a period of starvation; the pupal mass does not reflect the entire value of the late food input and this is more apparent after the longer period of starvation. This means that the triggering of pupation is dependent on at least: age or time, mass, environmental food conditions, and internal physiological status. Furthermore, the maximum size of pupae is probably also constrained by early life history due to the size of the head capsule, the feeding apparatus or some other physical or physiological feature determined by the molt to the 4^th^ instar.

Mosquito larvae could trigger larval molts and pupation based on interactions with other larvae in addition to the factors mentioned above. The mass at pupation of the Prime male is clearly affected by the sex ratio in the microcosm (see experiment 2 in [1]), which suggests the possibility of direct interactions between larvae. Females are not affected by the sex ratio in that experiment; this suggests that female larvae are not interacting with other larvae (male or female) and that they pupate only in response to their internal state and the environmental factors. This is supported by the lack of any residual effect of density on female mass. However, the ages at pupation for both Prime males and Prime females are significantly affected by the residual effect of density. This could indicate direct interactions among larvae affecting the timing of pupation but not the mass at pupation.

## Summary of conclusions

1. Male and female mosquitos compete for different subsets of the yeast cell food resource in microcosms. The growth of males and females suggests that females feed on a larger subset of the yeast cell food resource than males.
2. Males compete more intensely with males and females with females.
3. The amount and timing of food inputs alters both growth and competition for males and females, but the effects are different between sexes.
4. Increased density increases competition among males.
5. Among females, density operates primarily by changing the food/larva or total food per test tube; this affects competition in some interactions and growth in others.
6. There is an effect of density on the timing of pupation for females independent of competition or changes in food/larva or total food.
7. Food added earlier in the life span contributes more to mass than the same quantity added later.
8. After a period of starvation larvae appear to use some of the subsequent food input to rebuild internal physiological reserves in addition to building mass.
9. The timing of pupation is affected by the independent factors and competition, but not in the same way for the two sexes, and not in the same way as mass at pupation for the two sexes.
10. Male and female larvae have different life history strategies (see points 11-14 below).
11. Males grow quickly to a minimum size, then pupate depending on the amount of food available.
12. Males that do not grow quickly enough may delay pupation further to grow larger, resulting in a bimodal distribution of sizes and ages.
13. Males appear to have a maximum size determined by the food level before day 6.
14. Females grow faster than males and grow larger than males on the same food inputs. (See also point 1 above).
15. Females affect the growth and competition among males by manipulating the number of particles in the microcosm through changes in feeding behavior.
16. Mosquito larvae appear to have evolved to survive periods of starvation and take advantage of intermittent inputs of food into containers.

## Discussion

Competition is the result of multiple individual actions and interactions, but results in a specific distribution of larval sizes for each set of environmental conditions (in the first experiment: food level, density, aliquot and timespan). For females, at high relative food abundance, the Prime and the Average masses are large and there is a small difference between them. At low relative food abundance, the Prime and Average masses are small and there is a larger difference between them, but not the largest difference. At intermediate relative food abundance, high density favors the Prime female over the others; the Prime female mass is almost as large as at higher food abundances and the Average is similar to that at the low density. The controlling factors appear to be different aspects of the food resource (e.g. total food per test tube after day 4 and food/larva after day 4) as affected by the four factors above. The Prime female grows to a size determined by the total food (after day 4) in the test tubes where the food/larva (after day 4) is 4 mg or higher. Below that food/larva amount, the Prime female grows to a size determined by the food/larva. The age at pupation for the Prime female is determined by the total food at all levels of food/larva. The size at pupation of the Average females is determined by the food/larva at all levels of food/larva.

The relationships in the previous paragraph, taken together, mean that the Prime female outcompetes the non-Prime females for particles when they are most abundant and grows largest, fastest and pupates earliest. Because particles are abundant, the non-Prime females grow large also, but only as large as the food/larva indicates. At intermediate food levels (4 mg food/larva in the first experiment), the higher density results in larger Prime females because the total food is greater, but the Average size of females is the same as in the low density, intermediate competition treatment (also 4 mg food/larva). The high density, 4 mg food/larva results in the largest difference between the Prime and Average females. At lower levels of food/larva, the females switch from active filtering to retention and the sizes of the Prime and Average females are small, but the size distribution (the difference between the Prime and Average masses) is smaller than the high density, 4 mg food/larva test tubes.

The experiment crossed food level and density, which produce 4 different competitive environments, with aliquot and timespan, which alter the timing and amount of food across the experiment. Many Prime females pupate before the final aliquot is added for the 6 day timespan, so the amount of food after the last additions before that, on day 3 for the 3 day timespan treatments, and on day 4 for the 4 aliquot, 6 day timespan treatments, reflect the actual amount of food in the test tubes. Aliquot and timespan interact with FxD separately, with the exception of the 4-way interaction for the non-Prime females. When a large amount of food is added on day 3, the non-Prime females experience a release from competition and grow larger than expected; this causes a 4-way interaction in the ANOVA for the Average females. Both the 3-way interactions, FxDxT and FxDxA, are significant for Prime and Average female masses. FxD represents the residual effect of food level and density on these variables after the higher order interactions have been removed. The Prime female age and Average female mass are significant in this interaction. Competition extends the time to pupation and reduces the size of the non-Prime females after the effects of aliquot and timespan have been removed. Three other interactions involve density and one or more of the aliquot and timespan treatments; these are aspects of the abundance of food, and both aliquot and timespan interact with competition, so they could also indicate competition from aspects of the food availability independent of the factor, food level. If this were the case, the same pattern of distribution of sizes should appear. There is no evidence that these three interactions, DxAxT, DxA, or DxT, affect competition among females. They appear to describe the growth of females.

Competition among males also results in a pattern of size distributions. Prime males respond to total food and food/larva similarly to Prime females, but the Average males do not correspond to the Average females. Males appear to be competing for a different subset of the yeast particles than females; this is probably related to the different sizes of male and female larvae and their associated feeding structures. Males compete more intensely with other males and females with other females. Females do affect competition among males by reducing the low particle numbers even further due to a change in feeding behavior (retention). Non-Prime males experience a release from competition after the pupation of the Prime male, and also benefit from the day 6 addition of food. The release from competition results in a 0.01 mg increase in size (FxD), while the combined release and extra growth on the day 6 food results in a 0.14 mg increase (FxDxT). Unlike females, males appear to have two life history strategies: early pupation vs taking longer and growing larger. Early pupation may enhance mating success for males, while larger adults live longer and may have more opportunities to mate.

Unlike females, the DxAxT and DxT interactions appear to produce size distributions of males that resemble the FxDxT and FxD competitive outcomes. All of the significant interactions that include density as one of the factors result in outcomes that look like competition among males, even those that do not include total food. The factors that describe the distribution of food in time (aliquot, timespan) interact with density independently of the food level to alter the competitive environment for males, but not for females. Note that the 4-way interaction (FxDxAxT) is not significant for males in the MANOVA or the ANOVAs. This suggests that the mechanism of competition differs between males and females. Unlike females, the aliquot treatment does not interact with FxD to affect competition. This suggests that females switch from actively filtering to retention and back in response to the aliquot treatment, but males are unaffected because they are always actively filtering. There is some evidence that males respond quickly to small changes in the abundance of food, while females respond more slowly or not at all.

Competition results in one or a few winners and multiple losers in each container. The Prime individual (one male and one female in each microcosm) represent the winners and the Average individuals represent the losers. Intra specific competition among *A. aegypti* larvae is reviewed in [1]. Most of these studies do not take into account the concept of Prime individuals. Similarly, inter specific competition between *A. aegypti* larvae and other mosquito larvae is alleged to determine the abundance and distribution of the species in nature [3] [6] [11–13] [40–62]. None of these studies accounted for differential effects of competition on winners and losers in the microcosms. In the first experiment in this study, the Prime female and the Prime male are affected by the food, density and timespan treatments, but grow and pupate as though they had access to the total food in the microcosm. The non-Prime individuals, as represented in the Average male mass and Average female mass, experience negative effects of competition and these increase as the competition increases in intensity. Extrapolating to inter specific competitive contests, it is possible that the Prime individuals (of both species) are less affected or not affected by the competitive interactions that the average size and average age at pupation measure. Far from organizing communities as suggested in the introduction, larval competition may be largely irrelevant to the ecology and evolution of mosquitoes; this needs to be examined in further experiments.

The growth of larvae is described by interactions that don’t involve density for males, and for females, by interactions that don’t involve both food and density. Larvae are expected to grow in a sigmoid growth curve for each instar. The data suggest that males molt and pupate earlier on the growth curves than females, but both sexes alter the timing of the molts based on the availability of food. The first experiment suggests that early food inputs are better than late food inputs, and that timing the input during the 3^rd^ and 4^th^ instars results in larger pupae. The third experiment further extends the observation that early food inputs are better than late food inputs, but also indicates that the larvae replenish internal resources after a period of starvation. Not all the food in the late food inputs appears as mass at pupation, and less of the food ends up in mass at the day 8 delay than the day 6 delay (two days longer starvation).

The first experiment explains such a high percentage of the variance in mass and age at pupation for males and females that it is essentially a deterministic description of the responses of mosquito larvae to their food environment. Natural containers are larger and food inputs are less regular; both these alterations could reduce the deterministic aspect of the larval environment. However, it suggests that the choice of the larval environment is important to the success of each adult female’s progeny. To the extent possible ovipositing females should choose optimal containers. Furthermore, females should ideally provision their eggs to result in the largest possible first instar hatchlings. Opposing these forces are the possibility that any one container could be destroyed or drained or just barren of food, and that fewer, larger eggs mean a greater risk of complete reproductive failure in an unpredictable environment. Experiments on container choice, oviposition preference, and propagule size (egg size) and provisioning (egg quality) by females should indicate how deterministic the larval outcomes are in nature.

Females appear to prefer containers that host congeneric larvae, that are larger, and that are darker or shaded [28] [63–67] and that host other species’ larvae and pupae [44]. They may be attracted to chemical cues from bacteria in the containers [68–70]. This suggests that viable larvae, a larger container volume, and a protected container are the most important parameters that the ovipositing females search for. Propagule size (egg size) is directly related to female body size and blood meal size, which are also correlated with each other [34] [71, 72]. Some small females produce fewer, larger eggs than expected, suggesting that large eggs have an advantage over small eggs despite the greater risk of reproductive failure with fewer larger eggs. Provisioning (egg quality) does not appear to have been measured.

Mosquito larvae grow differently on different food sources and substrates [3] [6, 7] [42, 43] [45] [73–83]. In addition to the supplied food sources, usually yeast cells or bacteria, bacteria in the gut of the larvae may provide nutrients [84–87] and endoparasites may compete for nutrients [88]. The difference in feeding behavior between male and female larvae suggests that female mosquito larvae may harvest nutrients from retained yeast cells and/or bacteria while male larvae do not.

Ecologically, and evolutionarily, *A. aegypti* mosquitoes may have adapted to survive in impermanent aquatic habitats by growing quickly in response to food inputs, then resisting starvation until the next input, eventually reaching a size that permits pupation as a viable adult. Males and females appear to have different larval life history strategies probably due to their different reproductive roles. Adult males need to find virgin females [35] [89–93] to inseminate, while adult females need to be as large as possible to produce more eggs and larger eggs [4, 5] [34] [40, 41] [71, 72] [94–103], but see [96]. Size also increases longevity, which probably increases the likelihood of reproductive success. Males pupate earlier than females and at a smaller size in order to inseminate females of their own cohort, but some males aren’t able to pupate quickly and delay pupation to grow larger on later additions of food. This bi modal distribution of mass and age at pupation may result in reproductive success for the slower-growing males in subsequent cohorts.

It is also clear that there are physiological conditions that affect the age and mass at pupation [4] [10] [34] [94–95] [104–109]. These pupal triggers are also affected by the food environment in these experiments, but the relationship between the internal state and the external environment is not as clear as the relationship between competition and growth and the pupal mass and age.

## Acknowledgements

I dedicate this paper to the memory of Dr. Joanne M. Werts; she did not live to see the completed document. I would like to thank Dr. J. H. Frank for his advice and encouragement. Drs. W. Bradshaw, J. R. Linley, L. P. Lounibos, G. F. O’Meara, and J. M. Werts also read the manuscript and offered helpful suggestions. I am grateful to Dr. Werts and the Department of Zoology at Duke University for computer time. My late partner, Richard O. Merritt, provided posthumous support to allow me to complete the paper after many years delay. My husband, Dr. Douglas Kline, provided encouragement, editing and technical guidance in the completion of the task.

## Supporting information

S1 Dataset. Experiment 1. Excel file of the mass, age at pupation and sex of each individual pupa by treatment.

S2 Dataset. Experiment 2. Excel file of the mass, age at pupation and sex of each individual pupa by treatment.

S3 Dataset. Experiment 3. Excel file of the mass, age at pupation and sex of each individual pupa by treatment.

S1 Fig. Experiment 1. Scatterplot of mass at pupation (mg) in 0.2 mg increments versus the number of females at each mass.

S2 Fig. Experiment 1. Scatterplot of age at pupation (days) versus the number of females pupating on each day.

S3 Fig. Experiment 1. Scatterplot of mass at pupation (mg) in 0.2 mg increments versus the number of males at each mass.

S4 Fig. Experiment 1. Scatterplot of age at pupation (days) versus the number of males pupating on each day.

S5 Fig. Experiment 1. 3D visualization of Survival for FxDxT.

S6 Fig. Experiment 1. 3D visualization of Prime female mass for FxDxT.

S7 Fig. Experiment 1. 3D visualization of Prime female age for FxDxT.

S8 Fig. Experiment 1. 3D visualization of estimated growth rate for Prime females for FxDxT.

S9 Fig. Experiment 1. 3D visualization of Average female mass for FxDxT.

S10 Fig. Experiment 1. 3D visualization of Prime female mass MINUS Average female mass for FxDxT.

S11 Fig. Experiment 1. 3D visualization of Prime male mass for FxDxT.

S12 Fig. Experiment 1. 3D visualization of Prime male age for FxDxT.

S13 Fig. Experiment 1. 3D visualization of Average male mass for FxDxT.

S14 Fig. Experiment 1. 3D visualization of Prime male mass MINUS Average male mass for FxDxT.

S15 Fig. Experiment 1. 3D visualization of estimated growth rates for both Prime females and Prime males for FxDxT.

S16 Fig. Experiment 1. 3D visualization of Prime female mass MINUS Prime male mass for FxDxT.

S17 Fig. Experiment 1. 3D visualization of Prime female mass for FxDxA.

S18 Fig. Experiment 1. 3D visualization of Average female mass for FxDxA.

S19 Fig. Experiment 1. 3D visualization of Prime female mass MINUS Average female mass for FxDxA.

S20 Fig. Experiment 1. 3D visualization of estimated Prime female growth rate for FxDxA.

S21 Fig. Experiment 1. 3D visualization of Prime female age and Average female mass for FxD.

S22 Fig. Experiment 1. 3D visualization of Prime male age and Average male mass for FxD.

S23 Fig. Experiment 1. 3D visualization of Prime female mass for FxAxT.

S24 Fig. Experiment 1. 3D visualization of Average female mass for FxAxT.

S25 Fig. Experiment 1. 3D visualization of Prime male mass and age for FxA.

S26 Fig. Experiment 1. 3D visualization of Prime female mass and age for FxT.

S27 Fig. Experiment 1. 3D visualization of Prime male mass and age for FxT.

S28 Fig. Experiment 1. 3D visualization of mass versus total food after day 4 for FxT.

S29 Fig. Experiment 1. 3D visualization of Prime female mass and age for AxT.

S30 Fig. Experiment 1. 3D visualization of Prime male mass and age for AxT.

S31 Fig. Experiment 1. 3D visualization of Prime female mass for DxAxT.

S32 Fig. Experiment 1. 3D visualization of Average male mass for DxAxT.

S33 Fig. Experiment 1. 3D visualization of Prime female age at pupation for DxA.

S34 Fig. Experiment 1. 3D visualization of Prime female mass and age for DxT.

S35 Fig. Experiment 1. 3D visualization of Prime male mass and age for DxT.

S36 Fig. Experiment 1. 3D visualization of mass versus average food/larva after day 4 for DxT.

S37 Fig. Experiment 3. Heuristic 3D graph of male and female masses and ages at pupation for the 3-way interaction between second food input, delay and sex in the third experiment.

S38 Fig. Experiment 3. Graph of the wet weight (mg) of male and female pupae against the dry weight of the yeast food source.

S1 Table. Experiment 1. Food input schedule for each treatment according to the factors: food level (16 mg, 32 mg), aliquot (2 portions, 4 portions), and timespan (over days 0 to 3 or days 0 to 6). Density affects the food/larva which is also affected by the food input schedule.

S2 Table. Experiment 2. Treatment numbers with density (larvae/vial), food/larva (2 mg, 3 mg, 4 mg, and 5 mg dry weight of yeast), and number of replicates.

S3 Table. Experiment 3. Treatments showing the amount of the second food input (1 mg, 2 mg, 3 mg dry weight of yeast) and the delay (day of second food input, day 6 or day 8) with the number of replicates. All treatments received 1 mg dry weight of yeast on day 0 of the experiment.

S4 Table. Experiment 1. Significant correlations between composite scores and the 7 dependent variables with MANOVA significance levels and R squared by significant contrasts.

S5 Table. Experiment 1. Explanation of the meaning of the single df contrasts.

S6 Table. Experiment 1. ANOVA mean squares, (r squared), and significance [*] for each single df contrast across the 7 dependent variables.

S7 Table. Experiment 1. Means (SE) for the main effects: food, density, aliquot and timespan; for the 7 dependent variables.

S8 Table. Experiment 1. Number of replicates (N).

S9 Table. Experiment 1. Means (SD) of arcsin transformed percent Survival.

S10 Table. Experiment 1. Means (SD) Prime female mass at pupation (mg).

S11 Table. Experiment 1. Means (SD) Prime female age at pupation (days).

S12 Table. Experiment 1. Means (SD) Average female mass at pupation (mg).

S13 Table. Experiment 1. Means (SD) Prime male mass at pupation (mg).

S14 Table. Experiment 1. Means (SD) Prime male age at pupation (days).

S15 Table. Experiment 1. Means (SD) Average male mass at pupation (mg).

S16 Table. Experiment 1. Means (SE) for FxDxT for arcsin transformed percent Survival.

S17 Table. Experiment 1. Means (SE) for FxDxT for Prime female mass and age, and Average female mass. Estimated growth rate and difference between the Prime and Average female mass.

S18 Table. Experiment 1. Means (SE) for Prime female mass and age at pupation and Average female mass at pupation for the interaction FxDxT. Total food after day 4 and food/larva after day 4.

S19 Table. Experiment 1. Means (SE) for FxDxT for Prime male mass and age, and Average male mass.

S20 Table. Experiment 1. Means (SD) Average female mass for FxDxAxT.

S21 Table. Experiment 1. Means (SE) for FxDxA for Prime female mass and age, and Average female mass. Estimated growth rate and difference between the Prime and Average female mass.

S22 Table. Experiment 1. Means (SE) for FxDxA for Prime female mass and age, and Average female mass. Total food and food/larva after day 4.

S23 Table. Experiment 1. Means (SE) for FxD for Prime female mass and age, and Average female mass. Expected mean values for Prime female age and Average female mass.

S24 Table. Experiment 1. Means (SE) for FxD for Prime male mass and age, and Average male mass. Expected mean values for Prime male age and Average male mass.

S25 Table. Experiment 1. Means (SE) for FxAxT for Prime female mass and age, and Average female mass. Estimated growth rate and difference between the Prime and Average female mass.

S26 Table. Experiment 1. Means (SE) for FxAxT for Prime female mass and age, and Average female mass. Total food and food/larva after day 4.

S27 Table. Experiment 1. Means (SE) for FxA for Prime male mass and age, and Average male mass. Expected mean values for the Prime male mass and age and the Average male mass.

S28 Table. Experiment 1. Means (SE) for Prime female mass and Average female mass at pupation for the interaction FxA. Differences between the Prime female mass and the Prime male mass and between the Average female mass and the Average male mass.

S29 Table. Experiment 1. Means (SE) for arcsin transformed percent Survival for the interaction FxT.

S30 Table. Experiment 1. Means (SE) for Prime female mass and age and Average female mass for the interaction FxT. Total food, expected values, growth rates and the differences between Prime and Average female masses.

S31 Table. Experiment 1. Means (SE) for Prime male mass and age and Average male mass for the interaction FxT. Expected values, growth rates and the differences between Prime and Average female masses.

S32 Table. Experiment 1. Means (SE) for Prime male mass and age and Average male mass for the interaction FxT. Differences between the Prime female mass and Prime male mass and between the Average female mass and the Average male mass.

S33 Table. Experiment 1. Means (SE) for Prime female mass and age at pupation and Average female mass at pupation for the interaction AxT. Expected values, growth rates and the differences between the Prime female mass and the Average female mass.

S34 Table. Experiment 1. Means (SE) for Prime male mass and age at pupation and Average male mass at pupation for the interaction AxT. Expected values, growth rates and the differences between the Prime male mass and the Average male mass.

S35 Table. Experiment 1. Means (SE) for Prime female mass for the interaction AxT. Differences between Prime female mass and Average female mass, Prime male mass and Average male mass, Prime female mass and Prime male mass, and Average female mass and Average male mass. Total food after day 4 and food/larva after day 4.

S36 Table. Experiment 1. Means (SE) for Prime female mass and age at pupation and Average female mass at pupation for the interaction DxAxT. Estimated growth rates and the differences between the Prime female mass and the Average female mass. Total food after day 4 and food/larva after day 4.

S37 Table. Experiment 1. Means (SE) for Prime male mass and age at pupation and Average male mass at pupation for the interaction DxAxT. Estimated growth rates and the differences between the Prime male mass and the Average male mass.

S38 Table. Experiment 1. Means (SE) for Prime female mass and Average male mass for the interaction DxAxT. Prime female mass MINUS Average female mass, Prime female mass MINUS Prime male mass, Prime male mass MINUS Average male mass, Average female mass MINUS Average male mass.

S39 Table. Experiment 1. Means (SE) for Prime female mass and age at pupation and Average female mass at pupation for the interaction DxA. Estimated growth rates, differences between Prime and Average female masses, expected mean values for Prime female age, and food levels after day 4 and on day 0.

S40 Table. Experiment 1. Means (SE) for arcsin transformed percent Survival for the interaction DxT. Expected values.

S41 Table. Experiment 1. Means (SE) for Prime female mass and age at pupation and Average female mass at pupation for the interaction DxT. Expected mean values, growth rates, differences between Prime and Average female masses, food levels.

S42 Table. Experiment 1. Means (SE) for Prime male mass and age at pupation and Average male mass at pupation for the interaction DxT. Expected mean values, growth rates, differences between Prime and Average male masses.

S43 Table. Experiment 1. Means (SE) for Prime female mass at pupation for the interaction DxT. Differences between Prime and Average female masses, Prime and Average male masses, Prime female and Prime male masses, and Average female and male masses. Food/larva.

S44 Table. Experiment 3. Means (SE) for mass and age at pupation for main effects: food input amount, day of second food input, and sex.

S45 Table. Experiment 3. Means (SE) for mass (mg) for the interaction food 2 x delay x sex.

S46 Table. Experiment 3. Means (SE) for age (days) for the interaction food 2 x delay x sex.

S47 Table. Experiment 3. Means (SE) for estimated growth rates (mg/day) for the interaction food 2 x delay x sex.

S48 Table. Experiment 3. Means (SE) for mass (mg) for the interaction food 1 x delay x sex.

S49 Table. Experiment 3. Means (SE) for age (days) for the interaction food 1 x delay x sex.

S50 Table. Experiment 3. Means (SE) for estimated growth rates (mg/day) for the interaction food 1 x delay x sex.

S51 Table. Experiment 3. Means (SE), expected values and differences for mass (mg) for the interaction food 1 x delay.

S52 Table. Experiment 3. Means (SE), expected values and differences for age (days) for the interaction food 1 x delay.

S53 Table. Experiment 3. Means (SE), expected values and differences for mass (mg) for the interaction food 2 x delay.

S54 Table. Experiment 3. Means (SE) for age (days) for the interaction food 2 x delay (not significant in the ANOVA).

S55 Table. Experiment 3. Means (SE), expected values and differences for age (days) for the interaction food 1 x sex.

S56 Table. Experiment 3. Means (SE) for mass (mg) for the interaction food 1 x sex (not significant in the ANOVA).

S57 Table. Experiment 3. Means (SE) for estimated growth rates (mg/day) for the interaction food 1 x sex.

S58 Table. Experiment 3. Means (SE), expected values and differences for age (days) for the interaction food 2 x sex.

S59 Table. Experiment 3. Means (SE) for mass (mg) for the interaction food 2 x sex (not significant in the ANOVA).

S60 Table. Experiment 3. Means (SE) for estimated growth rates (mg/day) for the interaction food 2 x sex.

S1 Text. Experiment 1. First experiment results analysis by interaction.

S2 Text. Experiment 1. First experiment results summary by interaction.

S3 Text. Experiment 3. Third experiment analysis of individual ANOVA contrasts.

## References

1. Steinwascher K. Competition among Aedes aegypti larvae. PLoS One. 2018. doi:10.1371/journal.pone.0202455

2. Bedhomme S, Agnew P, Sidobre C, Michalakis Y. Sex-specific reaction norms to intraspecific larval competition in the mosquito Aedes aegypti. J Evol Biol. 2003;16: 721–730. doi:10.1046/j.1420-9101.2003.00576.x

3. Daugherty MP, Alto BW, Juliano S a. Invertebrate carcasses as a resource for competing Aedes albopictus and Aedes aegypti (Diptera: Culicidae). J Med Entomol. 2000;37: 364–372. doi:10.1603/0022-2585(2000)037[0364:ICAARF]2.0.CO;2

4. Telang A, Frame L, Brown MR. Larval feeding duration affects ecdysteroid levels and nutritional reserves regulating pupal commitment in the yellow fever mosquito Aedes aegypti (Diptera: Culicidae). J Exp Biol. 2007;210: 854–864. doi:10.1242/jeb.02715

5. Saul SH, Novak RJ, Ross QE. The Role of the Preadult Stages in the Ecological Separation of Two Subspecies of Aedes aegypti. Am Midl Nat. 1980;104: 118–134.

6. Murrell EG, Juliano SA. Detritus Type Alters the Outcome of Interspecific Competition Between *Aedes aegypti* and *Aedes albopictus* (Diptera: Culicidae). J Med Entomol. 2008. doi:10.1603/0022-2585(2008)45[375:dtatoo]2.0.co;2

7. Murrell EG, Damal K, Lounibos LP, Juliano SA. Distributions of Competing Container Mosquitoes Depend on Detritus Types, Nutrient Ratios, and Food Availability. Ann Entomol Soc Am. 2011. doi:10.1603/an10158

8. Moore CG, Fisher BR. Competition in Mosquitoes. Density and Species Ratio Effects on Growth, Mortality, Fecundity and Production of Growth Retardant. Ann Entomol Soc Am. 1969;62: 1325– 1331.

9. Steinwascher K. Competitive Interactions among Tadpoles : Responses to Resource Level. Ecology. 1979;60: 1172–1183. doi:10.2307/1936965

10. Padmanabha H, Bolker B, Lord CC, Lounibos LP, Rubio C, Lounibos LP. Food Availability Alters the Effects of Larval Temperature on Aedes aegypti Growth. J Med Entomol. 2011;48: 974–984. doi:10.3174/ajnr.A1256.Functional

11. Juliano SA. Species Introduction and Replacement among Mosquitoes : Interspecific Resource Competition or Apparent Competition? Ecology. 1998;79: 255–268. doi:10.1890/0012-9658(1998)079[0255:SIARAM]2.0.CO;2

12. Riback TIS, Honório NA, Pereira RN, Godoy WAC, Codeço CT. Better to be in bad company than to be alone? Aedes vectors respond differently to breeding site quality in the presence of others. PLoS One. 2015;10: 1–16. doi:10.1371/journal.pone.0134450

13. Lounibos LP, Suárez S, Menéndez Z, Nishimura N, Escher RL, O’Connell SM, et al. Does temperature affect the outcome of larval competition between Aedes aegypti and Aedes albopictus? J Vector Ecol. 2002;27: 86–95. Available: http://www.sove.org/Society_for_Vector_Ecology/Journal/Entries/2002/6/1_Volume_27,_Number_1_files/Lounibosetal.pdf

14. Wada Y. Effects of larval density on the development of Aedes aegypti (L.) and the size of adults. Quaest Entomol. 1965;1: 223–249.

15. Wilbur HM. Interactions of Food Level and Population Density in Rana Sylvatica. Ecology. 1977;58: 206–209.

16. Steinwascher K. Interference and Exploitation Competition Among Tadpoles of Rana Utricularia. Ecology. 1978;59: 1039–1046. doi:10.2307/1938556

17. Gilpin ME, McClelland GAH. Systems Analysis of the Yellow Fever Mosquito Aedes aegypti. Fortschritte Zool. 1979;25: 355–388.

18. Rubenstein DI. Individual Variation and Competition in the Everglades Pygmy Sunfish. J Anim Ecol. 1981;50: 337–350.

19. Agnew P, Hide M, Sidobre C, Michalakis Y. A minimalist approach to the effects of density-dependent competition on insect life-history traits. Ecol Entomol. 2002;27: 396–402. doi:10.1046/j.1365-2311.2002.00430.x

20. Antonaci Gama R, De Carvalho Alves K, Ferreira Martins R, Eiras ÁE, Carvalho De Resende M. Efeito da densidade larval no tamanho de adultos de Aedes aegypti criados em condições de laboratório. Rev Soc Bras Med Trop. 2005;38: 64–66. doi:10.1590/S0037-86822005000100014

21. Couret J, Dotson E, Benedict MQ. Temperature, larval diet, and density effects on development rate and survival of Aedes aegypti (Diptera: Culicidae). PLoS One. 2014. doi:10.1371/journal.pone.0087468

22. Steinwascher K. Host-parasite interaction as a potential population-regulating mechanism. Ecology. 1979;60: 884–890. doi:10.2307/1936856

23. Sunahara T, Mogi M. Priority effects of bamboo-stump mosquito larvae: Influences of water exchange and leaf litter input. Ecol Entomol. 2002;27: 346–354. doi:10.1046/j.1365-2311.2002.00417.x

24. O’Neal PA, Juliano SA. Seasonal variation in competition and coexistence of Aedes mosquitoes: Stabilizing effects of egg mortality or equalizing effects of resources? J Anim Ecol. 2013;82: 256– 265. doi:10.1111/j.1365-2656.2012.02017.x

25. Walker ED, Lawson DL, Merritt RW, Morgan WT, Lawson DL, Klug MJ. Nutrient Dynamics, Bacterial Populations, and Mosquito Productivity in Tree Hole Ecosystems and Microcosms. Ecology. 2016;72: 1529–1546. doi:10.2307/1940953

26. Bevins SN. Timing of resource input and larval competition between invasive and native container-inhabiting mosquitoes (Diptera: Culicidae). J Vector Ecol. 2007. doi:10.3376/1081-1710(2007)32[252:TORIAL]2.0.CO;2

27. Bevins SN. Invasive mosquitoes, larval competition, and indirect effects on the vector competence of native mosquito species (Diptera: Culicidae). Biol Invasions. 2008;10: 1109–1117. doi:10.1007/s10530-007-9188-8

28. Wong J, Stoddard ST, Astete H, Morrison AC, Scott TW. Oviposition site selection by the dengue vector Aedes aegypti and its implications for dengue control. PLoS Negl Trop Dis. 2011;5. doi:10.1371/journal.pntd.0001015

29. Ikeshoji T, Mulla MS. Overcrowding factors of mosquito larvae. J Econ Entomol. 1970;63: 90–96.

30. Ikeshoji TM, Mulla MS. Overcrowding factors of mosquito larvae. 2. Growth-retarding and bacteriostatic effects of the overcrowding factors of mosquito larvae. J Econ Entomol. 1970;63: 1737–1743.

31. Moore CG, Whitacre DM. Competition in Mosquitoes. 2. Production of Aedes aegypti Larval Growth Retardant at Various Densities and Nutrition Levels. Ann Entomol Soc Am. 1972;65: 915– 918. doi:10.1093/aesa/65.4.915

32. Dye C. Competition amongst larval Aedes aegypti: the role of interference. Ecol Entomol. 1984;9: 355–357. doi:10.1111/j.1365-2311.1984.tb00859.x

33. Bédhomme S, Agnew P, Sidobre C, Michalakis Y. Pollution by conspecifics as a component of intraspecific competition among Aedes aegypti larvae. Ecol Entomol. 2005;30: 1–7. doi:10.1111/j.0307-6946.2005.00665.x

34. Briegel H. Metabolic relationship between female body size, reserves, and fecundity of Aedes aegypti. J Insect Physiol. 1990;36: 165–172. doi:10.1016/0022-1910(90)90118-Y

35. Helinski MEH, Harrington LC. Male Mating History and Body Size Influence Female Fecundity and Longevity of the Dengue Vector Aedes aegypti. J Med Entomol. 2011;48: 202–211. doi:10.1603/ME10071

36. Ponlawat A, Harrington LC. Age and body size influence male sperm capacity of the dengue vector Aedes aegypti (Diptera: Culicidae). J Med Entomol. 2007;44: 422–426. doi:10.1603/0022-2585(2007)44[422:AABSIM]2.0.CO;2

37. Timm NH. Multivariate Analysis. Monterey, California: Brooks/Cole Publishing Company; 1975.

38. Morrison DF. Multivariate Statistical Methods. New York: McGraw-Hill Book Company; 1967.

39. Aznar VR, Alem I, Majo MS De, Byttebier B, Solari HG, Fischer S. Effects of scarcity and excess of larval food on life history traits of Aedes aegypti (Diptera : Culicidae). J Vector Ecol. 2018;43: 117–124. doi:10.1111/jvec.12291

40. Reiskind MH, Lounibos LP. Effects of intraspecific larval competition on adult longevity in the mosquitoes Aedes aegypti and Aedes albopictus. Med Vet Entomol. 2009;23: 62–68. doi:10.1111/j.1365-2915.2008.00782.x

41. Leisnham PT, Juliano SA. Interpopulation differences in competitive effect and response of the mosquito Aedes aegypti and resistance to invasion by a superior competitor. Oecologia. 2010;164: 221–230. doi:10.1007/s00442-010-1624-2

42. Bara JJ, Clark TM, Remold SK. Utilization of larval and pupal detritus by Aedes aegypti and Aedes albopictus. J Vector Ecol. 2014;39: 44–47. doi:10.1111/j.1948-7134.2014.12068.x

43. Yee DA, Kesavaraju B, Juliano SA. Larval feeding behavior of three co-occurring species of container mosquitoes. J vector Ecol. 2004;29: 315–322.

44. Rey JR, Lounibos P. Ecology of aedes aegypti and aedes albopictus in the Americas and disease transmission. Biomedica. 2015. doi:10.7705/biomedica.v35i2.2514

45. Farjana T, Tuno N, Higa Y. Effects of temperature and diet on development and interspecies competition in Aedes aegypti and Aedes albopictus. Med Vet Entomol. 2012. doi:10.1111/j.1365-2915.2011.00971.x

46. Black WC, Rai KS, Turco BJ, Arroyo DC. Laboratory Study of Competition Between United States Strains of Aedes albopictus and Aedes aegypti (Diptera: Culicidae). J Med Entomol. 1989. doi:10.1093/jmedent/26.4.260

47. Leisnham PT, LaDeau SL, Juliano SA. Spatial and temporal habitat segregation of mosquitoes in Urban Florida. PLoS One. 2014;9. doi:10.1371/journal.pone.0091655

48. Noden BH, O’Neal PA, Fader JE, Juliano SA. Impact of inter- and intra-specific competition among larvae on larval, adult, and life-table traits of Aedes aegypti and Aedes albopictus females. Ecol Entomol. 2016;41: 192–200. doi:10.1111/een.12290

49. Juliano SA, Lounibos LP, O’Meara GF. A field test for competitive effects of Aedes albopictus on A. aegypti in South Florida: Differences between sites of coexistence and exclusion? Oecologia. 2004;139: 583–593. doi:10.1007/s00442-004-1532-4

50. Oliveira S De, Antunes D, Villela M, Braga F, Dias S. How does competition among wild type mosquitoes influence the performance of Aedes aegypti and dissemination of Wolbachia pipientis ? PLoS One. 2017; 1–20.

51. Camara DCP, Codeço CT, Juliano SA, Lounibos LP, Riback TIS, Pereira GR, et al. Seasonal differences in density but similar competitive impact of aedes albopictus (Skuse) on aedes aegypti (L.) in rio de janeiro, Brazil. PLoS One. 2016;11: 1–15. doi:10.1371/journal.pone.0157120

52. Lounibos LP, Bargielowski I, Carrasquilla MC, Nishimura N. Coexistence of Aedes aegypti and Aedes albopictus (Diptera: Culicidae) in Peninsular Florida Two Decades After Competitive Displacements. J Med Entomol. 2016;53: 1385–1390. doi:10.1093/jme/tjw122

53. Braks MAH, Honório NA, Lounibos LP, Lourenço-De-Oliveira R, Juliano SA. Interspecific Competition Between Two Invasive Species of Container Mosquitoes, *Aedes aegypti* and *Aedes albopictus* (Diptera: Culicidae), in Brazil. Ann Entomol Soc Am. 2006. doi:10.1603/0013-8746(2004)097[0130:icbtis]2.0.co;2

54. Ho BC, Ewert A, Chew LM. Interspecific competition among Aedes aegypti, Ae. albopictus, and Ae. triseriatus (Diptera: Culicidae): larval development in mixed cultures. J Med Entomol. 1989;26: 615–623. doi:10.1093/jmedent/26.6.615

55. Leisnham PT, Lounibos LP, O’Meara GF, Juliano SA. Interpopulation divergence in competitive interactions of the mosquito Aedes albopictus. Ecology. 2009. doi:10.1890/08-1569.1

56. Kaplan L, Kendell D, Robertson D, Livdahl T, Khatchikian C. Aedes aegypti and Aedes albopictus in Bermuda: Extinction, invasion, invasion and extinction. Biol Invasions. 2010. doi:10.1007/s10530-010-9721-z

57. Bargielowski IE, Lounibos LP, Shin D, Smartt CT, Carrasquilla MC, Henry A, et al. Widespread evidence for interspecific mating between Aedes aegypti and Aedes albopictus (Diptera: Culicidae) in nature. Infect Genet Evol. 2015. doi:10.1016/j.meegid.2015.08.016

58. Muturi EJ, Kim CH, Alto BW, Berenbaum MR, Schuler MA. Larval environmental stress alters Aedes aegypti competence for Sindbis virus. Trop Med Int Heal. 2011;16: 955–964. doi:10.1111/j.1365-3156.2011.02796.x

59. Serpa LLN, Kakitani I, Voltolini JC. Competição entre larvas de Aedes aegypti e Aedes albopictus em laboratório. Rev Soc Bras Med Trop. 2008;41: 479–484. doi:10.1590/S0037-86822008000500009

60. Barrera R. Competition and resistance to starvation in larvae of container-inhabiting Aedes mosquitoes. Ecol Entomol. 1996;21: 117–127. doi:10.1111/j.1365-2311.1996.tb01178.x

61. Alto BW, Bettinardi DJ, Ortiz S. Interspecific larval competition differentially impacts adult survival in dengue vectors. J Med Entomol. 2015;52: 163–170. doi:10.1093/jme/tju062

62. Costanzo KS, Kesavaraju B, Juliano SA. Condition-specific competition in container mosquitoes: The role of noncompeting life-history stages. Ecology. 2005. doi:10.1890/05-0583

63. Kokkinn MJ, Roberts DM, Williams CR. Larval development rate of the mosquitoes Culex quinquefasciatus and Aedes aegypti (Diptera: Culicidae) varies between clutches: Implications for population ecology. Aust J Entomol. 2012;51: 22–27. doi:10.1111/j.1440-6055.2011.00837.x

64. Wong J, Morrison AC, Stoddard ST, Astete H, Chu YY, Baseer I, et al. Linking oviposition site choice to offspring fitness in Aedes aegypti: Consequences for targeted larval control of dengue vectors. PLoS Negl Trop Dis. 2012;6: 1–12. doi:10.1371/journal.pntd.0001632

65. Harrington LC, Ponlawat A, Edman JD, Scott TW, Vermeylen F. Influence of container size, location, and time of day on oviposition patterns of the dengue vector, Aedes aegypti, in Thailand. Vector-Borne Zoonotic Dis. 2008. doi:10.1089/vbz.2007.0203

66. Chadee DD. Oviposition strategies adopted by gravid Aedes aegypti (L.) (Diptera: Culicidae) as detected by ovitraps in Trinidad, West Indies (2002-2006). Acta Trop. 2009. doi:10.1016/j.actatropica.2009.05.012

67. Guagliardo SA, Barboza JL, Morrison AC, Astete H, Vazquez-Prokopec G, Kitron U. Patterns of Geographic Expansion of Aedes aegypti in the Peruvian Amazon. PLoS Negl Trop Dis. 2014. doi:10.1371/journal.pntd.0003033

68. Ponnusamy L, Xu N, Nojima S, Wesson DM, Schal C, Apperson CS. Identification of bacteria and bacteria-associated chemical cues that mediate oviposition site preferences by Aedes aegypti. Proc Natl Acad Sci U S A. 2008. doi:10.1073/pnas.0802505105

69. Ponnusamy L, Xu N, Böröczky K, Wesson DM, Ayyash LA, Schal C, et al. Oviposition responses of the mosquitoes aedes aegypti and Aedes albopictus to experimental plant infusions in laboratory bioassays. J Chem Ecol. 2010. doi:10.1007/s10886-010-9806-2

70. Fader JE, Juliano SA. Oviposition habitat selection by container-dwelling mosquitoes : responses to cues of larval and detritus abundances in the field. Ecol Entomol. 2014;39: 245–252. doi:10.1111/een.12095

71. Steinwascher K. Relationship Between Pupal Mass and Adult Survivorship and Fecundity for Aedes aegypti. Environ Entomol. 1982;11: 150–153. doi:10.1093/ee/11.1.150

72. Steinwascher K. Egg Size Variation in Aedes aegypti: Relationship to Body Size and Other Variables. Am Midl Nat. 1984;112: 76–84. doi:10.2307/2425459

73. Muturi EJ, Orindi BO, Kim CH. Effect of Leaf Type and Pesticide Exposure on Abundance of Bacterial Taxa in Mosquito Larval Habitats. PLoS One. 2013;8. doi:10.1371/journal.pone.0071812

74. Ahmad R, Chu W-LW-L, Lee H-LH-L, Phang S-MS-M. Effect of four chlorophytes on larval survival, development and adult body size of the mosquito Aedes aegypti. J Appl Phycol. 2001;13: 369– 374. doi:10.1023/A:1017966802600

75. Ahmad R, Chu WL, Ismail Z, Lee HL, Phang SM. Effect of ten chlorophytes on larval survival, development and adult body size of the mosquito Aedes aegypti. Southeast Asian J Trop Med Public Heal. 2004;35: 79–87. doi:10.1023/A:1017966802600

76. Lounibos LP, Nishimura N, Escher RL. Fitness of a treehole mosquito: influences of food type and predation. Oikos. 1993;66: 114–118. doi:10.2307/3545203

77. Muturi EJ, Allan BF, Ricci J. Influence of Leaf Detritus Type on Production and Longevity of Container-Breeding Mosquitoes. Environ Entomol. 2012;41: 1062–1068. doi:10.1603/EN11301

78. Walker ED, Merritt RW, Kaufman MG, Ayres MP, Riedel MH. Effects of variation in quality of leaf detritus on growth of the eastern tree-hole mosquito, Aedes triseriatus (Diptera: Culicidae). Can J Zool. 1997;75: 706–718. doi:10.1139/z97-091

79. Pelz-Stelinski KS, Walker ED, Kaufman MG. Senescent leaf exudate increases mosquito survival and microbial activity. Ecol Entomol. 2010;35: 329–340. doi:10.1111/j.1365-2311.2010.01183.x

80. Muturi EJ, Gardner AM, Bara JJ. Impact of an Alien Invasive Shrub on Ecology of Native and Alien Invasive Mosquito Species (Diptera: Culicidae). Environ Entomol. 2015;44: 1308–1315. doi:10.1093/ee/nvv121

81. Kim C-H, Muturi EJ. Relationship between leaf litter identity, expression of cytochrome P450 genes and life history traits of Aedes aegypti and Aedes albopictus. Acta Trop. 2012;122: 94–100. doi:10.1016/j.actatropica.2011.12.006

82. Liedo P, Dor A. Efficiency of two larval diets for mass-rearing of the mosquito Aedes aegypti. PLoS One. 2017; 1–13.

83. Bond JG, Ramírez-Osorio A, Marina CF, Fernández-Salas I, Liedo P, Dor A, et al. Efficiency of two larval diets for mass-rearing of the mosquito Aedes aegypti. PLoS One. 2017;12. doi:10.1371/journal.pone.0187420

84. Luxananil P, Atomi H, Panyim S, Imanaka T. Isolation of bacterial strains colonizable in mosquito larval guts as novel host cells for mosquito control. J Biosci Bioeng. 2001;92: 342–345. doi:10.1016/S1389-1723(01)80237-1

85. Gaio ADO, Gusmão DS, Santos A V., Berbert-Molina MA, Pimenta PFP, Lemos FJA. Contribution of midgut bacteria to blood digestion and egg production in aedes aegypti (diptera: Culicidae) (L.). Parasites and Vectors. 2011. doi:10.1186/1756-3305-4-105

86. Terenius O, Lindh JM, Eriksson-Gonzales K, Bussière L, Laugen AT, Bergquist H, et al. Midgut bacterial dynamics in Aedes aegypti. FEMS Microbiol Ecol. 2012. doi:10.1111/j.1574-6941.2012.01317.x

87. Coon KL, Brown MR, Strand MR. Gut bacteria differentially affect egg production in the anautogenous mosquito Aedes aegypti and facultatively autogenous mosquito Aedes atropalpus (Diptera: Culicidae). Parasites and Vectors. 2016. doi:10.1186/s13071-016-1660-9

88. Caragata EP, Rancès E, O’Neill SL, McGraw EA. Competition for Amino Acids Between Wolbachia and the Mosquito Host, Aedes aegypti. Microb Ecol. 2014. doi:10.1007/s00248-013-0339-4

89. Tripet F, Lounibos LP, Robbins D, Moran J, Nishimura N, Blosser EM. Competitive reduction by satyrization? Evidence for interspecific mating in nature and asymmetric reproductive competition between invasive mosquito vectors. Am J Trop Med Hyg. 2011. doi:10.4269/ajtmh.2011.10-0677

90. Bargielowski IE, Lounibos LP. Rapid evolution of reduced receptivity to interspecific mating in the dengue vector Aedes aegypti in response to satyrization by invasive Aedes albopictus. Evol Ecol. 2014;28: 193–203. doi:10.1007/s10682-013-9669-4

91. Bargielowski IE, Lounibos LP. Satyrization and satyrization-resistance in competitive displacements of invasive mosquito species. Insect Sci. 2016;23: 162–174. doi:10.1111/1744-7917.12291

92. De Jesus CE, Reiskind MH. The importance of male body size on sperm uptake and usage, and female fecundity in Aedes aegypti and Aedes albopictus. Parasites and Vectors. 2016;9: 1–7. doi:10.1186/s13071-016-1734-8

93. Duvall LB, Basrur NS, Molina H, McMeniman CJ, Vosshall LB. A Peptide Signaling System that Rapidly Enforces Paternity in the Aedes aegypti Mosquito. Curr Biol. 2017. doi:10.1016/j.cub.2017.10.074

94. Chambers GM, Klowden MJ. Correlation of nutritional reserves with a critical weight for pupation in larval Aedes aegypti mosquitoes. J Am Mosq Control Assoc. 1990;6: 394–399.

95. Briegel H. Physiological bases of mosquito ecology. J Vector Ecol. 2003;28: 1–11. Available: http://www.ncbi.nlm.nih.gov/pubmed/12831123

96. Maciel-De-Freitas R, Codeço CT, Lourenço-De-Oliveira R. Body size-associated survival and dispersal rates of Aedes aegypti in Rio de Janeiro. Med Vet Entomol. 2007;21: 284–292. doi:10.1111/j.1365-2915.2007.00694.x

97. Nasci RS. The size of emerging and host-seeking Aedes aegypti and the relation of size to blood-feeding success in the field. J Am Mosq Control Assoc. 1986;2: 61–62. doi:10.1101/gr.3715005

98. Briegel H, Knuesel I, Timmerman SE, Knüsel I, Timmermann SE. Aedes aegypti size reserves survival and flight potential. J Vector Ecol. 2001;26: 21–31.

99. Maciá A. Differences in performance of Aedes aegypti larvae raised at different densities in tires and ovitraps under field conditions in Argentina. J Vector Ecol. 2006;31: 371–377. doi:10.3376/1081-1710(2006)31[371:dipoaa]2.0.co;2

100. Telang A. Effects of larval nutrition on the endocrinology of mosquito egg development. J Exp Biol. 2006;209: 645–655. doi:10.1242/jeb.02026

101. Koenraadt CJM. Pupal dimensions as predictors of adult size in fitness studies of Aedes aegypti (Diptera: Culicidae). J Med Entomol. 2008;45: 331–336. doi:10.1603/0022-2585(2008)45[331:pdapoa]2.0.co;2

102. Mitchell-Foster K, Ma BO, Warsame-Ali S, Logan C, Rau ME, Lowenberger C. The influence of larval density, food stress, and parasitism on the bionomics of the dengue vector Aedes aegypti (Diptera: Culicidae): implications for integrated vector management. J Vector Ecol. 2012;37: 221– 229. doi:10.1111/j.1948-7134.2012.00220.x

103. Yeap HL, Endersby NM, Johnson PH, Ritchie SA, Hoffmann AA. Body size and wing shape measurements as quality indicators of aedes aegypti mosquitoes destined for field release. Am J Trop Med Hyg. 2013;89: 78–92. doi:10.4269/ajtmh.12-0719

104. Timmermann SE, Briegel H. Larval growth and biosynthesis of reserves in mosquitoes. J Insect Physiol. 1999;45: 461–470. doi:10.1016/S0022-1910(98)00147-4

105. Nishiura JT, Burgos C, Aya S, Goryacheva Y, Lo W. Modulation of larval nutrition affects midgut neutral lipid storage and temporal pattern of transcription factor expression during mosquito metamorphosis. J Insect Physiol. 2007;53: 47–58. doi:10.1016/j.jinsphys.2006.09.014

106. Timmermann SE, Briegel H. Water depth and larval density affect development and accumulation of reserves in laboratory populations of mosquitoes. Bull Soc Vector Ecol. 1993;18: 174–187.

107. Telang A, Peterson B, Frame L, Baker E, Brown MR. Analysis of molecular markers for metamorphic competency and their response to starvation or feeding in the mosquito, Aedes aegypti (Diptera: Culicidae). J Insect Physiol. 2010;56: 1925–1934. doi:10.1016/j.jinsphys.2010.08.020

108. Padmanabha H, Lord CC, Lounibos LP. Temperature induces trade-offs between development and starvation resistance in Aedes aegypti (L.) larvae. Med Vet Entomol. 2011;25: 445–453. doi:10.1111/j.1365-2915.2011.00950.x

109. Padmanabha H, Correa F, Legros M, Nijhout HF, Lord C, Lounibos LP. An eco-physiological model of the impact of temperature on Aedes aegypti life history traits. J Insect Physiol. 2012;58: 1597–1608. doi:10.1016/j.jinsphys.2012.09.015

